# A limited set of transcriptional programs define major cell types

**DOI:** 10.1101/857169

**Authors:** Alessandra Breschi, Manuel Muñoz-Aguirre, Valentin Wucher, Carrie A. Davis, Diego Garrido-Martín, Sarah Djebali, Jesse Gillis, Dmitri D. Pervouchine, Anna Vlasova, Alexander Dobin, Chris Zaleski, Jorg Drenkow, Cassidy Danyko, Alexandra Scavelli, Ferran Reverter, Michael P. Snyder, Thomas R. Gingeras, Roderic Guigó

## Abstract

We have produced RNA sequencing data for a number of primary cells from different locations in the human body. The clustering of these primary cells reveals that most cells in the human body share a few broad transcriptional programs, which define five major cell types: epithelial, endothelial, mesenchymal, neural and blood cells. These act as basic components of many tissues and organs. Based on gene expression, these cell types redefine the basic histological types by which tissues have been traditionally classified. We identified genes whose expression is specific to these cell types, and from these genes, we estimated the contribution of the major cell types to the composition of human tissues. We found this cellular composition to be a characteristic signature of tissues, and to reflect tissue morphological heterogeneity and histology. We identified changes in cellular composition in different tissues associated with age and sex and found that departures from the normal cellular composition correlate with histological phenotypes associated to disease.

**One Sentence Summary:** A few broad transcriptional programs define the major cell types underlying the histology of human tissues and organs.

## Main Text

Transcriptional profiles reflect cell type, condition and function. In tissues and organs, they are monitored in RNA extracted from millions to billions of cells (11^6^-10^9^) *(1)* likely including multiple cell types. As a consequence, the transcriptional profiles obtained from tissue samples represent the average expression of genes across heterogeneous cellular collections, and gene expression differences measured in bulk tissue transcriptomes may thus reflect changes in cellular composition rather than changes in the expression of genes in individual cells. Single-cell RNA sequencing (scRNA-seq) has indeed revealed large cellular heterogeneity in many tissues and organs *(2)*, and the Human Cell Atlas (HCA) project *(3)* has been recently initiated with the aim of defining all human cell types and to infer the cellular taxonomy of the human body. As a step in that direction and to bridge the transcriptomes of tissues with the transcriptomes of the constituent primary cells, and to understand how these impact tissue phenotypes, we have generated bulk expression profiles of 53 primary cell lines isolated from ten different anatomical sites in the human body. These profiles include long and short strand-specific RNA-seq, and RAMPAGE data (Fig. 1a, Table S1-4).

**Fig. 1.**
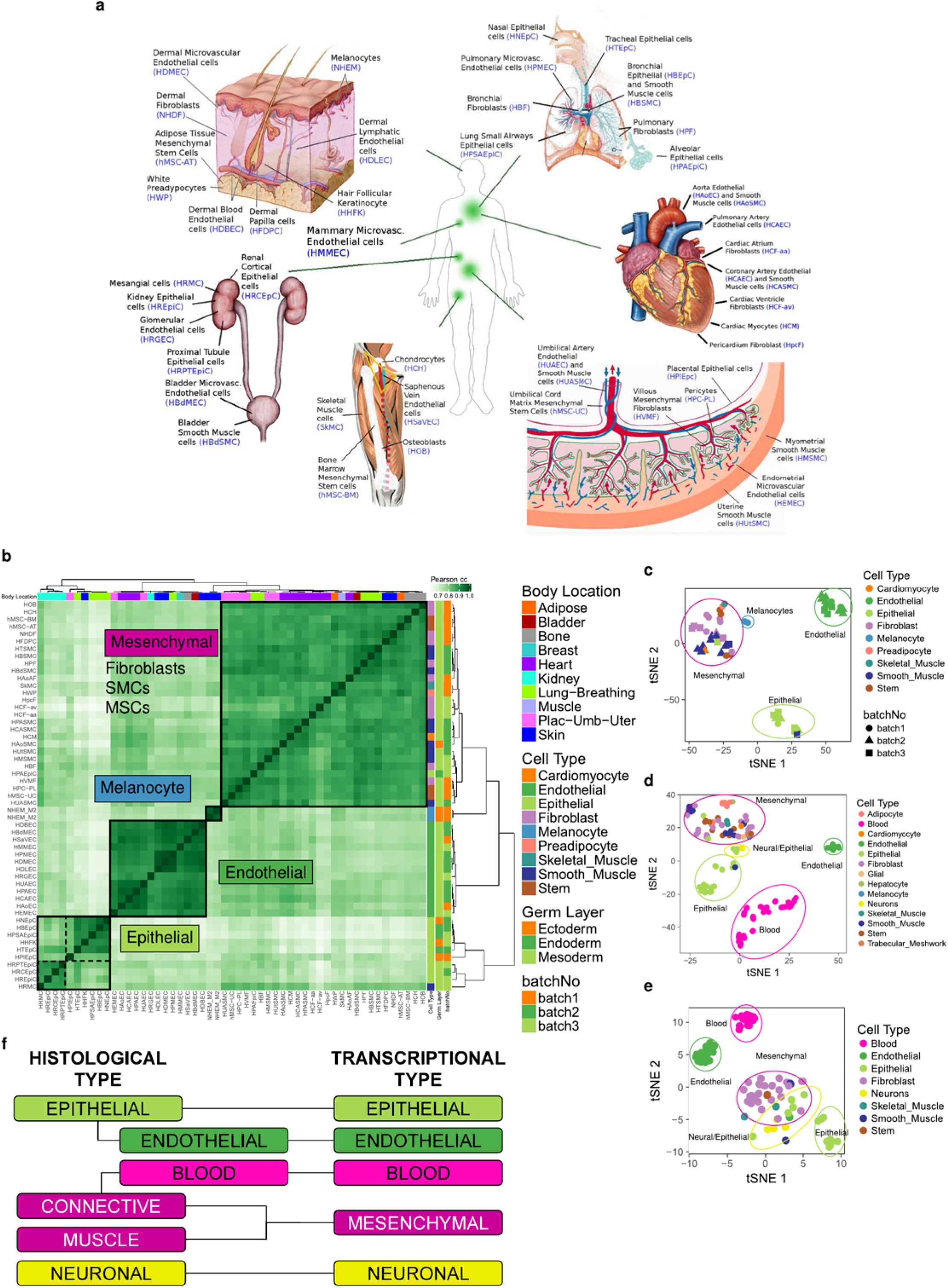
Basic transcriptional programs of human primary cells. **(a)** Overview of primary cells analyzed in this study and the body location they are extracted from. **(b)** Hierarchical clustering of human primary cells based on the correlation of gene expression. The clustering in four major clusters is supported by the silhouette analysis and the elbow method (Fig. S2a-b). tSNE of human primary cells based on gene expression measured here **(c)**, on gene expression measured by CAGE by the FANTOM consortium **(d)** and on Candidate Regulatory Elements (cREs) by the ENCODE encyclopedia scored DNAse hypersensitivity signal **(e)**. **(f)** Correspondence between transcriptionally derived major cell types and classical histological types.

### Major cell types in the human body

Clustering of the primary cells based on gene expression profiles revealed a number of well-defined clusters (Fig. 1b-c, Fig. S1, S2a-b, Supplementary Information). One cluster was composed of endothelial cells, a second large cluster included a mixture of cell types: fibroblasts, stem cells and muscle cells, among others, which we collectively termed as mesenchymal, two smaller clusters, which clustered together, were composed of epithelial cells, and finally, the melanocytes clustered separately. Almost all of the individual primary cells are assigned to the proper major cell type. The exceptions are renal mesangial cells, which have contractile properties, but are classified as epithelial, and lung epithelial cells, that are classified as mesenchymal. These two cell types, however, are of embryonic origin — in contrast to the vast majority of primary cells in our study, which are adult (Table S1) — and their transcriptomes may not reflect the transcriptomes of fully differentiated cells.

The clustering of primary cells does not appear to be dominated by body location, or embryological origin. Body location actually contributes very little to the expression profile of primary cells, explaining only about 4% of the variance in gene expression (Fig. S2c). Variation of gene expression among organs is similar for the different clusters (Fig. S2d). Remarkably, the transcriptional diversity among cells within a given organ can be as high as that across the entire human body (Fig. S2e). A similar clustering is obtained using FANTOM CAGE-based transcriptomic data on 105 primary cells *(4)* (Fig. 1d, Fig. S3a-b, Table S5), which reveals, in addition, two clusters corresponding to blood and neural cells, which were not represented in our set of primary cells. The analysis of a different set of primary cells from the ENCODE encyclopedia Candidate Regulatory Elements (cREs *(5)*, Table S6), based on DNAse Hypersensitive Sites (DHSs), also recapitulates the clustering (Fig. 1e, Fig. S3c). The clustering remains in the set of 146 non-redundant primary cells, that results from merging the RNA-Seq, the CAGE and the DHS data. The clustering is thus conserved despite the heterogeneity of the underlying assays and experimental protocols used to generate these different data sets (Fig. S4). In the clustering, neural cells (mostly astrocytes from different brain regions and neurons) cluster together with a few neuroepithelial primary cells (we labelled them epithelial, but they are mostly ciliate cells from different sites in the eye). While the neural cells profiled by CAGE seem to have a distinct transcriptional signature (Fig. S3a), neural cells profiled by DNAse-seq exhibit a gene expression pattern similar to mesenchymal cells (Fig. S3c). However, the neural cells profiled by DNAse-seq are, in contrast to most primary cells investigated here, of embryonic origin, and thus they are not likely to express the transcriptional program characteristic of adult neural cells. The analysis of publicly available transcriptomics data from nervous tissues including single-cell and bulk RNA-seq strongly support that the neural cell type is a proper major type clearly differentiated from the other types (Supplementary Information, Fig. S5-S7).

Comparable multi-tissue RNASeq data has become recently available at the single cell level for twenty mouse organs and tissues, through the “Tabula Muris” project *(6)*. Principal Component Analysis (PCA) of the individual cells and hierarchical clustering of the primary cell types show that most individual cells, and most cell types, clustered into the five major cell types above, irrespective of the organ of origin (Fig. S8, S9). As in the case of melanocytes above, we also found a few specialized cell types which do not properly belong to these types. Hepatocytes are a notable example (Fig. S8a, S9a). While closer to the epithelial cells than to cells of other types, they seem to have a quite specialized transcriptional program.

These results, all together, suggest the existence of a limited number of core transcriptional programs encoded in the human genome, and likely in mammalian genomes, in general. These programs underlie the morphology and function common to a few major cellular types, which are at the root of the hierarchy of the many cell types that exist in the human body (Table 1). They all show similar transcriptional heterogeneity, with blood, and epithelial within the solid tissues, being the most transcriptionally diverse (Fig. S10). These transcriptionally defined major cell types correspond broadly, but not exactly, the basic histological types in which tissues are usually classified (see for example *(7–9)*): epithelial, of which endothelial is often considered a subtype, muscular, connective, which includes blood, and neural. However, from the transcriptional standpoint, endothelial constitutes a separate type, closer, if any, to the mesenchymal than to the epithelial type. Blood is also a separate major cell type, while the connective (but not blood) and the muscular histological types cluster together into a single mesenchymal transcriptional type (Fig. 1f).

**Table 1:** Cell types in the human body.

|  |  |
| --- | --- |
| **Cell Type:** | sets of cells with similar phenotype (morphology and functions). The similarity threshold induces a taxonomic hierarchy of cell types, by means of which similar cell types are recursively aggregated into higher order types. |
| **Primary Cell Type:** | cell types at the bottom of the taxonomic hierarchy. They denote specialized cells phenotypically identical (to some resolution); they cannot further be segregated into biologically meaningful subtypes; for example, pancreatic beta cells. In our work, we do not include here, **cell lines**, which are primary cells that have been transformed to proliferate indefinitely. |
| **Major Cell Type:** | cell types at the root of the taxonomic hierarchy. They cannot be further aggregated in biologically meaningful higher order types; for example, epithelial cells. |
| **Tissue-Specific Cell Type:** | cell type topologically restricted to a specific anatomical region (tissue, organ, body location); for instance, hepatocytes. |
| **Transcriptional Program:** | The pattern of gene expression characteristic of a given cell type. |

Within each of the major types, further hierarchical organization of cell types may exist. While we have not profiled enough diversity of primary cells to resolve the taxonomic substructure within each major cell type, hints of this substructure can be clearly seen in the epithelial type. Within the epithelial cluster, two well defined subclusters can be identified (Fig. 1b-e; see also Fig. S2a). One of the clusters is made mostly by renal cells, indicating that body location may actually play a role in subtype specialization. Remarkably, the epithelial cluster includes primary cells of all embryonic origins (ectoderm, endoderm and mesoderm), suggesting that the transcriptional programs of cells may not be fully inherited through development, but partially adopted through function. The more heterogeneous composition of the epithelial type is also apparent in the mouse scRNASeq (Fig. S8, S9).

Our results also suggest that, while many cells are likely to adhere to these basic transcriptional programs, many other primary cells are likely highly specialized and very tissue specific. As with melanocytes and hepatocytes in our analyses, these specialized cells are likely to have their unique transcriptional program.

### Cell type specific genes

We identified a total of 2,871 genes (including 2,463 protein coding genes, 283 long non-coding RNAs and 125 pseudogenes), the expression of which is specific to epithelial, endothelial, mesenchymal or melanocyte cell types (Fig. 2a, Fig. S11, Table S7). These cell type specific genes include nearly all genes that we identified as the major drivers of the clustering (Supplementary Information, Fig. S12). Examples of these genes include collagen (*COL1/3/6*), expressed in mesenchymal cells, epithelial transcription factors genes *OVOL1/2*, *VWF* gene encoding for the endothelial marker von Willebrand Factor, and *TYR* gene encoding for the melanocyte-specific enzyme tyrosinase (see Table S8 for a list of manually curated driver genes). Figure 2b shows the expression pattern of *RP11-536O18.2*, an endothelial specific long non-coding RNA (lncRNA) of unknown function. The gene is expressed in nearly all endothelial cells analyzed here, but not in cells from other types, and its expression is correlated to protein coding genes with endothelial-related functions (Fig. S13a). The gene, however, is expressed in multiple tissues, and, therefore, it is not tissue specific.

**Fig. 2.**
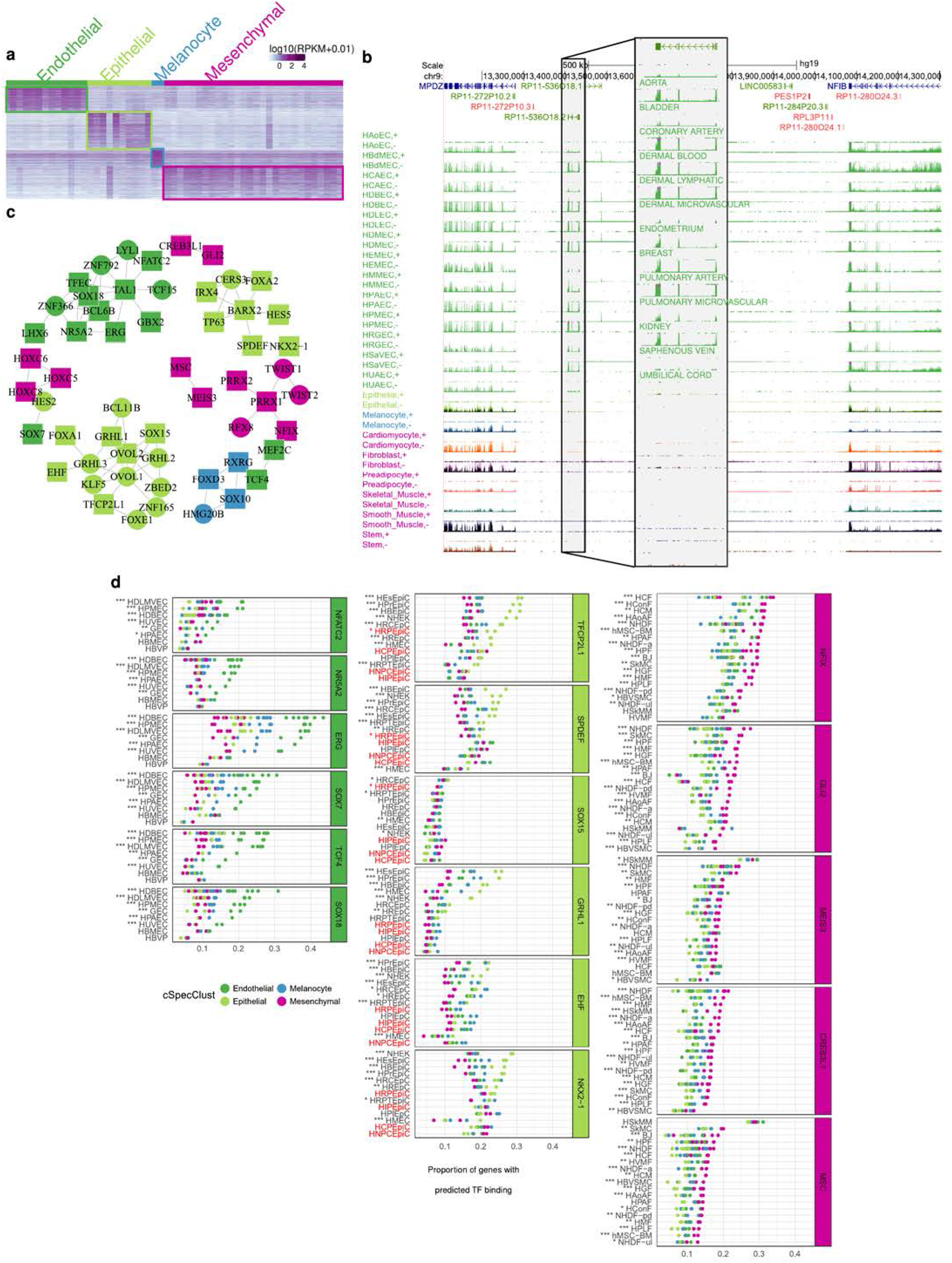
Cell-cluster-specific genes. **(a)** Expression of 2,871 genes specific to major cell types. **(b)** Expression of the endothelial-specific lncRNA RP11-536O18.1. Separate strand-specific signal tracks are shown for endothelial cells, while the other tracks contain overlaid signal for each cell type. The lncRNA has highly correlated (correlation coefficient > 0.9) expression with 72 protein coding genes across our set of primary cells. Nearly all these genes are endothelial specific, and they are functionally enriched for vessel development and angiogenesis (Fig. S13A). The gene appears to be under relatively strong regulation, since it has almost 1,500 eQTLs across multiple tissues in GTEx (v7) well above the average eQTLs for lncRNAs (about 450). **(c)** Network of the most strongly co-expressed (Pearson’s correlation coefficient > 0.85) cell type specific transcription factors (TFs). Nodes are colored according to the cell-type-specificity of the TF, and shaped based on the availability of sequence motif (square: available, circle: not available). **(d)** Proportion of cell type specific genes with predicted TF binding over cell type specific genes that harbor a DHS around their TSS (−10kb/+5kb), individually for each cell type specific TF (with binding motif available) and cell line for which DNAse-seq data was available. In general, we found that genes specific to a given type are enriched for binding motifs for TFs specific to that type. For instance, the proportion of endothelial specific genes with DHS sites that harbor motifs for the endothelial specific TF ERG in dermal blood endothelial cells (HDBEC) is larger than the proportion of genes with DHS sites specific of other major cell types. Primary cells highlighted in red, although included within the epithelial major cell type, they have been labelled as neural/epithelial in Fig. 1d, and they are therefore not proper epithelial; consistently, they do not show the enrichment in binding motifs for epithelial specific transcription factors. Refer to Table S6 for a complete description of the acronyms. Enrichment adjusted p-values: “*” < 0.05, “**” < 0.01, “***” < 0.001.

The functions of annotated tissue-specific genes closely match the expected biology of the primary cells in each type (Fig. S13b). Cell type specific genes show consistent restricted expression in the FANTOM CAGE data (Fig. S14), and they are enriched for encyclopedia cREs *(10)* specifically in the primary cells of that type (Fig. S15). Using ChIP-seq histone modification data obtained in a number of primary cells *(11)* (Supplementary Information, Table S9), we found the promoters of genes specific to a given type to be enriched for activating chromatin marks in primary cells of that type compared with primary cells of different type (Fig. S16a). However, overall, except for H3K4me1, we found low levels of most activating marks in the promoters of cell type specific genes compared with all genes, even after controlling for differences in gene expression. In contrast, the promoters of cell type specific genes exhibit similar or higher levels of repressive histone modifications compared to all genes (Fig. S16b). This is consistent with previous reports showing that genes under tighter regulation show lower levels of activating histone modifications than broadly expressed genes (see for example *(12–13)*).

Among cell type specific genes, we identified 167 Transcription Factors (TFs) from a total of 1,544 TFs annotated in the human genome *(14)*. We focused on 56 that showed the strongest co-expression patterns (Pearson’s correlation coefficient ≥ 0.85, Fig. 2c, Fig. S17). They include previously annotated cell type-specific transcriptional regulators, such as ERG, which has been shown to regulate endothelial cell differentiation *(15)*, and TP63, which is an established regulator of epithelial cell fate and is often altered in tumor cells *(16)*. Consistent with the hypothesis that the cell type specific TFs might regulate cell type specificity, we found that genes specific to a given type are enriched for binding motifs for TFs specific to that type in most cell lines (Fig. 2d). The enrichment arises specifically when the motifs occur in open chromatin domains in primary cells of that type (e.g. in epithelial primary cells, epithelial specific genes are enriched, compared to genes specific to other types, in epithelial specific TF motifs occurring in open chromatin domains, Fig. 2d, Fig. S18).

We found that transcriptional regulation appears to play the major role compared to post-transcriptional (splicing) regulation, both in defining the major cell types as well the individual primary cells within the types. We estimated the fraction of the variation in isoform abundance explained by variation in gene expression *(17)* to be on average 67% across transcriptional types and 55% across primary cells (Fig. 3a). The lower proportion of variance explained across primary cells suggests that splicing plays comparatively a more important role in defining the transcriptomes of primary cells within a given type, than in setting the transcriptional programs of the major cell types. In additional support of this conclusion, we have found that while the number of differentially expressed genes in pairwise comparisons of primary cells is much larger between than within cell types, the number of differentially spliced genes is similar (Fig. 3b, Fig. S19, Supplementary Information).

**Fig. 3.**
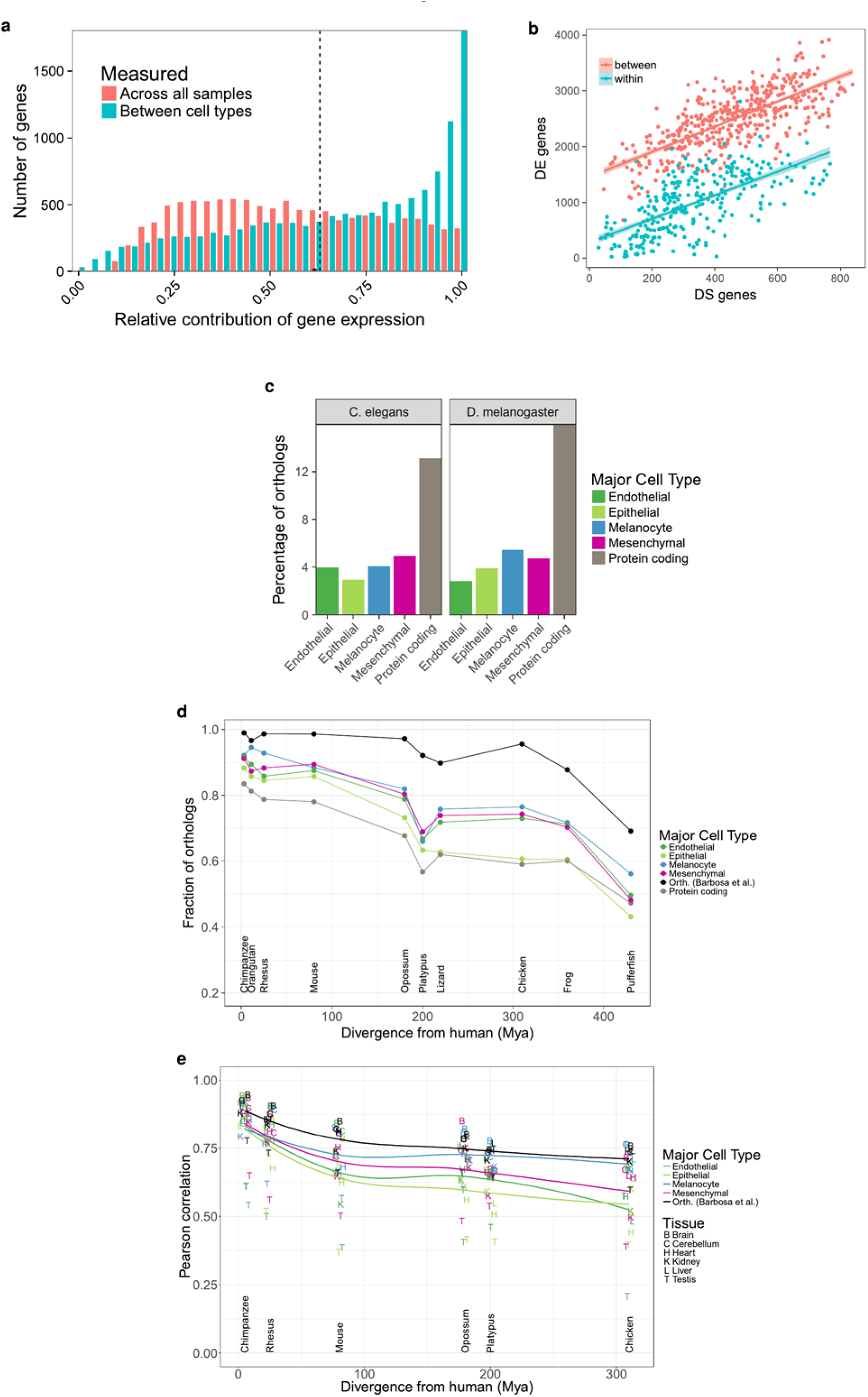
Transcriptional complexity of human primary cells and evolutionary conservation of cell type specific genes. **(a)** Distribution of the relative contribution of gene expression to the variation in isoform abundance between major cell types (blue) and between all primary cells. Large values of the contribution of gene expression indicate that changes in isoform abundance from one condition (primary cell, cell type) to another can be simply explained by changes in gene expression. Small values, by contrast, indicate that changes of isoform abundance are mostly independent of changes in gene expression, and can obey to changes in the relative abundance of the isoform. **(b)** Number of differentially expressed genes (DE, y axis) vs number of genes with differentially spliced exons (DS, x axis), between pairs of samples of the same cell type (within, blue) or different cell types (between, red). DS genes have been obtained using IPSA (https://github.com/pervouchine/ipsa-full). See also Fig. S19. **(c)** Percentage of cell type specific genes and protein coding genes with detected 1 to 1 orthologs in worm (*Caenorhabditis Elegans*) and fly (*Drosophila Melanogaster*). See also Fig. S22a. **(d)** Fraction of 1 to 1 orthologs between each species and human for major cell type specific genes and for protein coding genes overall. Species are sorted by increasing evolutionary distance from human. The black line is given as a reference and it indicates the proportion of 6-way orthologs (chimpanzee, rhesus, mouse, opossum, platypus and chicken) that are present in each species. The proportion is not 100% in these species because different versions of the GENCODE gene set reference were used. The genes in this set of 6-way orthologs are used for the comparison of gene expression in **(c)**. See also Fig. S22b. **(e)** Pearson’s correlation coefficient between gene expression in each human organ and the corresponding one in every other species. The correlation is computed across all the genes in each major cell type separately. See also Fig. S23.

While bulk gene expression is the main contributor to define cell type specificity, other transcriptional events are also cell type specific. First, using the RNA-seq data, we identified cell-type specific splicing events, independent of the tissue of origin (Table S10, Fig. S20, Supplementary Information). Second, using the RAMPAGE data, we identified cell-type specific TSSs (Fig. S21, Table S11, Supplementary Information).

The basic human transcriptional programs seem to have been established early in vertebrate evolution: genes orthologous of cell type specific genes are underrepresented compared to orthologues of all genes in invertebrate genomes (Fig. 3c, Fig. S22a), but they are overrepresented in vertebrates, as early as in tetrapoda. One exception are epithelial genes, which are overrepresented only in mammals (Fig. 3d, Fig. S22b). Within the set of orthologous genes across tetrapoda *(18)*, the expression of cell type specific genes is less conserved than that of protein coding genes overall, especially at larger evolutionary distances (Fig. 3e, Fig. S22c, Fig. S23). This suggests an important role for the evolution of gene expression regulation in shaping the basic transcriptional programs in the human genome. Epithelial specific genes also show the lowest conservation of expression levels. The transcriptional program characteristic of the epithelium appears to be therefore the most dynamic evolutionarily — possibly reflecting a greater need for adaptation of the epithelial layer in constant interaction with the environment, and it is also consistent with the greater transcriptional heterogeneity of this major cell type.

### Estimation of the cellular composition of complex organs from the expression of cell type specific genes

We used the patterns of expression of cell type specific genes to estimate the cellular composition of human tissues and organs from GTEx bulk tissue transcriptome data *(19)* (version 6, 8,555 samples, 31 tissues, 544 individuals). We employed xCell *(20)*, using the sets of genes specific to epithelial, endothelial, and mesenchymal major cell types derived from ENCODE, and specific to brain (neural) and blood derived from GTEx *(21)* as signatures, and computed the enrichments of these cell types in each GTEx tissue sample (Supplementary Information).

The xCell enrichments (Fig. 4a, Table S12) are largely consistent with the histology of the tissues. For instance, esophagus mucosa is enriched for epithelial cells, while esophagus muscularis is enriched for mesenchymal cells. Skin (both exposed and unexposed) is enriched in epithelial cells; fibroblasts, in mesenchymal cells, etc. Blood and brain are only enriched in blood and neural cells, respectively. Most other tissues are not enriched in these two major cell types, with the expected exceptions of spleen enriched in blood cells, and pituitary enriched in neural cells. Testis, which is widespread transcription *(22)*, is also enriched in neural cells, a reflection of the similarity of the expression programs of these two organs *(23)*. Consistent with previous observations *(24)*, we found enrichment of cells of endothelial type in adipose tissue. The analysis of the pathology reports of the subcutaneous adipose tissue shows that often is contaminated with other tissues, in particular blood vessels, which would explain the enrichment in cells of the endothelial type. We have further processed and analyzed the histopathology images available from the GTEx adipose samples (Supplementary Information), and estimated that on average about 84% of the adipose tissue does actually correspond to adipocytes (Fig. S24), which would explain the endothelial enrichment. In skeletal muscle we do not observe a particularly large enrichment in cells of the mesenchymal type, in apparent contradiction with our initial classification (Fig. 1b, f). The samples in GTEx, however, are all from differentiated skeletal muscle, while the ENCODE primary cells that we used to identify the mesenchymal specific genes are undifferentiated satellite cells (SkMC), and smooth muscle cells (Table S1). We analyzed single cell RNA-seq data produced during skeletal myoblast differentiation *(25)*, and found that differentiating skeletal muscle cells retain the mesenchymal signature through most of the differentiation pathway, acquiring only the GTEx muscle specific signature when fully differentiated (Fig. S25a-c). Further supporting that muscle is indeed of mesenchymal type, potentially forming a well-defined subtype, gene expression profiles cluster together myoblast differentiating single cells with ENCODE mesenchymal cells, rather than with epithelial or endothelial cells, or forming a separate cluster (Fig. S25d).

**Fig. 4.**
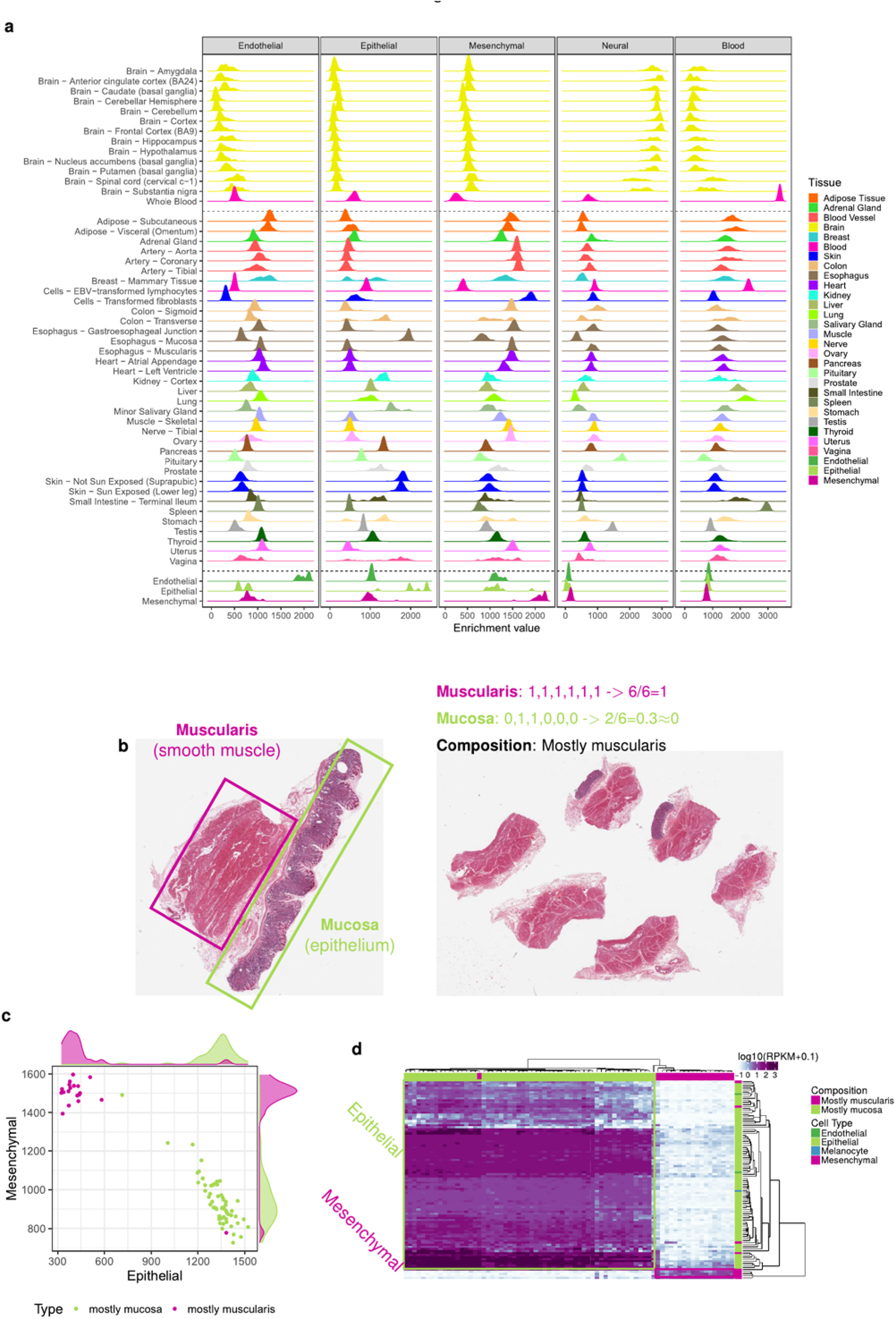
Expression of cell type cluster-specific genes in GTEx organs. **(a)** Enrichment of each major cell type in GTEx tissues, estimated from bulk tissue RNA-seq using the xCell method. As a control, we also include the enrichments in the primary cells monitored here. As expected, the highest enrichment for cells of a particular cell type occurs in cells of that cell type. **(b)** Example of stomach histological slides which represent the two main tissue layers and the procedure for the manual annotation of the images based on the presence of those layers. Each GTEx histological image displays up to six tissue slices. For the stomach samples, we scored each slice for the presence (1) or absence (0) of the muscularis and mucosa layers, summed up the values for each layer separately and divided by the number of slices. If the proportion of slices with mucosa layer, or muscularis layer, is more than 50% we classify the entire slide as mc1, or ms1, respectively. If the proportion is lower, we classify the slide as mc0 or ms0. A combined class, for example mc0ms1, is assigned to the slides. Thus, samples labeled mc0ms1 are mostly muscularis, while samples labelled mc1ms0 are mostly mucosa. **(c)** Enrichment of cells of epithelial and mesenchymal types in stomach samples containing mostly the mucosa (green) or mostly the muscularis (purple) layer. **(d)** Expression of the cell type-specific genes that drive the separation of stomach samples in mostly muscularis or mostly mucosa samples. Among discriminant cell type specific genes, mucosa only samples express almost exclusively epithelial specific genes, while muscularis only samples express exclusively mesenchymal specific genes.

To independently assess the xCell enrichments, we analyzed the histological images of the few tissues in which samples were obtained from different subregions. This is most notable in the case of transverse colon and stomach. The GTEx stomach samples are all from the gastric body, whose walls consist of two broad layers: the mucosa, which is mostly epithelial, and the muscularis, which is smooth muscle (Fig. 4b). We processed the histological images, and identified a subset of samples that presented mostly the muscularis or the mucosa layer (Supplementary Information). This partition of the samples has been also observed by the GTEx consortium using laser capture microdissection (K. Ardlie, personal communication). The enrichment of epithelial cells in the samples from the muscularis layer is much lower than in the samples from the mucosa layer; conversely, the enrichment of mesenchymal cells is much higher in the muscularis than in the mucosa layer. The two sets of samples are almost perfectly separated by our cellular enrichments (Fig. 4c), explaining the bimodality in the distribution of cell type enrichments observed specifically in the stomach samples (Fig. 4a). Consistently, we found that epithelial specific genes were exclusively expressed in the mucosa layer and mesenchymal specific genes were exclusively expressed in the muscularis layer (Fig. 4d). Next, we used the classification of stomach images to train an SVM model (Fig. S26a-b), and used this model to predict the presence of the two layers in 196 transverse colon samples — with histology similar to that of stomach (Supplementary Information). The SVM-predicted classification closely matches the differences observed at the transcriptional level, and confirms that the bimodality of cellular composition (Fig. 4a) is again related to the unbalanced presence of the two tissue layers across samples (Fig. S26c). Considering that stomach and colon were not represented in our primary cell collection, this constitutes a strong validation of our estimates of the cellular enrichments in tissues.

### Alterations of cellular composition in pathological states

We projected the solid non neural GTEx tissue samples on a 3-dimensional space according to the enrichments of epithelial, endothelial and mesenchymal cell types in each sample (Fig. 5a, Fig. S27). The spatial arrangement of the samples recapitulates tissue type as strongly as the clustering based on gene expression (Fig. S28). This suggests that the basic cell type composition is a characteristic signature of tissues, and that departures from this composition may reflect pathological or diseased states. To assess this hypothesis, we analyzed the histological reports associated with the GTEx images (7,911 reports). We employed fuzzy string search and parse trees to convert the natural language annotations produced by the pathologists to annotations in a controlled vocabulary that can be analyzed automatically (Supplementary Information, Table S13). In this way, we identified 19 histological phenotypes affecting one or more tissues for which there were at least 30 affected samples. From these, we identified six conditions with significant (FDR < 0.01) altered contributions of major cell types when comparing composition of affected and normal tissue (Fig. 5b-e)). Atherosclerosis in the tibial artery, which is more prevalent in older donors (Fig. S29a) is associated with an increase in endothelial cells (Fig. 5b); this might be attributed to endothelial proliferation stimulated in peripheral artery occlusion *(26)*. Atrophic skeletal muscle, a phenotype which is also correlated with age (Fig. S29b), is associated with an increase in mesenchymal cells, which is consistent with the reported increase of connective tissue *(27)* and intermuscular fat *(28–29)* in atrophy (Fig. 5c). Indeed, analysis of the pathology reports of GTEx muscle histological images reveals that the proportion of fat is almost twice as high in atrophic than in non-atrophic muscle (24% vs 13%, Supplementary Information). Elevated enrichments of mesenchymal cells are also observed in liver congestion (Fig. S30a), a condition that often precedes fibrosis, which is characterized by an activation of matrix-producing cells, including fibroblasts, fibrocytes and myofibroblasts *(30)*. In spite of the low presence of cells of the major cell types in the testis, we found a further reduction of cells of all these types, mostly endothelial, in testis undergoing spermatogenesis (Fig. S30b). In lung pneumonia, we also observe alteration of all cell types (Fig. S30c). The sixth condition is gynecomastia, a pathology which is characterized by ductal epithelial hyperplasia *(31)*. We investigated differences in cellular composition between males and females, and found them significant only in mammary tissue, where female breasts exhibit much higher enrichment in epithelial cells than male breasts, possibly due to the presence of epithelial ducts and lobules (Fig. 5d). Remarkably, males diagnosed with gynecomastia show a cellular composition similar to that of females, mirroring tissue morphology.

**Fig. 5.**
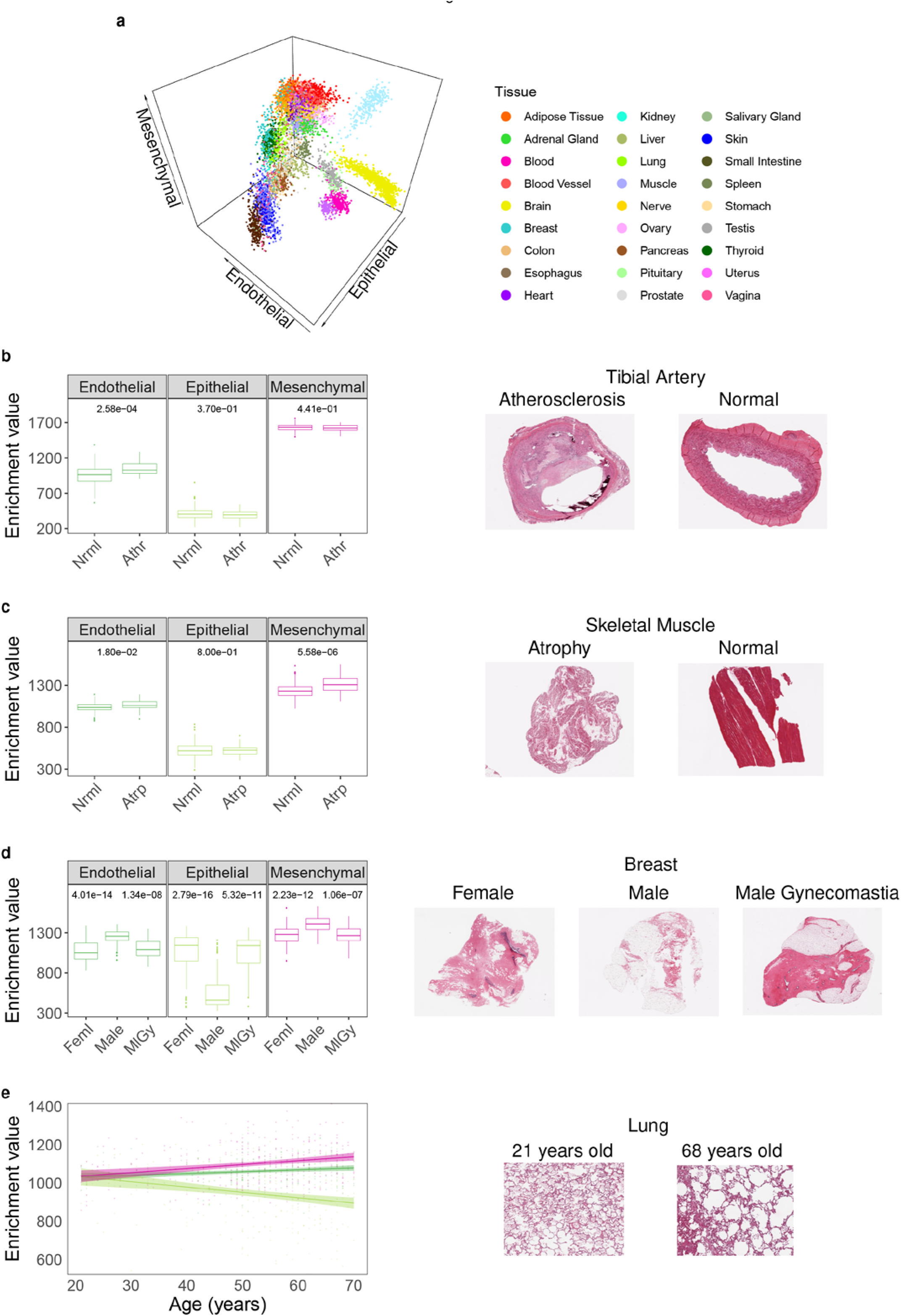
Alterations of the contributions of the major cell types to tissues in histological phenotypes. **(a)** GTEx samples represented in a 3D space where the axes are the enrichments of endothelial, epithelial and mesenchymal cells. (**b** and **c**) Differences in xCell enrichments of major cell types (Wilcoxon test, adjusted p-values as FDR) between affected and normal states. Histological images of affected and normal tissues are displayed (see text for details). Nrml: normal, Athr: atherosclerosis and Atrp: atrophy. **(d)** Major cell type xCell enrichments in female (Fml) breast samples, and male breast samples with (MlGy) or without gynecomastia (Male). Only significant FDR (≤ 0.05) are shown, all of them being between female and male without gynecomastia (left FDR) and between male without gynecomastia and male with gynecomastia (right FDR). **(e)** Changes in major cell type xCell enrichments in lung samples with age. Pearson’s *r* and adjusted p-values as FDR for: Endothelial *r*=0.17 and FDR=3.20e-03, Epithelial *r*=−0.23 and FDR=6.00e-05, Mesenchymal *r*=0.25 and FDR=2.40e-05.

We also observed specific age-related changes in cellular composition in lung and ovarian tissues. In lung samples we observe changes of all cell types, in particular, a significant reduction of epithelial cells in older donors (Fig. 5e), which is consistent with the impaired re-cellularization of lung epithelium that has been observed in decellularized lungs of aged mice *(32)*. Consistently, a similar pattern can be observed in the lungs of the individuals that died of respiratory related causes (Fig. S30d-e). In ovarian samples of women older than 48, a lower bound for menopause occurrence, we observe a decrease in endothelial cells (Fig. S30f), potentially related to an age-dependent decline in ovarian follicle vascularity *(33)*.

Altered cellular composition is likely to be particularly relevant in cancer. We analyzed, therefore, transcriptome data from the Cancer Genome Atlas Pan-Cancer analysis project (PCAWG) *(34)* for 19 cancers affecting tissues also profiled in the GTEx collection, and estimated the cellular enrichments of the major cell types (Fig. S31). For some cases there is also transcriptome data for normal samples from the same cancer project, which serve as a control for the highly different methodologies employed in GTEx and in the cancer projects. Thus, in lung cancer, there is an increase in epithelial cells (Fig. 6a-b), likely reflecting the epithelial origin of most lung cancers. In kidney primary tumors, in contrast, there is an overall increase of endothelial cells across most cancer subtypes, consistent with the increased vascularity associated with the cancer (Fig. 6c-d). The exceptions are renal papillary cell carcinomas, which present, instead, reduced vascularity *(35)*. In both cases, the cellular composition of GTEx samples and normal samples from the cancer projects are similar, supporting the robustness of our cellular characterization. Alterations in cellular composition can also reflect cancer progression. For ovary, even though we lack a comparable set of normal samples from the cancer projects, there is data on different stages of the disease, which serve as an internal control (Fig. 6e-f). Compared to GTEx normal data, there is markedly increase in epithelial cells in cancer, which is more evident as the severity of the cancer progresses, from primary to recurrent.

**Fig. 6.**
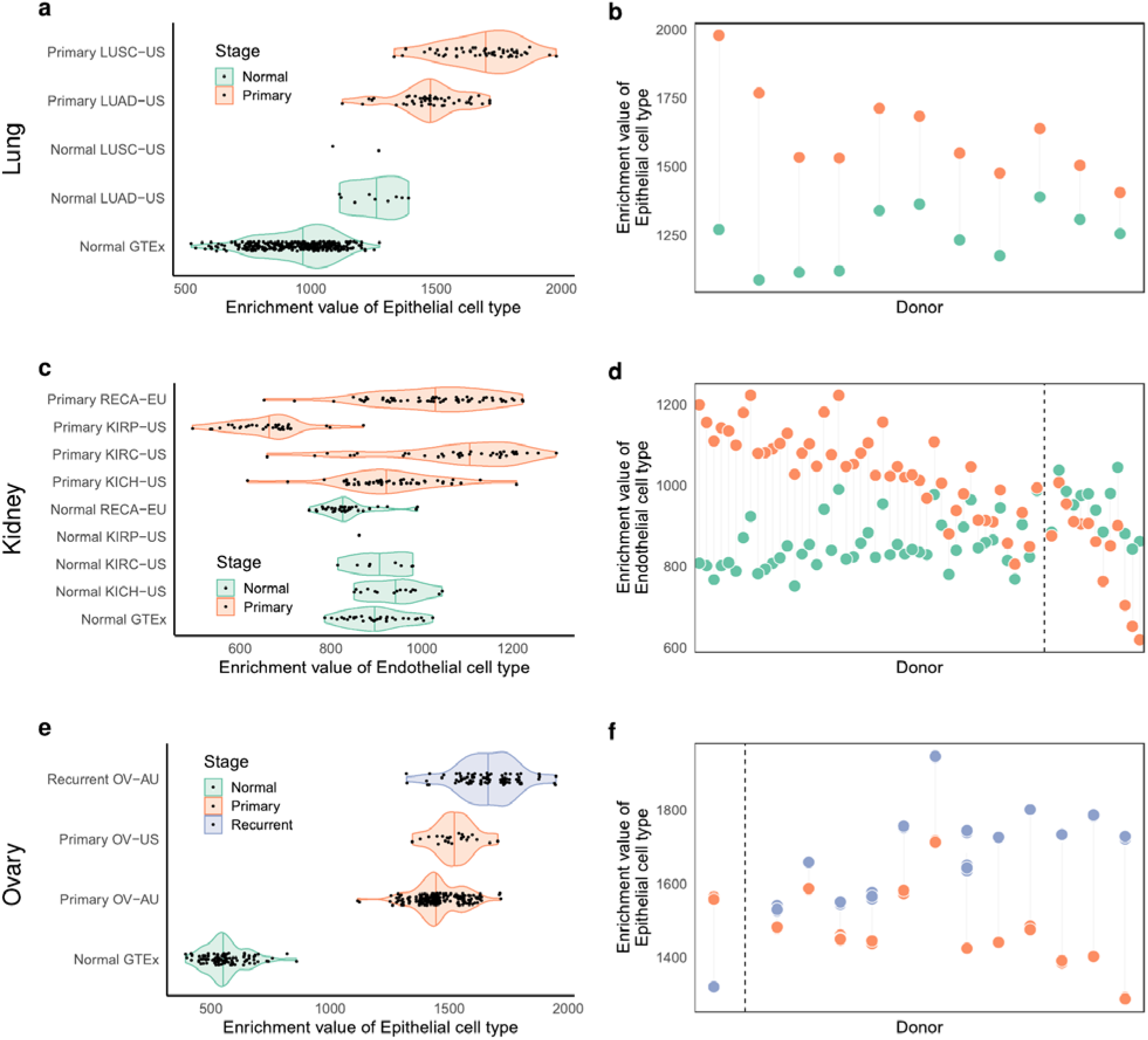
Alterations of the contributions of the major cell types to tissues in cancer. **(a)** xCell enrichments in epithelial cells in lung cancers and matched normal controls from the PCAWG project separated by cancer project. LUAD-US: Lung Adenocarcinoma, TCGA, USA; LUSC-US: Lung Squamous Cell Carcinoma, TCGA, USA. **(b)** Enrichment in matched normal and cancer lung samples by donor, pooled across the cancer projects. The p-value for the Wilcoxon test for the differences in epithelial contribution between normal and cancer samples in the LUAD-US project is: 8.1e-06. **(c)** xCell enrichment in endothelial cells in kidney cancers and matched normal controls from the PCAWG project separated by cancer project. RECA-EU: Renal Cell Cancer, France, EU; KIRP-US: Kidney Renal Papillary Cell Carcinoma, TCGA, USA; KIRC-US: Kidney Renal Clear Cell Carcinoma, TCGA, USA; KICH-US: Kidney Chromophobe, TCGA, USA. **(d)** xCell Enrichments in matched normal and cancer kidney samples by donor. The adjusted p-values for the Wilcoxon tests for the differences in endothelial contribution between normal and cancer samples in the RECA-EU, KIRC-US, KICH-US projects are respectively: 3.8e-12, 0.0024, 0.65. **(e)** xCell enrichments in epithelial cells in ovarian cancers from the PCAWG project separated by cancer project or by **(f)** donor for matched primary and recurrent samples. OV-AU: Ovarian Cancer, Austria; OV-US: Ovarian Serous Cystadenocarcinoma, TCGA, USA. The p-value for the Wilcoxon test for the differences in endothelial contribution between primary and recurrent samples in the OV-AU project is: 3.6e-27. The donors in displays b, d, f are sorted based on the difference between the enrichments. The dashed lines in d, f separate the matched samples in which the enrichment of endothelial (epithelial) cells is larger in the cancer sample from those in which it is larger in the normal sample.

Overall, the data collected here on the transcriptomics of human primary cells constitute a unique resource, serving as an intermediate resolution of complexity between single cell and whole organ transcriptomics. This resource will contribute to the understanding of how the interplay between transcription and cellular composition shapes tissue histology, and ultimately impacts, human phenotypes. Our analyses suggest that a large fraction of human cells and cell types in tissues belong to a few major cell types, providing a high level transcriptionally-based hierarchical classification of human cells. Extending the variety of profiled cell types, achieving single cell resolution and integrating expression data with epigenetics data, as proposed in the Human Cell Atlas project *(3)*, will enrich our understanding of the constitutive cell types in the human body and of their functional relationship.

## Materials and Methods

### RNA Isolation, Library Construction and Sequencing

For each cell type to be made into a library we obtained cell pellets that were stored in RNAlater (Thermofisher) as catalogue items from PromoCell (http://www.promocell.com) and ScienCell (https://www.sciencellonline.com/) (see Table S1 for a list of primary cells). We rely on the providers’ standards for quality assurance. Quality sheets are available through the ENCODE portal (see for example:https://www.encodeproject.org/search/?type=Biosample&organism.scientific_name=Homo+sapiens&biosample_ontology.classification=primary+cell&lab.title=Thomas+Gingeras%2C+CSHL&source.title=PromoCell&award.rfa=ENCODE3). We ordered 3 vials per cell type per donor for a total of 3 million cells. The 3 vials were combined together and we isolated Total RNA from them using the Ambion mirVana miRNA Isolation kit (cat #AM1561). The rRNA was removed using the RiboZero Gold Protocol (cat #RZG1224). The libraries are made using a homebrew “dUTP” protocol *(36)*, which generates stranded libraries. They were sequenced on the Illumina platform in mate-pair fashion and processed though the data processing pipeline at the ENCODE DCC. Additional, information about each of these steps, metadata and files can be found at: https://www.encodeproject.org/.

### RAMPAGE sample preparation

Isolation of RNA is described in the above section. The RAMPAGE protocol *(37)* was used to make libraries. Each library was sequenced in mate-pair fashion on the Illumina platform. Detailed protocol and quality control images and metrics on a per library basis can be found in the “Production Documents” appended to each RAMPAGE assay at the ENCODE Data Coordination center: https://www.encodeproject.org/.

### Small RNA Isolation, Library Construction and Sequencing

Isolation of RNA is described in the above section. The Illumina TruSeq protocol was used to make libraries. Each library was sequenced in single end fashion on the Illumina platform. Detailed protocol and quality control images and metrics on a per library basis can be found in the “Production Documents” appended to each Small RNA assay at the ENCODE Data Coordination center: https://www.encodeproject.org/.

### RNA-seq processing pipeline

Raw reads from the 106 RNA-seq libraries (see Table S1 for a list of ENCODE library ids and https://www.encodeproject.org/ for submitted fastq files) were aligned with STAR v2.3.1z *(38)* to the human genome assembly hg19. Reads mapping to more than 20 multiple positions were discarded. Read counts for all long genes annotated in GENCODE v19 *(39)* were computed with RSEM 1.2.19 *(40)* (expected read counts).

Since for most of the analyses we average expression values for a given pair of replicates and sometimes the two biological replicates are from donors of opposite sex, we remove genes on chromosome Y. The lack of an enrichment step for polyadenylated transcripts preserves the presence of some short biotype genes, which are still longer than 200bp. Thus, we remove genes with at least one transcript annotated as short RNA in GENCODE. These genes are often of repetitive nature which makes the quantification of their expression problematic, this is why we decided to remove them.

Read counts which are not reproducible between two replicates (npIDR > 0.1) *(41)* are set to 0. The matrix of read counts after npIDR is provided as Table S2. After filtering for reproducibility, read counts are normalized to a slightly modified version of RPKM (reads per kilobase of exon model per million mapped reads *(42)*). Specifically, read counts were first normalized to cpm (counts per million), where the library sizes are the TMM (trimmed mean of M values *(43)*) scaled sums of exonic reads, and then normalized by gene length. Finally, RPKM values from the two replicates were averaged, and genes with RPKM<1 in all samples were discarded, resulting in 16,265 genes, including 13,990 protein coding, 1,380 long non-coding RNAs and 895 pseudogenes.

As the samples were prepared and sequenced in three known distinct batches (see Table S1), we used the *removeBatchEffect()* function from R limma package *(44)* to build a linear model with the batch information and the cell types on log10-transformed RPKM (with a pseudocount of 0.01), and we regressed out the batch variable.

## Data and materials availability

All experimental protocols for the samples described here are available on the ENCODE portal www.encodeproject.org. Detailed information about data processing and analyses are available as Supplementary Information. All the data generated for this study are also publicly available on the ENCODE portal www.encodeproject.org. Additional data tables derived from the analyses are included in this published article (and its supplementary information files). GTEx gene expression is available in the GTEx portal at www.gtexportal.org.

## Acknowledgments

We thank Kristin Ardlie and Detlev Arendt for useful discussions. We acknowledge and thank the donors and their families for their generous gifts of organ donation for transplantation and tissue donations for the GTEx research study. The Genotype-Tissue Expression (GTEx) project was supported by the Common Fund of the Office of the Director of the National Institutes of Health (commonfund.nih.gov/GTEx). The content is solely the responsibility of the authors and does not necessarily represent the official views of the National Institutes of Health. We acknowledge the Spanish Ministry of Economy, Industry and Competitiveness (MEIC) to the EMBL partnership.

## Disclosure declaration

The authors declare no competing financial interests.

## Author contributions

A.B., C.A.D., M.M., V.W., R.G. and T.R.G. conceived and designed the experiments and analyses. J.D., C.A.D., A.S. and C.D. performed the experiments. A.B., M.M., V.W., D.G. analyzed the data. J.G., D.D.P., A.V., A.D., C.Z., D.G., F.R., M.P.S. contributed with ideas and statistical advice. A.B., M.M., V.W., R.G. and T.R.G. wrote the manuscript.

## Funding

This project was supported by awards U54HG007004, U41HG007234 and R01MH101814 from the National Human Genome Research Institute of the National Institutes of Health, as well as from the Spanish Ministry of Economy and Competitiveness, Centro de Excelencia Severo Ochoa 2013–2017, SEV-2012-0208, Programa de Ayudas FPI del Ministerio de Economía y Competitividad BES-2012-055848 to A.B., and Ministerio de Educación, Cultura y Deporte, under the FPU programme (Formación de Profesorado Universitario) with pre-doctoral fellowship FPU15/03635 to M.M-A., as well as the support of the CERCA programme / Generalitat de Catalunya. D.G.-M. is supported by a “la Caixa”-Severo Ochoa pre-doctoral fellowship LCF/BQ/SO15/52260001. We would also like to acknowledge support from the European Research Council (ERC) under the European Union’s Seventh Framework Programme (FP7/2007-2013)/ERC grant agreement 294653.

## Supplementary Materials

Supplementary text

Figures S1-S31

Tables S1-S13

References (*45-72*)

