## Supplementary text and figures for "A limited set of transcriptional programs define major cell types"

### 1 Gene expression analyses

#### 1.1 Hierarchical clustering and t-SNE

Hierarchical clustering based on gene expression (Fig. 1b) was performed on log10-transformed RPKM (with a pseudocount of 0.01) after filtering and batch correction. Complete clustering algorithm is applied to the vectors of Pearson's correlation coefficients between each pair of samples. The distance between two vectors is computed as abs(1-cc), where cc is again the Pearson's correlation coefficient between the vectors.

t-distributed stochastic neighbor embedding (t-SNE, R package Rtsne v0.11) is computed over the distance matrix between each pair of samples with perplexity 5, where the distance is calculated as abs(1-cc) on the same log10-transformed RPKM as before.

#### 1.2 Clustering statistics

##### 1.2.1 Silhouette score

For a set of samples partitioned into $k$ clusters, the silhouette score [45] $s_i$ of a sample $i$ is calculated as: $s_i = \frac{(b_i - a_i)}{\max(a_i, b_i)}$, where $a_i$ is the average distance between $i$ and all samples of the cluster to which $i$ belongs to, and $b_i$ being the minimal average distance between $i$ and all samples from the rest of the clusters to which $i$ does not belong. The silhouette score ranges from -1 to 1, where a score of 1 means that the data is clustered accurately. The silhouette score was calculated with the silhouette() function from the *cluster* R package.

##### 1.2.2 Elbow method

The elbow plot [46] for the number of clusters in ENCODE data (Fig. S2b) was generated as follows: first, we perform hierarchical clustering using $1 - cor(X)$ as the distance criterion, where $X$ is the matrix of Pearson's correlation between ENCODE primary cells (Fig. 1b). Then, for each $k \in (2, \ldots, 10)$, we cut the hierarchical clustering tree into $k$ clusters. For each cluster, we compute the sum of the squared differences between the samples and the corresponding centroid. We take the sum of the $k$ scalar values to obtain the total within group sum of squares.

Both the Silhouette and the Elbow methods weakly support four over five clusters, which will result from splitting the epithelial cluster into two. In any case, these two epithelial subclusters can be grouped together into a single cluster, clearly differentiated from the rest of the clusters/types.

#### 1.3 Network modularity

Modularity was computed similarly to what was described in Breschi et al. [47]. Briefly, we built a graph where vertices (or nodes) are samples and where two vertices, samples, are connected if the Pearson's correlation coefficient between the corresponding samples, computed on the gene expression values or the xCell enrichment, is higher than a certain threshold (excluding connections of a sample with itself). The correlation thresholds have been computed independently for the RPKM and the xCell enrichment values (endothelial, epithelial, mesenchymal, neural and blood cell types), and each threshold corresponds to a specific density value, where the density is calculated as the ratio between the number of edges in the graph over the number of possible edges. The density was sampled from 20% to 45% with a step of 2.5%. Like in hierarchical clustering, gene expression values are log10-transformed RPKM after adding a pseudocount of 0.01. When computing the modularity, the membership of each node/sample is the GTEx organ. To compute the modularity we used the modularity() function from the igraph v0.7.1 R package. The modularity is plotted as a function of the density and a line was fitted using the default parameters of the geom_smooth() function from the ggplot2 R package (Fig. S28).

#### 1.4 Estimation of proportions of explained variance

To estimate the proportion of variance explained by the factors body location, germ layer and major cell type, we built a separate linear model for each gene and factor. The proportion of explained variance is usually defined as the ratio of the variance across levels of a factor over the total variance. However, to account for different number of levels in each factor, we compute $\omega^2$, which takes into account also the degrees of freedom of that factor: $\omega^2 = \frac{SSQ-(k-1)MSE}{SST+MSE}$

Where $SSQ$ is the sum of squares for a given factor, $k-1$ is the number of degrees of freedom, $MSE$ is the mean squared error and $SST$ is the total sum of squares.

#### 1.5 Identification of cell type specific genes

Cell cluster specificity was surveyed with the edgeR package [48]. Since edgeR relies on a negative binomial model which requires discrete read counts, for this analysis we used the read counts of the filtered 16,265 genes, and kept the counts for individual samples without averaging the replicates. To find genes specific of each major cell type, endothelial, epithelial, melanocytes, and mesenchymal, we performed pairwise differential expression between samples of a given cluster and all the others. Genes with false discovery rate (FDR) $<$0.01 and at least 4-fold change were considered cell-cluster-specific. The number of cell cluster-specific genes changes depending on the cluster: 627 in the endothelial cluster, 966 epithelial, 438 melanocytes and 840 mesenchymal (2,871 in total, Table S7).

Gene Ontology (GO) enrichment was performed with the R package GOstat [49].

#### 1.6 Neural cell clustering

##### 1.6.1 Identification of neural specific genes

To identify neural specific genes we compared the expression inferred from CAGE data for 11,000 genes between neural cells, including neurons, astrocytes, and neuroepithelial cells, and all other cell types. We retained only genes with more than 50 reads in at least 3 neural samples. For consistency with the identification of the other cell type specific genes, we used the edge R package [48] to perform pairwise differential expression between neural cells and the rest of the samples. We identified 333 neural specific genes with false discovery rate (FDR) $<$0.01 and at least 4-fold change. Gene Ontology (GO) enrichment was performed with the R package GOstat [49].

##### 1.6.2 Sample selection and processing

To perform neural cell clustering we collected publicly available data from GEO (`http://www.ncbi.nlm.nih.gov/gds`) and the ENCODE portal (`https://www.encodeproject.org/`).

Read counts for ENCODE RNA-seq data were obtained for a subset of the initial primary cells with the addition of neural cells from experiments ENCSR968WKR (bipolar neurons originated from GM23338), ENCSR244ISQ (neural progenitor cells originated from H9) and ENCSR233IJT (astrocytes). Read counts for ARPE-19 retinal pigmented epithelial cells and HFF fibroblasts were downloaded from GEO series GSE120891 [50]. Gene ids were mapped between Gencode version 24 to Gencode version 19 to match our annotation. The read counts were concatenated to ENCODE RNA-seq reads and normalized to RPKM as a single matrix following TMM normalization. They are collectively referred to as RNA-seq in Figure S6.

Read counts for single cell RNA-seq from adult brain were collected from GEO series GSE67835 [51]. We relied on the original paper classification for neural cell subtype definition. Read counts for all available samples were normalized to RPKM following TMM normalization. Only postnatal non-hybrid cells were included in the analysis, and cell GSM1657976 was removed, since it was an outlier for number of expressed genes. The average RPKM within each neural subtype was used to summarize the expression values across multiple single cells, when needed. They are collectively referred to as scRNA-seq in Figure S6.

Raw fastq reads from RNA-seq of purified human brain cell populations were collected from GEO series GSE73721 [52]. Since the available processed data included only FPKM calculated from a very different pipeline, we processed the raw reads directly and quantified gene expression with the ENCODE pipeline and parameters. Similarly to the previous datasets, read counts were normalized to RPKM after TMM normalization. These samples are collectively referred to as brainRNA-seq in Figure S6.

Microarray data was obtained for a subset of samples for which DNAse-seq was also available from the ENCODE portal. Raw .CEL files were downloaded from the GEO series GSE19090 [10]. CEL files were converted from CEL format version 3 to CEL format version 4, and genes were quantified with the R packages oligo 1.46.0 [53] and pd.huex.1.0.st.v2 [54] with rma normalization on core targets. Genes probeset ids were mapped to Gencode gene ids with Affymetrix annotation file (`http://www.affymetrix.com/Auth/analysis/downloads/na36/wtexon/HuEx-1_0-st-v2.na36.hg19.transcript.csv.zip`). These samples are collectively referred to as Array in Figure S6.

Because the dynamic range of gene expression measured with microarrays is smaller than with RNA-seq, we applied a modified quantile normalization to enable comparison between the two platforms to generate Figure S6. RNA-seq RPKM and microarray expression values were separately binned into 1000 bins for cell type specific and neural specific genes across all cell types when available in both datasets. Then microarray values were mapped to the RPKM values of the corresponding bin in the ENCODE data. We could not apply other scaling procedures for this specific case, because other RNA-seq data, including single cell and bulk RNA-seq, did not include any cell types other than neural cells, while the rest of RNA-seq and microarray data included endothelial, epithelial and mesenchymal cells as well. Thus, genes specific of the latter cell types, would not be assigned scaled values comparable across the different experimental settings.

Overall, we found that neural specific genes are highly expressed in all neural samples, but not in samples from other major cell types (Fig. S5a,b, Fig. S6), supporting the fact that neural cells constitute a major distinct cell type. Moreover, we found that brain endothelial cells also exhibit a neural specific profile, despite expressing endothelial specific genes as well (Fig. S6). Microglia cells, on the other hand, although still clustering with other neural cells, seem to have a less evident neural specific signature, which could be related to their different embryological origin, from the mesoderm layer instead of neuroectoderm, like the other neural cells (Fig. S6).

##### 1.6.3 Identification of neural subtype specific genes

We used single cell RNA-seq data [51] to identify genes specific of each neural cell subtype, namely astrocytes, neurons, oligodendrocytes, oligodendrocyte precursors (OPC), brain endothelial cells, and microglia. Pairwise differential expression was performed with edgeR between cells each cell type and the rest of the cells. We identified 3,136 differentially expressed genes with false discovery rate (FDR) $<$0.01 and at least 4-fold change, consistently expressed in multiple cells of the same type ($>$ 1 cpm in at least 10 cells). Specifically, we identified 722 astrocyte, 264 brain endothelial, 285 microglia, 1,287 neuron, 174 OPC and 404 oligodendrocyte specific genes, after removing cell type and neural specific genes (Fig. S7). These results show that despite sharing a common neural transcriptional program, different neural cells, express different sets of genes, which further characterize each subtype.

#### 1.7 Identification of cell type specific TFs

We intersected our list of 2,871 cell type specific genes with a list of 1,558 TFs from the AnimalTFDB database [14] to get a set of 167 cell type specific TFs. To identify the most correlated TFs (Fig. 2c) we computed the Pearson's correlation coefficient between log10-transformed RPKM values (pseudocount 0.01) of pairs of TFs across all samples. A final set of 56 TFs had a correlation of at least 0.85 with at least another TF. Of these, 38 have at least one known or novel sequence motif in [55].

##### 1.8 Identification of driver genes through PCA

In addition to differential gene expression, PCA was used to identify driver genes through their loadings. As a result of PCA, each gene is scored for its correlation with each principal component (loadings). Major cell type clusters are better separated in a space defined by more than one principal component: for example, endothelial cells segregate from the other cell types both along PC1 and PC2. Thus, to find driver genes, we computed the cosine similarity between each gene vector of loadings and the centroid of each major cell type in principal component coordinates. We selected 233 genes with cosine $> 0.9$, of which 228 (98%) are cell type specific according to differential expression analysis. The 5 remaining genes, while significantly differentially expressed, did not pass the chosen logFC threshold. To facilitate interpretation, we manually curated a list of 43 driver genes with reported cell type specific functions in the literature (Table S8).

##### 1.9 Relative contribution of gene expression to variability in isoform abundances

Gene expression contribution in the transcript abundance variation across all samples was computed following the methodology presented in Gonzalez-Porta et al. [17] and further improved in Melé et al. [56]. In a nutshell, for each gene, samples are represented in a multidimensional space using their transcript abundances as coordinates. The contribution of gene expression in the transcript abundance variation is computed by the variation of transcript abundance after projecting the samples into a model of constant splicing (a line in the multidimensional space) divided by the total variation of transcript abundance without projection. If this ratio is close to 1, projecting into the ”no splicing” model didn't reduce the transcript abundance variation, pointing at mainly gene expression contribution. Inversely, if close to 0, alternative splicing (or post-transcriptional regulation, in general) is mostly responsible for the major part of the transcript abundance variation.

A generalization of this approach allows to estimate the effect of a given factor, in this case the cell type, to the contribution of gene expression in transcript abundance variation. Precisely, we asked how much of the transcript variation attributed to cell type is due to changes in gene expression. In practice we compare the proportion of variation explained by the cell type classification after and before projecting the samples into the ”no splicing” model. The proportion of variance explained by cell type classification is derived from the classical ANOVA decomposition. The ”no splicing” model is represented by a line in the multidimensional space formed by the different transcripts abundances. Like in the analysis across all samples, a value around 1 means that the projection didn't affect the estimate of variance explained, supporting a full contribution of gene expression. A ratio around 0 means that the variance explained was greatly reduced after projection, supporting a major contribution of alternative splicing.

#### 1.10 Conservation of cell type specific genes

##### 1.10.1 Sequence conservation

As a measure of sequence conservation of cell type specific genes, we computed the fraction of genes in each set which have a one-to-one orthology relationship with human and other ten vertebrates, as well as worm and fly. In Figure 3d, species are sorted by increasing evolutionary distance from human[57]. The list of orthologous genes was retrieved from Ensembl Compara [58] v75, which is compatible to Gencode v19 (`http://feb2014.archive.ensembl.org/biomart/martservice`). As a control, the fraction of orthologs was computed also for all protein coding genes (20,731 genes) and for the set of already defined orthologous genes in Barbosa et al. [18] and filtered in Breschi et al. [47]. As this latter set of orthologous genes was from a previous version of Ensembl, we used a subset of them which are retained through the version 75 (6,268 out of 6,283).

##### 1.10.2 Expression conservation

Using a similar approach to what is described in Breschi et al. [47], expression conservation was measured as the Pearson's correlation coefficient between expression in human and any other vertebrate species in each organ for the different gene sets. Expression data were obtained from Breschi et al. [47, 18]. Only cell type specific genes with orthologous genes in that dataset were used: 207 of 635 endothelial genes, 277 of 950 epithelial, 130 of 353 melanocyte, 295 of 935 mixed.

### 2 Analysis of Cap Analysis of Gene Expression data (CAGE)

Gene expression data from CAGE in human primary cells was obtained using FANTOM5 CAGE data [4] and through a private collaboration. Read counts for each gene were obtained by summing up the read counts for all the promoters of that gene and then normalized to cpm. Only samples that passed FANTOM5 quality filters were retained and two biological replicates were selected at random, when more than two were available for a given cell line, resulting in 142 cell lines. We applied npIDR filtering [41] on read counts between the two replicates, set to 0 the cpm values when the read counts were not reproducible (npIDR>0.1), and averaged the cpm between replicates. Genes with cpm<5 in all samples were discarded, resulting in 21,269 genes.

tSNE was computed on the distance matrix between each pair of samples (Fig. 1d) using log10-transformed cpm after adding a pseudocount of 1. The distance between two samples is computed as abs(1-cc), where cc is again the Pearson's correlation coefficient between the vectors of peak quantifications.

PCA was computed on the same cpm values as tSNE, after mean-centering and scaling for each gene across all samples (Fig. S1c).

### 3 Analysis of DNase-seq data

#### 3.1 DHS filtering

DHSs for 127 human primary cells were downloaded from the ENCODE portal as narrowPeak files (Table S6). The peaks were filtered within each sample to remove the ones with a read count lower than the minimum number of reads for which there is the maximum p-value. The filtered peaks are merged across experiment if they are separated by at most one nucleotide, resulting in 742,099 merged peaks. Read counts are normalized to counts per million with TMM normalization [43]. The sum of read counts within each sample after filtering is used as library size. Finally, we retained 555,693 peaks after removing the ones that do not have 1 cpm in at least two samples.

#### 3.2 tSNE and PCA based on DHSs

As with CAGE peaks, tSNE was computed on the distance matrix between each pair of samples (Fig. 1e) using log10-transformed cpm after adding a pseudocount of 1. The distance between two samples is computed as abs(1-cc), where cc is again the Pearson's correlation coefficient between the vectors of peak quantifications.

PCA was computed on the same cpm values as tSNE, after mean-centering and scaling for each gene across all samples (Fig. S1b).

##### 3.3 TF binding prediction in DHSs

We used the 742,099 merged peaks to look for cell type specific binding sites. First, for each DNase-seq sample we computed the proportion of cell type specific genes with a peak within their TSS region, extended 10kb upstream and 5kb downstream. TSSs for all transcripts of a given gene are considered to take into account for alternative TSS usage. Next, we scanned the peaks for known and novel motifs [55] of our defined cell type specific TF, with FIMO [59] (default parameters). We used CENTIPEDE [60] to filter for motif hits with high likelihood of predicted binding (simple model, lambda=0.1, posterior probability $> 0.95$). The proportions shown in Figure 2d for each sample are calculated as the number of cell type specific genes with predicted TF binding in their TSS over the total number of cell type specific genes with DNase peaks in their TSS.

#### 4 Analysis of chromatin marks

ChIP-seq histone modification data was obtained from the ENCODE portal [11] for 15 cell lines for which RNA-seq data was also available (Table S9). The data was generated with antibodies for the following histone marks: H3K27ac, H3K27me3, H3K36me3, H3K4me1, H3K4me2, H3K4me3, H3K79me2, H3K9ac, H3K9me1, H3K9me3. For each gene in each experiment, we averaged the ChIP-seq signal every 10 nucleotides in the region around the TSS (+/- 2kb), with the software bwtool [61].

To assess whether cell type specific genes have stronger activating marks (H3K4me3, H3K27ac) in the corresponding cell type compared to the other cell types (Fig. S16), we normalized the signal over the sum of the signal at the TSS of all genes.

To compare chromatin modifications for the different sets of cell type specific genes (Fig. S16b), we selected a subset of genes, that are not cell type specific, but have comparable expression levels to each set of cell type specific genes separately. This was achieved by binning the reference set of cell type specific genes into 200 bins of expression and by randomly selecting an equivalent number of non cell type specific genes belonging to those expression bins. In addition, because we aimed to compare chromatin signal within the same cell, the signal was normalized between 0 and 1, so within-sample differences are on similar scales across samples.

### 5 Splicing

#### 5.1 Computation of exon inclusion levels

Exon inclusion levels, measured as percent spliced-in (PSI) values [62], were computed for 439,778 internal exons with IPSA Splicing Analysis Pipeline (https://github.com/pervouchine/ipsa). PSI values were computed only for splicing events supported by at least 10 reads. PSI values for exons with an absolute difference in PSI values larger than 0.1 between two replicates were set to NA in both samples. PSI values were averaged between replicates, when available.

We also used LeafCutter [63] with the default parameters and the protocol described by the authors. First, we converted the BAM files described in the ”RNA-seq processing pipeline” section into junction files. Then, we perform intron clustering using the default parameters of 30 minimum reads in a cluster, 100,000 bp as maximum intron length and 0.001 for minimal fraction of reads in a cluster that support a junction. In order to be consistent with the analysis on IPSA, the average intron clustering value is computed for each pair of replicates.

#### 5.2 Gene Expression vs Exon inclusion

We compared the differences of gene expression and of exon inclusion within and between cell types. The difference of gene expression within and between cell types was computed as the number of genes differentially expressed (edgeR [48], FDR$<$0.01, log2-fold-change$>$2) between each pair of samples belonging to the same or different cell types, respectively. The difference of exon inclusion within and between cell types, instead, was computed as the number of genes containing at least one exon differentially included between pairs of the same or different cell types, respectively, when using IPSA. We defined an exon as differentially included between two samples if the absolute difference between its PSIs in the two samples was larger than 0.2. We also used other cutoffs, 0.1, 0.3, 0.4, 0.5, obtaining similar results (Fig. S19b). Finally, we computed the ratio between the number of differentially expressed genes (DE genes) and the number of genes with differentially included exons (DS genes) for each pair of samples. As there was evident bias between PSI values from the first two batches compared to the third, we restricted the analysis to the first and the second batches, which include enough representative samples for the different cell types (Fig. 3b).

For LeafCutter, the differential splicing (DS) analysis has then been performed using the Leaf-

Cutter script between each pair of primary cell lines, using a threshold on the FDR of 0.01 to define the differential intron cluster. The DS analysis has been run using the following parameters: *–min_samples_per_group=1* and *–min_samples_per_intron=1* to be able to run DS with only two samples. Like for IPSA, a gene is considered as DS if it contains at least one differentially spliced intron cluster with a deltaPSI $>$ 0.2, and only the first and second batches were used.

##### 5.3 Cell type specific alternative splicing

We compute event-specific PSI values for well defined alternative splicing events using an in-house script, from exclusion and inclusion read counts obtained by the IPSA pipeline. Alternative splicing events were defined based on the annotation between pairs of transcripts with the AStalavista tool [64]. We ensure reproducibility between replicates by requiring that the event is supported by at least 10 reads in both replicates and that the difference in PSI between the replicates is less 0.15. PSI values are then averaged between replicates. Events that do not pass these filters are assigned NA in that pair of sample. Because we are interested in cell type specific events, we filter for events with no more than 75% missing values of the samples within each cell type. Events are further filtered by total variance $>$ 0.01 across all samples. Finally, missing values are imputed for the remaining 1,113 events with impute.knn() function [65] from the R package impute which employs the k-nearest neighbors algorithm. The 1,113 events are tested for cell type specificity with the Kruskal-Wallis test, a non-parametric alternative to one-way ANOVA, and p-values are corrected for multiple testing with the Benjamini-Hochberg procedure. With 1% FDR we identified: 3 alternative acceptor events, 3 alternative donor events, 46 multiple exon skipping events, 14 mutually exclusive exon events, 7 retained introns and 157 single cassette exons (Table S10). Sashimi plots were created with ggsashimi [66]. In Figure S20c, the exon has been previously reported to be more often included in muscle cells [67].

#### 6 Analysis of alternative TSS usage with RAMPAGE data

We analyzed RAMPAGE data from 28 experiments, corresponding to 14 primary cells with 2 replicates. The TSS peaks were downloaded from the ENCODE portal (Table S3) in the gff format. The TSS peaks are associated to genomic regions, but not directly to genes. In order to associate a TSS peak to a gene, we assign the TSS to the closest gene with the largest overlap with the associated genomic region. If the closest gene is farther than 10kb, the TSS is annotated as novel and is not assigned to any gene. TSSs that are overlapping at least by one nucleotide are merged across samples, yielding 34,802 TSSs. Read counts are normalized to counts per million of mapped reads. We further retained 9,068 genes (16,918 TSSs), which have at least one TSS with cpm $> 0.5$ in at least 2 samples. Of these, 3,856 genes which had more than one TSS were tested for differential TSS usage with an approach similar to the one described in the previous section.

For each gene, we estimated the variance in TSS quantifications between cell types that can be attributed to changes in gene expression (var.bwGp.ge) or to changes in TSS usage (var.bwGp). To select genes with differential TSS usage between cell types, we binned them based on their variation in gene expression (var.bwGp.ge) in 10 quantiles and computed the Z-score of the variation in TSS usage with respect to each quantile (Fig. S21). We defined 17 genes whose variation in TSS usage is larger than the variation in expression and with a Z-score $> 1.5$.

### 7 Analysis of GTEx data

#### 7.1 Selection of tissue-specific genes

Tissue-specific genes were selected according to a tissue-specificity (ts) score described in [21], which accounts for the fact that genes can be expressed in a selective way in multiple tissues rather than just in a single one. In order to select only the genes with restricted expression to few tissues, we used a more stringent threshold than the one used in the original work (ts score$>$4 instead of ts score$>$3). In addition, genes are considered specific of a given tissue if they have a median RPKM$>$1 across the samples of that tissue. These filters led to a set of 10,365 tissue-specific genes for all the GTEx tissues and sub-tissues, most of which are testis and brain specific (4,693 and 1,980 genes, respectively).

#### 7.2 Estimation of cellular enrichments

In this analysis we aimed to estimate the cellular enrichments of the five major cell types described so far in whole organs, using gene expression data from the GTEx consortium [68].

To this end, we used the xCell 1.1.0 R package, which is signature-based cellular enrichment method that depends on lists of marker genes that are representative of a given cell type. The methodology details for the enrichment score estimation can be found in the original xCell publication [20]. We replace the xCell default signature gene sets by a signature composed of six gene sets: four sets correspond to the genes specific to three major cell types (endothelial, epithelial and mesenchymal), and melanocytes, obtained as previously described (see section ”Identification of cell type specific genes”). The fifth set corresponds to the Neural cell type, which is the union of the genes specific to the 13 GTEx brain subregions (see section ”Selection of tissue-specific genes”). The last set corresponds to the Blood major cell type, and it is the set of genes specific to the Whole Blood tissue (see section ”Selection of tissue-specific genes”).

xCell authors state that ”xCell performs best with heterogeneous datasets. Thus it is recommended to use all data combined in one run, and not break down to pieces (especially not cases and control in different runs)” (see `http://xcell.ucsf.edu/`). For this reason, we have combined the gene expression matrices of ENCODE primary cells, GTEx tissues, and PCAWG samples into a single matrix in order to perform the enrichment estimation, keeping only the genes that are present in at least one of the six gene sets.

Results are presented in Figure 4a. As a control, we also include the enrichments in the ENCODE primary cells monitored here. As expected, these samples have the highest enrichment for major cell types (endothelial, epithelial and mesenchymal) to which they correspond. Since xCell works on ranks, it follows that the enrichment of the GTEx and PCAWG samples should be between 0 and the enrichment of the ENCODE samples, as they should be the ”maximally enriched” ones, providing a fair way to compare tissue samples within the same major cell type. However, it is important to realize that these enrichment scores are not comparable across cell types (for example, an enrichment score of 1000 in Endothelial cell type might not be comparable with an enrichment score of 1000 in Epithelial cell type). The same applies for brain and whole blood GTEx samples and their Neural and Blood major cell type enrichment estimations.

We projected the tissue samples on a three-dimensional space where the axes correspond to the enrichment for the major cell types (endothelial, epithelial and mesenchymal). The projection shows that each tissue has its own characteristic cellular composition (Fig. 5a). This is observed as well when performing dimensionality reduction (tSNE and UMAP) over the enrichment scores of the five major cell types (Fig. S27).

The matrix of RPKM values for tissues was downloaded from the GTEx portal (`http://www.gtexportal.org/home/datasets/`, file: `GTEx_Analysis_v6_RNA-seq_RNA-SeQCv1.1.8_gene_rpkm.gct.gz`) for 8,527 samples (after filtering) and 56,319 genes of the v6 release. Samples extracted from tissues with less than 20 individuals were removed from the analysis.

These included bladder, cervix and fallopian tubes.

##### 7.3 Classification of stomach histological samples

Histological images for the corresponding samples with RNA-seq data can be publicly found on the Biospecimen Research Database website (`https://brd.nci.nih.gov/brd/image-search/searchhome`). Images of histological slides of stomach sections were classified based on the presence of the mucosa and muscularis layers (Fig. 4b). To each slide we manually assign two binary vectors, one for the mucosa and one for the muscularis layer, where 1 and 0 indicate presence or absence of the layer, respectively. The length of the vectors depends on the number of tissue sections on the slide, and the order reflect the order of the tissue sections from left to right and from top to bottom. Then, we computed the proportion of sections with a given layer and rounded the proportion to be binary. Thus, each slide will have one of the following possible compositions: mc1ms1, mc1ms0, mc0ms1, mc0ms0. Finally, we focused on the samples with either one or the other layer, i.e. mc1ms0 (mostly mucosa) and mc0ms1 (mostly muscularis).

##### 7.4 Identification of variable genes among stomach samples

To find the genes at the base of the transcriptional differences we observed amongst stomach samples, we selected in an unsupervised fashion the most variable genes, by using the projection score [69]. We identified 500 most variable genes which maximize the projection score for the first three principal components. Of these, 96 are cell type specific (84 epithelial-specific, 9 mesenchymal, 2 endothelial, 1 melanocyte), according to our definition. The expression of these 96 genes clearly discriminates stomach samples with only mucosa or muscularis layer (Fig. 4d).

##### 7.5 Automated classification of colon samples

We used a machine learning approach to automatically classify the histological slides based on the presence of the mucosa and/or the muscularis layer.

First, we implemented a script in Python 2.7 using the OpenCV2 library in order to preprocess the images, find the tissue piece contours, and create binary masks that allowed to save each piece into a separate image. Because the two layers have distinctive color patterns in the histological slides, we used all the channels from the RGB color model and the first two channels from the

HSV color model as features. For each image, we computed the normalized histogram for each color channel. The concatenation of these histograms will be the feature set for each observation.

We used the manually annotated labels (muscularis/mucosa/both) for stomach samples to train a support vector machine (SVM) model with radial basis function (RBF) kernel. The SVM was fitted with 10-fold cross validation using the caret package in R. 75% of the samples (550 images) were used to train and validate the model and the other 25% (182 images) for testing, obtaining an accuracy of 87% over the test set (Fig. S26b). The 95% CI for the prediction is (81.64%, 91.82%), the No Information Rate (NIR, which is the proportion of the largest class) is 48.35%, and the p-value [Acc $>$ NIR] is $< 2.2 \times 10^{-16}$.

Since the color composition of stomach and colon tissues is very similar, we applied the classifier to colon pieces as well in order to produce the class labels for each image. The descriptive features for colon were generated in the same way as the ones for stomach.

#### 7.6 Analysis of single cell RNA-seq data

##### 7.6.1 Analysis of muscle single cell RNA-seq data

Muscle single cell RNA-seq data was downloaded from the R package *HSMMSingleCell*[25]. The data consists of gene expression measurements from an induced differentiation of primary human myoblasts and was processed following the authors' guidelines (`http://monocle-bio.sourceforge.net/monocle-vignette.pdf`). Briefly, the gene expression matrix was filtered to remove lowly expressed genes and cells with few detected genes. Differentially expressed genes to use as reference for the pseudotemporal sorting were inferred from a predefined list of 21 marker genes known in myoblast differentiation. Pseudotemporal sorting was performed with the R package 'monocle' v0.99.0 [25]. For cell type specific genes and muscle specific genes from GTEx, the Pearson's correlation coefficient is computed between their expression in each single cell and the average expression from the muscle gene signature matrix, i.e. the average expression within each major cell type and median expression in skeletal muscle samples from GTEx (Fig. S25c).

tSNE was performed on the joined gene expression matrix for single cells and for the ENCODE primary cells, after normalization. Specifically, for ENCODE data, the batch-normalized gene expression matrix was used, generated as described above. The expression values for each gene of matrices were then mean-centered and scaled separately for each matrix. Principal component analysis was used to reduce the dimensionality of the joined dataset. Finally, tSNE was applied on the resulting principal components with a perplexity of 10 using the R package *Rtsne* v0.13 [70].

##### 7.6.2 Analysis of Tabula Muris data

The Tabula Muris project generated a compendium of single-cell transcriptomic data from more than 100,000 cells isolated from 20 organs from three female and four male three-month-old mice [6]. For each mouse, several organs were analyzed, controlling in this way for age, environment and epigenetic effects, enabling direct comparison of cell type composition between organs. Two distinct technological approaches were employed: microfluidic droplet-based 3'-end counting using the 10x Genomics platform, and FACS-based full length transcript analysis using SMART-Seq2 libraries (hereafter "10x" and "FACS", respectively). These platforms generated gene expression information of 55,656 and 44,949 cells, assigned to 55 and 81 known cell types, respectively.

To preprocess the data, we used the original "data ingest" scripts provided by the Tabula Muris Consortium (`https://github.com/czbiohub/tabula-muris`), but excluding cells that were labeled generically: bladder cells (1,203 in 10x and 695 in FACS), kidney cells (45 in 10x), and cells with no assigned ontology class (791 in 10x and 170 in FACS). This results in 53,617 and 44,084 cells, grouped into 53 and 80 cell types, respectively. The generated *Seurat* objects were upgraded to version 3.0.1 using the *UpgradeSeuratObject* function. For downstream analyses, only the sets of variable genes identified by *Seurat* across the cells were used (2,010 and 4,694 genes, respectively).

We assigned to the cells a major cell type (blood, epithelial, mesenchymal and endothelial, plus neural in the case of the FACS data) based on the known histology of the cells. Cells with specialized functions were annotated as tissue-specific (TS) cells, and were not assigned a major cell type category. In 10x, these were bladder urothelial cells, mast cells and neuroendocrine cells; for FACS, this was the case for 11 cell types: Kupffer cells, Slamf1-negative multipotent progenitor cells, Slamf1-positive multipotent progenitor cells, pancreatic A cells, pancreatic D cells, pancreatic PP cells, pancreatic acinar cells, pancreatic ductal cells, pancreatic stellate cells, professional antigen presenting cells, and type B pancreatic cells.

PCA embeddings of the cells based on the expression of variable genes were computed using *Seurat* (Fig. S8a and S9a). Most cells clearly grouped into the blood, mesenchymal, endothelial, and epithelial major cell types, plus neural in the case of FACS. The exception was hepatocytes (1,764 and 391 cells in 10x and FACS, respectively) which formed a clearly separated transcriptional group. For this reason, we excluded them from further analyses, recomputed variable genes (1,894 and 4,671 for 10x and FACS, respectively) and regenerated the PCA for the cells in both platforms (Fig. S8b an S9b).

We computed Spearman's correlation between the mean expression profiles of the cells assigned to each cell type - tissue pair, and then performed hierarchical clustering using Ward2's clustering criterion with euclidean distance (Fig. S8c and S9c).

In 10x cells, nearly all cell types clustered within the corresponding major cell types, with the exception being cardiac muscle cells which we labelled as mesenchymal, but cluster within the epithelial types. In FACS cells, most cells also properly clustered within the major cell types. Indeed, blood, mesenchymal, endothelial and epithelial cells clustered mostly within their major cell types. Neural cells clustered together with different types of pancreatic endocrine cells, consistent with the strong morphological, and physiological similarities between these two types of cells [71], and with a few epithelial cells. This resembles the neuroepithelial cluster that we found in human. The exception is oligodendrocytes, that we labelled as neural, but that unexpectedly cluster within the epithelial type. Finally, a very few cell types that we labelled as mesenchymal and epithelial clustered together separated from the rest, but connected to the epithelial cluster.

#### 7.7 Relationship between estimated cellular enrichments and histological phenotypes

##### 7.7.1 Parsing histopathological annotation

Each GTEx histological image has an associated free-form text comment detailing the findings of the pathology review. We seek to automatically recover morphological and/or medical phenotype information from these comments and structure it in a friendly way to perform statistical analyses, however, the free-text poses a challenge since the format is inconsistent and difficult to standardize. To solve these issues, we developed a text processing procedure in Python 2.7. First, we search which annotations contain a specific phenotype within a certain tolerance threshold, using Levenshtein distance to account for typographical errors (insertions, deletions, substitutions). Next, we perform part-of-speech tagging to find dependencies between the words, using the TextBlob NLP library. With these part-of-speech tags, we can build rules to keep words on which we are interested. Using this methodology we are able to reshape automatically many of the free-form text annotations into a friendly format to perform statistical analyses (Table S13), however, a small fraction of the annotations still had to be manually curated due to some annotations not having a correct part-of-speech tag.

##### 7.7.2 Testing the relationship

We tested the significance of alterations in the xCell enrichments for 61 phenotypes. We performed a Wilcoxon test, and corrected for multiple testing (Benjamini-Hochberg correction [72]).

#### 7.8 Estimation of fat proportions in muscle based on pathology annotations

Each histological image is accompanied by a pathology review comment. For the case of muscle tissue, many of these comments contain a numerical estimation of the proportion of fat present in the tissue. We were able to extract these estimations using regular expressions and spaCy 2.0.7, which is a Python API for text processing. For samples that contain fat estimations that are expressed as a range (for example, 5%-10%) we take the midpoint (7.5%) as an approximation of the value for the sample. Once we have the estimates at the sample level, we compute the summary statistics stratifying by atrophic and non-atrophic samples.

#### 7.9 Estimation of fat proportion in Adipose - Subcutaneous histology images

Each histology image is encoded as an SVS file which contains several layers that contain the image at different resolutions. We extract a downscaled version of the histology images from 305 SVS files corresponding to Adipose - Subcutaneous samples in order to perform the estimation. After which, we preprocess the images to delete small components and reduce noise. Then, we generate a mask over the regions of interest, which are the tissue cuts. We infer the two most dominant colors inside the masked regions, and then we assign to each pixel in the masked region one of these two most dominant colors with respect to their color proximity. Once the image has been transformed, we can calculate the proportions of the colors to obtain an estimation of the proportion of fat. With this procedure, we obtain a mean fat proportion of 84%. All the image manipulations were performed using Mathematica 11.0.

#### 8 Analysis of cancer expression data

We used gene expression data for 1,963 cancer and normal samples, measured as fragments per million mapped reads (FPKM), from the Cancer Genome Atlas Pan-Cancer analysis project (PCAWG) [34] encompassing 30 cancer sequencing projects and 20 organs and tissues. To have a comparable expression measure to the other data sets in this study, we converted FPKM to RPKM by multiplying each value by a factor of 2. The cellular enrichments of cancer and normal samples from the PCAWG project were estimated at the same time that the GTEx enrichments, as previously described.

### Supplementary Tables and Figures

**Table S1.** ENCODE long RNA-seq experiments from human primary cells. The column ”labExpId” contains the library id which can be used to uniquely identify the samples on the ENCODE portal `http://www.encodeproject.org`.

**Table S2.** Gene expression matrix of expected read counts after npIDR.

**Table S3.** ENCODE RAMPAGE experiments from human primary cells. The column ”labExpId” contains the library id which can be used to uniquely identify the samples on the ENCODE portal

**Table S4.** ENCODE short RNA-seq experiments from human primary cells. The column ”labExpId” contains the library id which can be used to uniquely identify the samples on the ENCODE portal

**Table S5.** List of CAGE experiments from the FANTOM consortium. The column ”labExpId” contains the unique library identifier.

**Table S6.** List of DNAse-seq ENCODE experiments from human primary cells. The column ”fileId” contains the id of the bigbed files, which can be used to uniquely identify the samples on the ENCODE portal.

**Table S7.** List of 2,871 cell type specific genes.

**Table S8.** List of 43 driver genes selected with PCA with literature references to their cell type specificity. Columns are: gene id, gene type, gene name, cosine, major cell type, PMID.

**Table S9.** List of ChIP-seq samples with antibodies for histone marks. Only experiments with corresponding RNA-seq data were included.

**Table S10.** Exons involved in 230 cell type specific alternative splicing events.

**Table S11.** List of 17 genes with cell type specific TSS usage by RAMPAGE. The coordinates of all detected TSSs and the gene ids are provided.

**Table S12.** Estimated cell type enrichments using xCell for 8,527 GTEx samples, 1,963 PANCANCER samples and 53 ENCODE samples.

**Table S13.** Histopathological annotations of 7,911 GTEx tissue slides, classified with fuzzy string search.

**Fig. S1**. Principal component analysis (PCA) replicates the clustering and t-SNE of the primary cells samples. While PCA of the ENCODE RNA-seq and DNA-seq data reveals the major cell types, the PCA of the FANTOM data is dominated by the separation of the blood samples from the rest. **a)** Principal component analysis of the ENCODE human primary cells, based on RNA-seq data. **b)** Principal component analysis of ENCODE human primary cells, based on DNAse-seq peaks. **c)** Principal component analysis of FANTOM human primary cells, based on CAGE data. **d)** Principal component analysis of FANTOM human primary cells, without blood cells, based on CAGE expression data of 209 genes selected with the projection score[67]. The projection score removes non-informative variables (i.e. genes, in our case) including only the optimal number in the contexts of visualization.

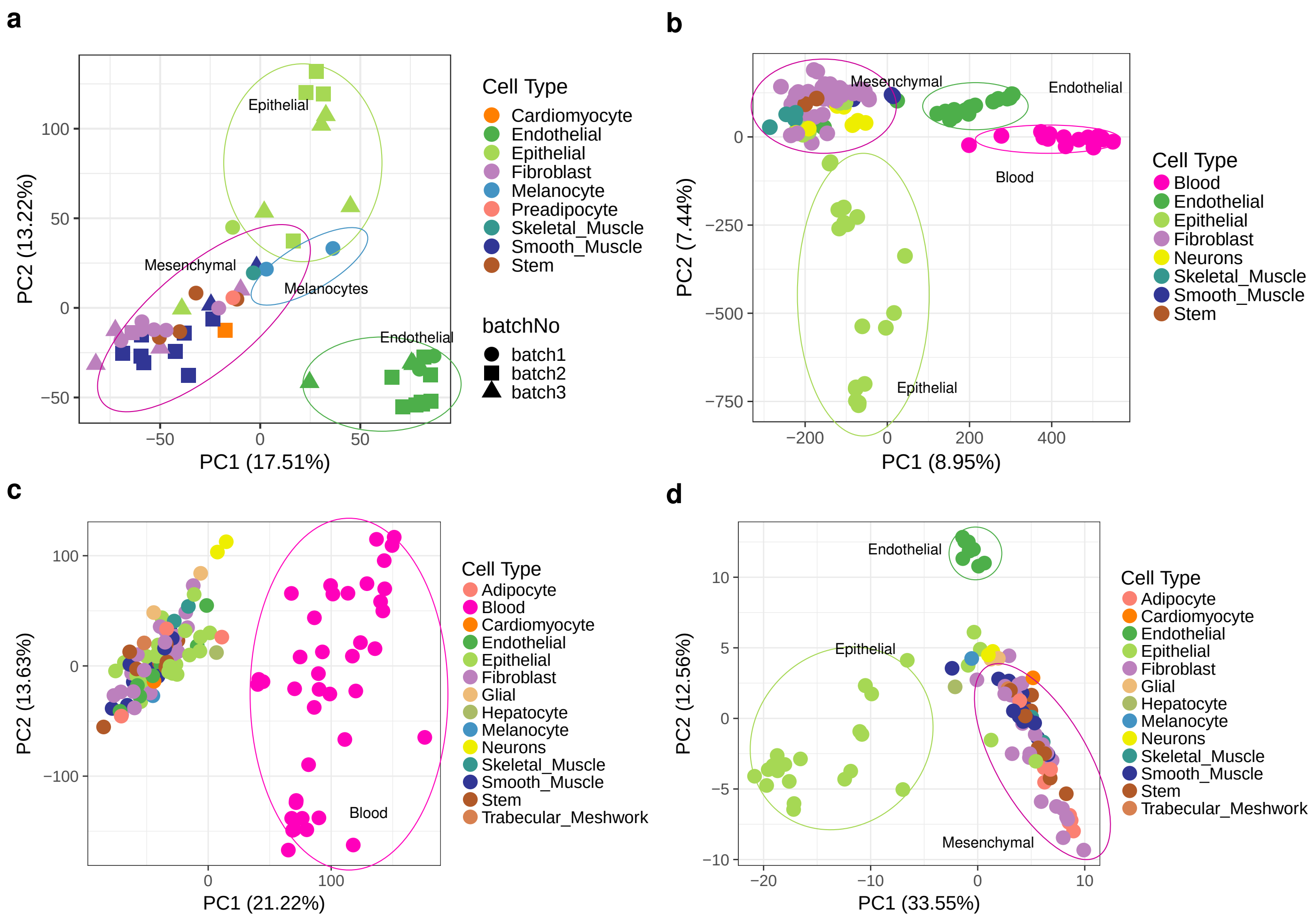

**Fig. S2.** **a)** Silhouette widths (x axis) for 3, 4 and 5 clusters based on the hierarchical clustering in Fig. 1b. Silhouette provides a succinct graphical representation of how well each object lies within its cluster. The silhouette ranges from -1 to +1, where a high value indicates that the object is well matched to its own cluster and poorly matched to neighboring clusters. The average width for 3, 4 and 5 clusters is 0.825, 0.850 and 0.848, respectively, supporting the four groups that we identify from the clustering (highest scoring). The silhouette provides also support for the epithelial sub-clustering. **b)** Elbow plot of the total within group sum of squares (y axis) with respect to the number of clusters (x axis). The clusters are obtained with hierarchical clustering using $1 - cor(X)$ as the distance criterion, where $X$ is the matrix of Pearson's correlation between ENCODE primary cells (Fig. 1b). This elbow plot suggests an optimal number of clusters of 4, after which the graph stabilizes. **c)** Estimation of proportion of variance explained by the factors body location, germ layer and major cell type. **d)** Distance (y axis) for each ENCODE primary cell line (x axis) from their respective cluster centroid based on the hierarchical clustering in Fig. 1b. Each colour represents one body location. Distance obtained as $1 - cor$, with $cor$ the Pearson's correlation between the primary cell line and the centroid. The average distances to the cluster centroids are similar across different tissues–comparatively much smaller than the distances to the centroids from other clusters (off-cluster mean). The distances are only slightly larger for kidney samples, consistent with samples from the kidney driving the sub-clustering of the epithelial samples **e)** Correlation coefficient between each pair of cell lines within each body location and for all samples. On average the correlation of gene expression between any pair of lung samples is comparable to that between any pair of samples taking anywhere within the human body.

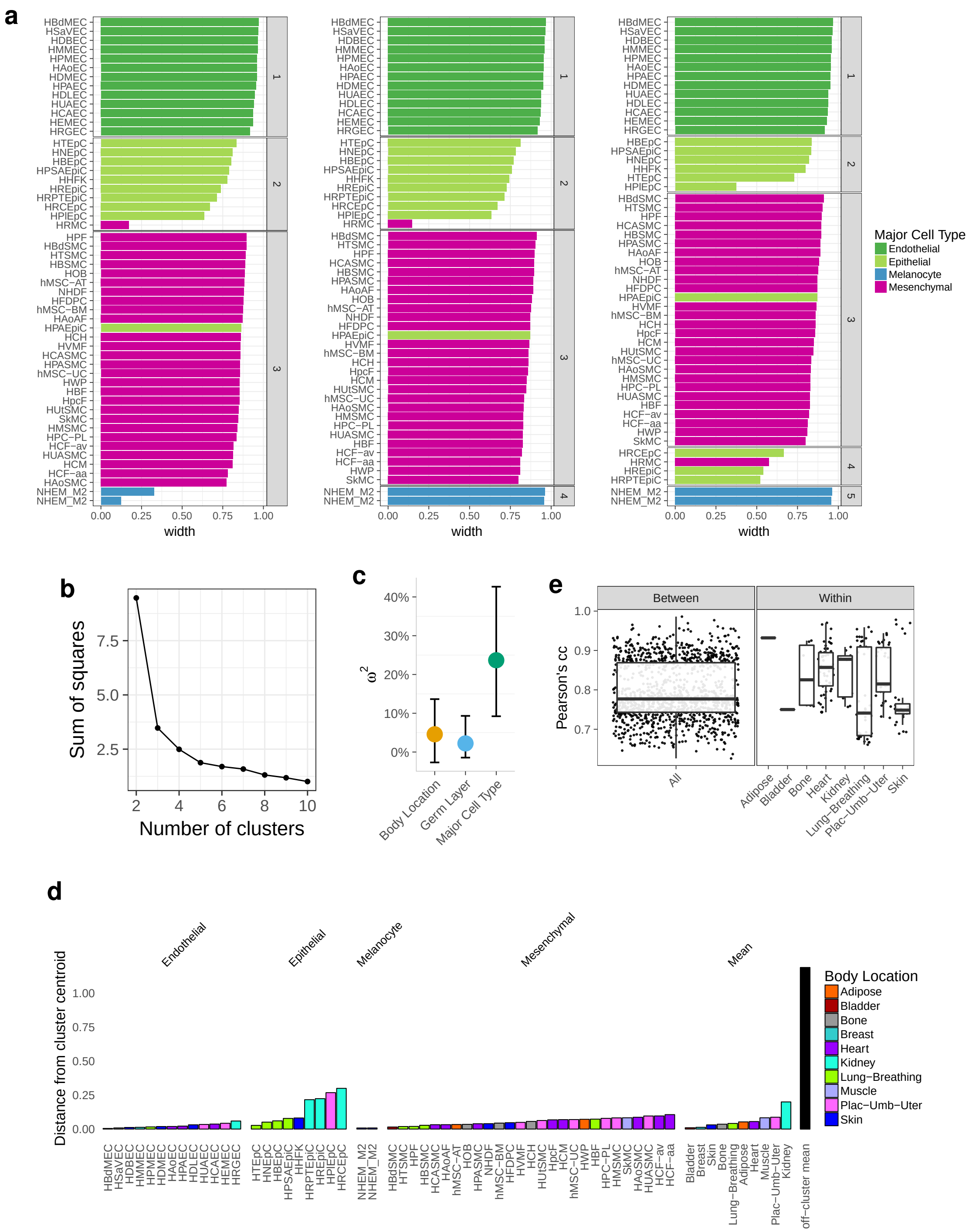

**Fig. S3**. **a)** Hierarchical clustering of primary cells profiled by CAGE. **b)** Hierarchical clustering of primary cells profiled by CAGE, except blood cells. This is based on 209 most variable genes across all samples, excluding blood cells, selected by the projection score. **c)** Hierarchical clustering of primary cells profiled by DNAse-seq.

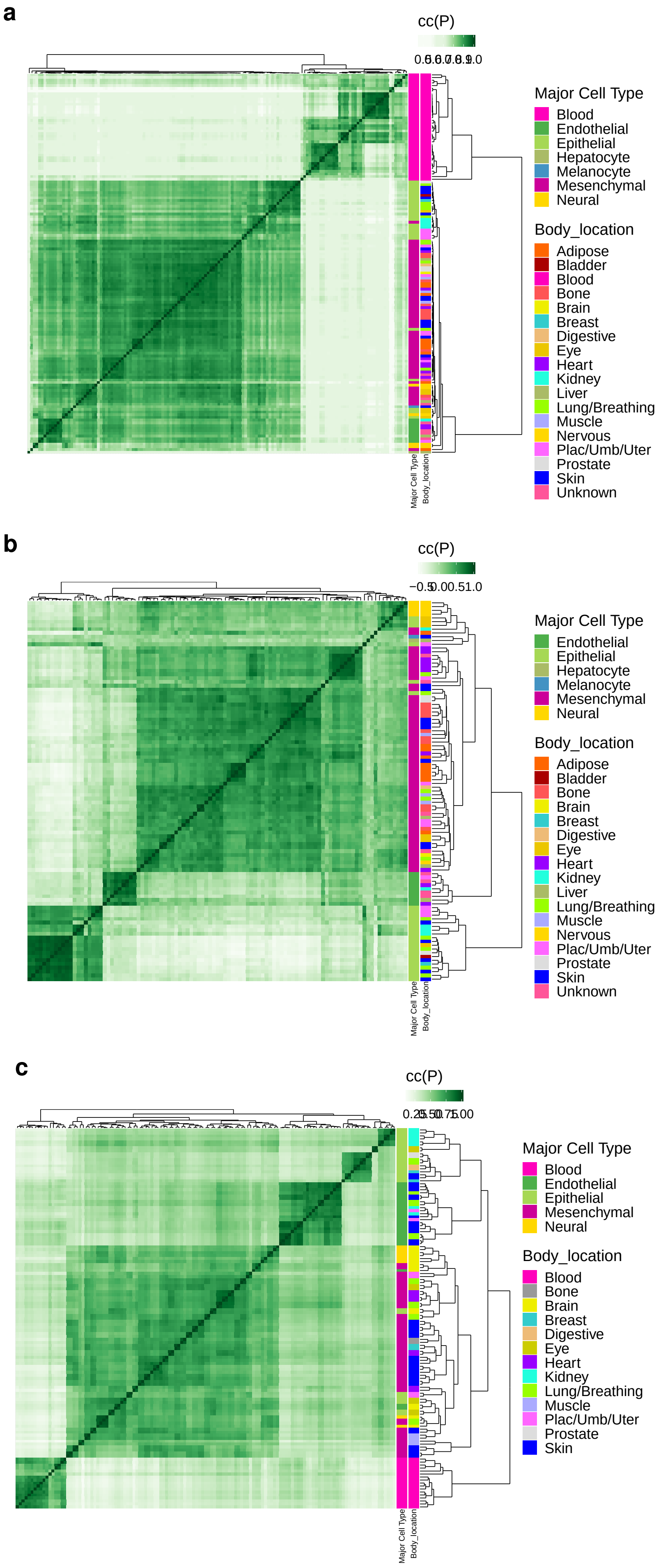

**Fig. S4**. tSNE of primary cells profiled by CAGE, RNA-seq and DNAse-seq.

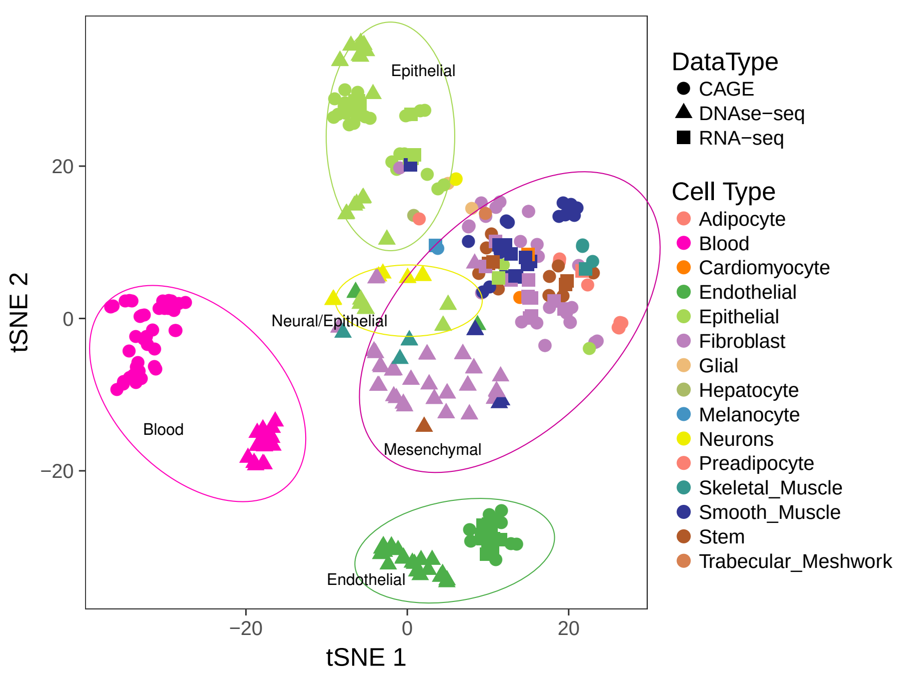

**Fig. S5**. **a)** Expression of 333 neural specific genes in FANTOM data. **b)** GO enrichment of neural specific genes.

**a**

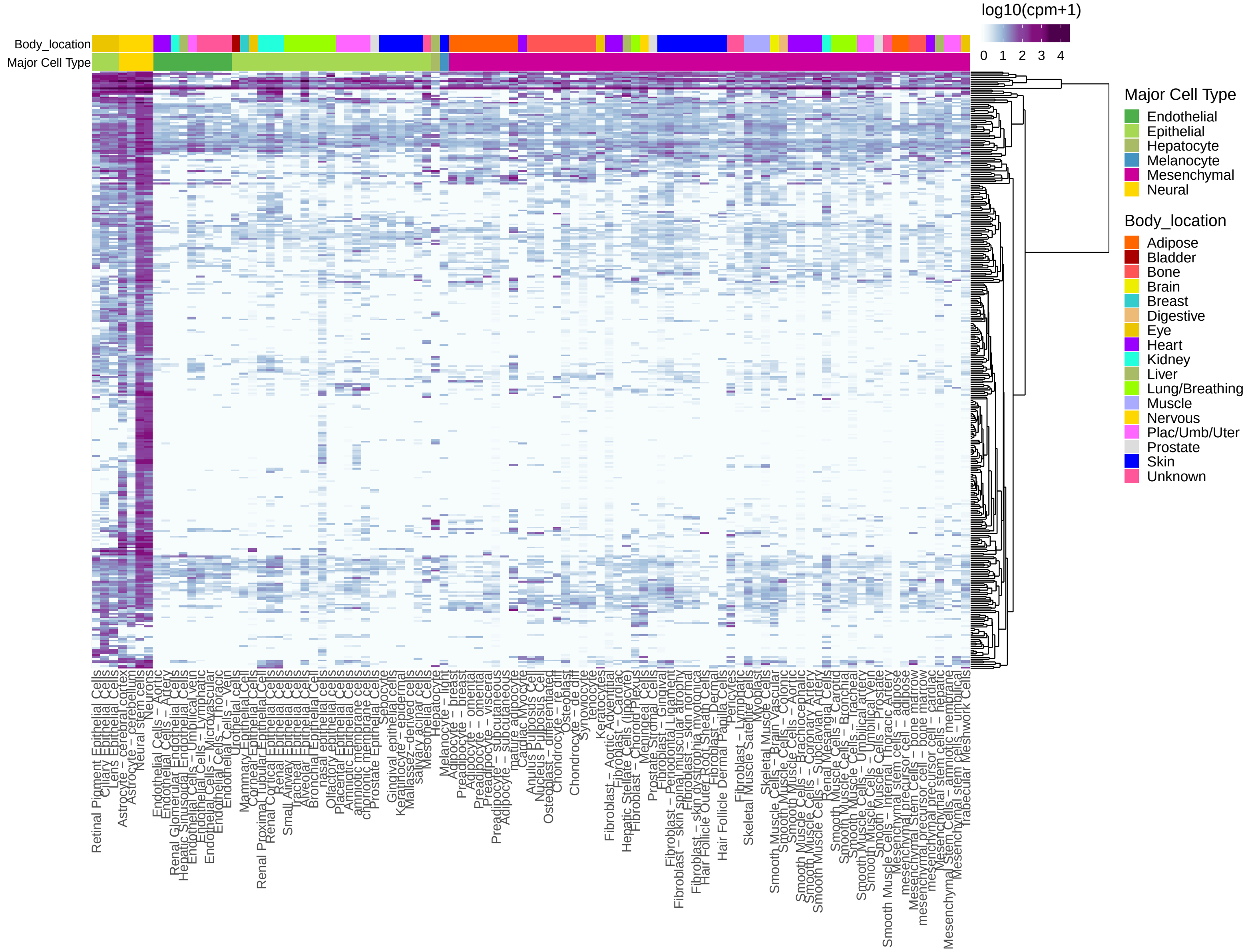


**b**

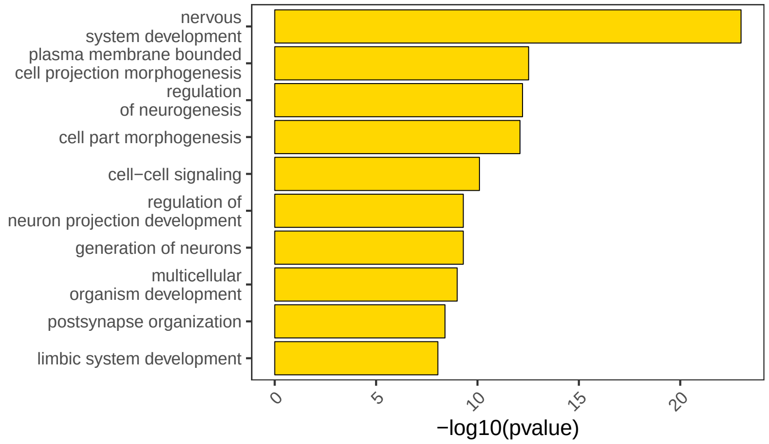

**Fig. S6**. Expression of endothelial, epithelial, mesenchymal and neural specific genes, across different experiments. "Array" data includes microarray data for which DNAse-seq data was available [10], "brainRNA-seq" data includes data from brain bulk cell populations [52], "RNA-seq" refers to ENCODE data of primary cells as well as induced neurons, and ARPE-19 retinal pigmented epithelial cells and HFF fibroblasts from GEO series GSE120891 [50], "scRNA-seq" data includes single cell RNA-seq experiments on adult brain [51]. For all RNA-seq data, genes are quantified as log10(RPKM+0.01), while microarray data is quantile normalized to match the distribution of ENCODE "RNA-seq" genes. The expression in single cell data is shown as the average expression across all the cells of a given type. The samples are clustered with complete linkage and distance based on Pearson's correlation coefficient. Genes are sorted by cell specificity and maximum expression in the related cells. Gray cells represent genes with missing values in microarray data. "DE Cell Type" refers to the cell type specificity of the genes (rows).

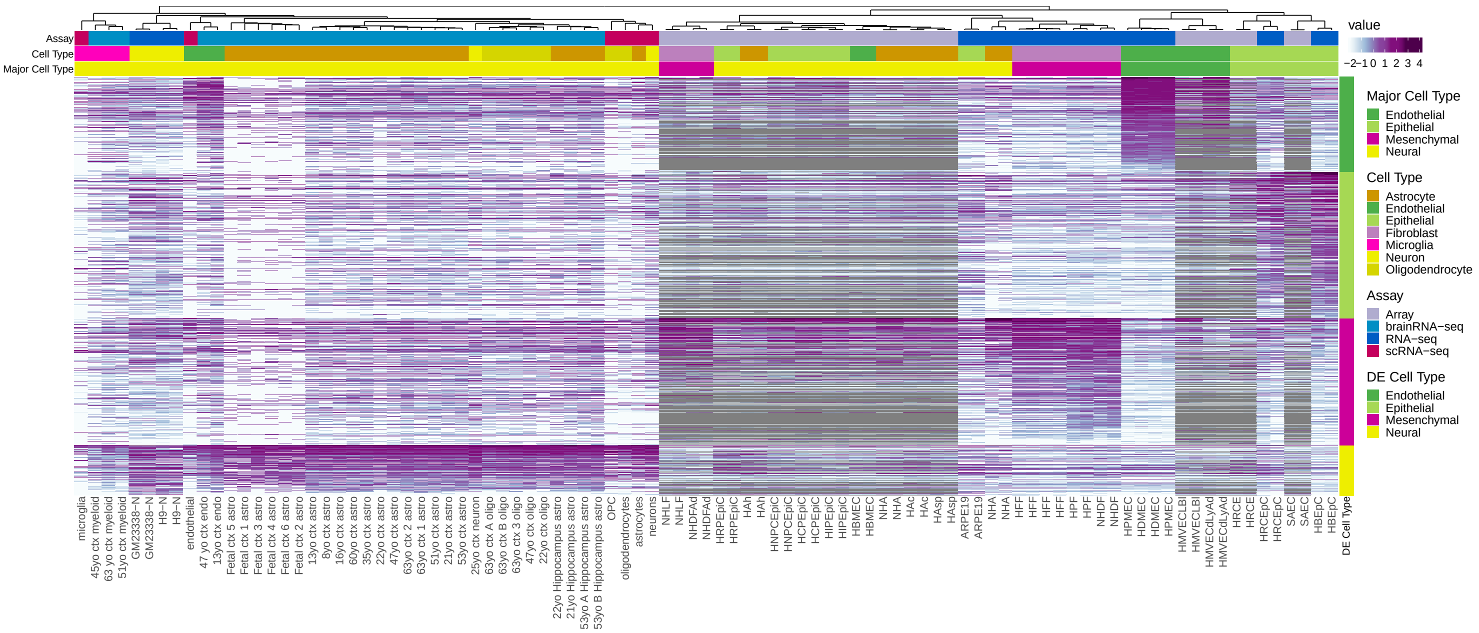

**Fig. S7**. Expression of neural genes as log10(RPKM+0.1), specific of each neural subtypes, identified from single cell RNA-seq data [51]. The expression in single cell RNA-seq data is consistent with the expression in the experiments on bulk cell populations [52]. The samples are clustered with complete linkage and distance based on Pearson's correlation coefficient. Genes are sorted by cell specificity and mean e The samples are clustered with complete linkage and distance based on Pearson's correlation coefficient. Genes are sorted by cell specificity and mean expression in the related cells. "DE Cell Type" refers to the cell type specificity of the genes (rows).

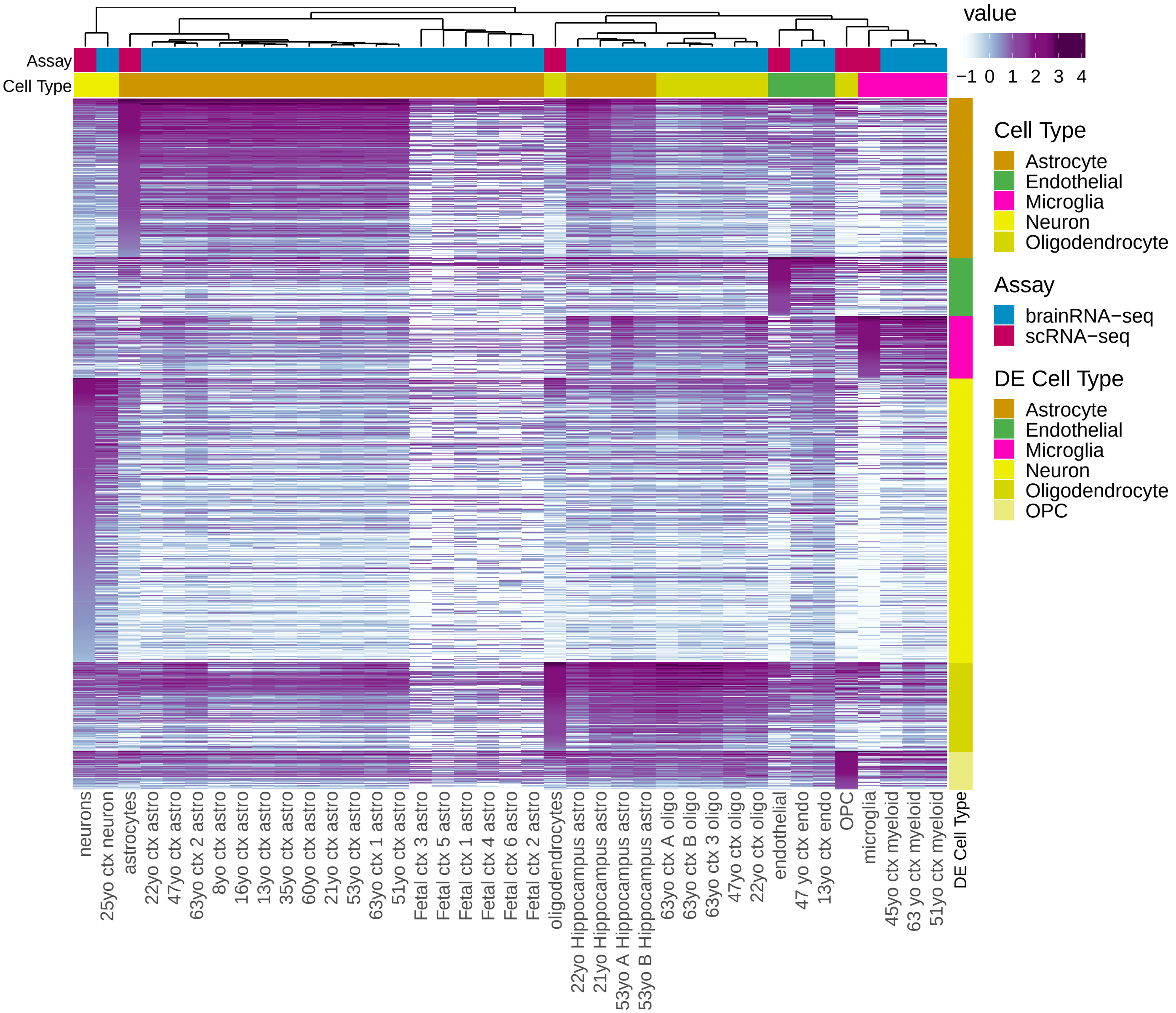

**Fig. S8**. Major cell types in the *Tabula Muris* single-cell transcriptomic data from microfluidic droplet-based 3'-end counting with the 10x Genomics platform. **a)** PCA of 53,617 cells, colored by their assigned major cell type. Cells with specialized functions were not assigned a major cell type category and are depicted in grey under the "TS" (tissue-specific) category. Most single cells clustered by major cell type, with epithelial cells exhibiting greater heterogeneity. Hepatocytes, in particular, albeit closer to epithelial cells than to cells from other major types, clustered separately (shown in yellow in the PCA). **b)** PCA excluding the 1,764 hepatocyte cells. **c)** Spearman's correlation computed between the mean expression profiles of each cell type tissue pair, using the variable genes identified by *Seurat* and the set of cells in (b). Hierarchical clustering was computed using Ward2's criterion with euclidean distance. Nearly all cell types clustered within the corresponding major cell types, with the exception being cardiac muscle cells which we labelled as mesenchymal, but cluster within the epithelial type.

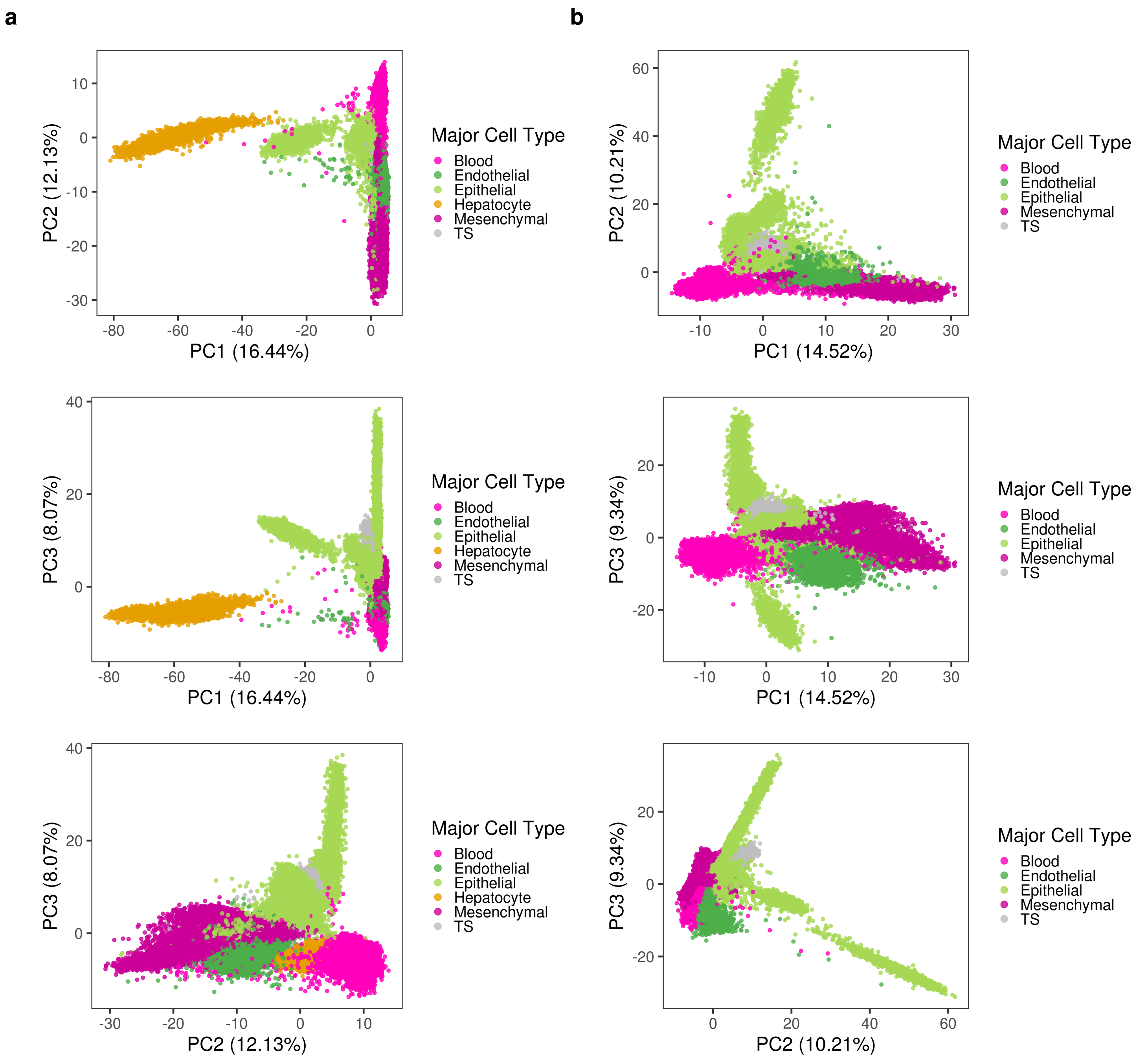

c

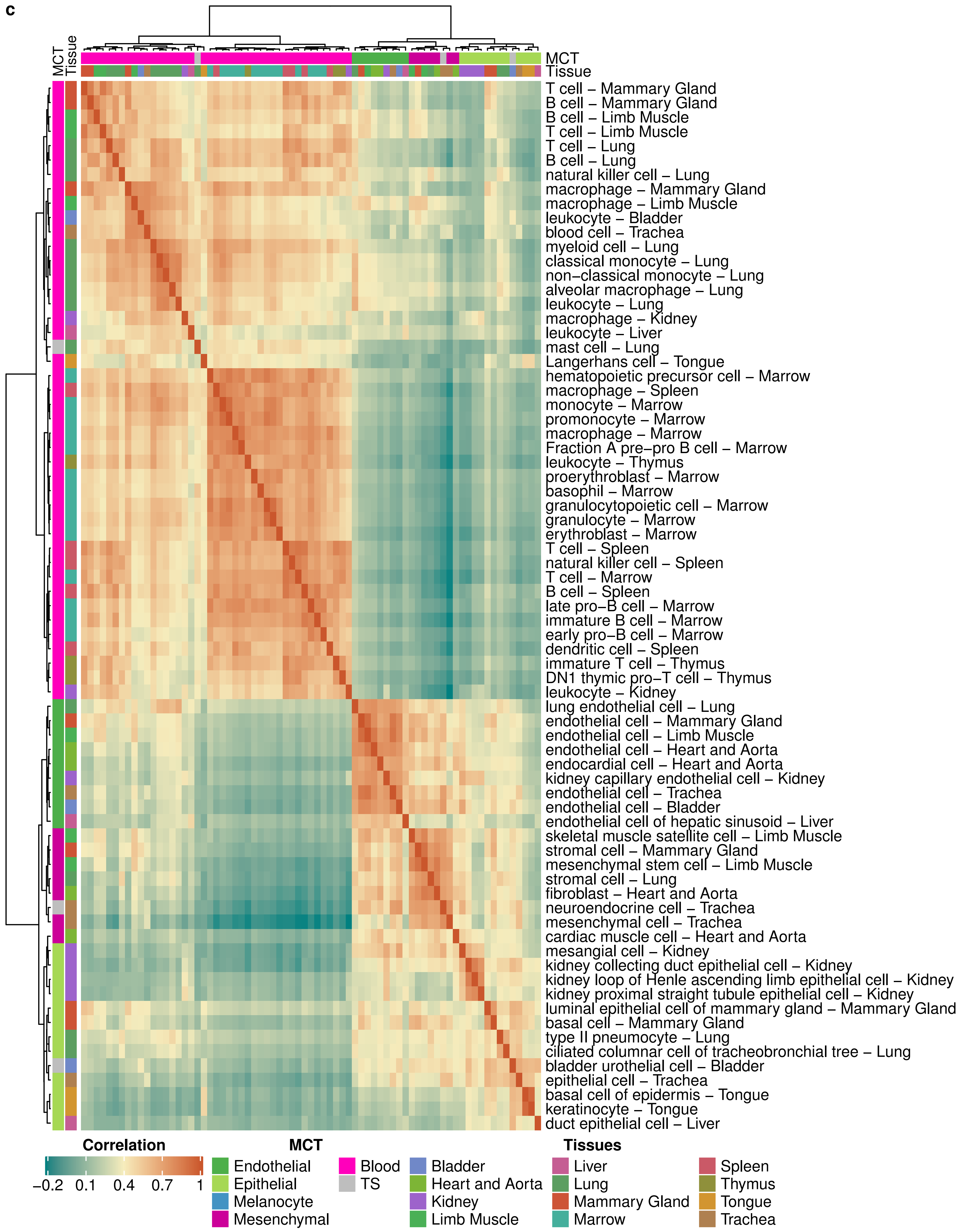
MCT
Tissue
T cell – Mammary Gland
B cell – Mammary Gland
B cell – Limb Muscle
T cell – Limb Muscle
T cell – Lung
B cell – Lung
natural killer cell – Lung
macrophage – Mammary Gland
macrophage – Limb Muscle
leukocyte – Bladder
blood cell – Trachea
myeloid cell – Lung
classical monocyte – Lung
non–classical monocyte – Lung
alveolar macrophage – Lung
leukocyte – Lung
macrophage – Kidney
leukocyte – Liver
mast cell – Lung
Langerhans cell – Tongue
hematopoietic precursor cell – Marrow
macrophage – Spleen
monocyte – Marrow
promonocyte – Marrow
macrophage – Marrow
Fraction A pre–pro B cell – Marrow
leukocyte – Thymus
proerythroblast – Marrow
basophil – Marrow
granulocytopoietic cell – Marrow
granulocyte – Marrow
erythroblast – Marrow
T cell – Spleen
natural killer cell – Spleen
T cell – Marrow
B cell – Spleen
late pro–B cell – Marrow
immature B cell – Marrow
early pro–B cell – Marrow
dendritic cell – Spleen
immature T cell – Thymus
DN1 thymic pro–T cell – Thymus
leukocyte – Kidney
lung endothelial cell – Lung
endothelial cell – Mammary Gland
endothelial cell – Limb Muscle
endothelial cell – Heart and Aorta
endocardial cell – Heart and Aorta
kidney capillary endothelial cell – Kidney
endothelial cell – Trachea
endothelial cell – Bladder
endothelial cell of hepatic sinusoid – Liver
skeletal muscle satellite cell – Limb Muscle
stromal cell – Mammary Gland
mesenchymal stem cell – Limb Muscle
stromal cell – Lung
fibroblast – Heart and Aorta
neuroendocrine cell – Trachea
mesenchymal cell – Trachea
cardiac muscle cell – Heart and Aorta
mesangial cell – Kidney
kidney collecting duct epithelial cell – Kidney
kidney loop of Henle ascending limb epithelial cell – Kidney
kidney proximal straight tubule epithelial cell – Kidney
luminal epithelial cell of mammary gland – Mammary Gland
basal cell – Mammary Gland
type II pneumocyte – Lung
ciliated columnar cell of tracheobronchial tree – Lung
bladder urothelial cell – Bladder
epithelial cell – Trachea
basal cell of epidermis – Tongue
keratinocyte – Tongue
duct epithelial cell – Liver
Correlation
−0.2
0.1
0.4
0.7
1
MCT
Endothelial
Epithelial
Melanocyte
Mesenchymal
Blood
TS
Tissues
Bladder
Heart and Aorta
Kidney
Limb Muscle
Liver
Lung
Mammary Gland
Marrow
Spleen
Thymus
Tongue
Trachea

**Fig. S9**. Major cell types in the *Tabula Muris* single-cell transcriptomic data from FACS-based full length transcript analysis using SMART-Seq2 libraries. **a)** PCA of 44,084 cells, colored by their assigned major cell type. Cells with specialized functions were not assigned a major cell type category and are depicted in grey under the "TS" (tissue-specific) category. As with the 10x data, most single cells clustered by major cell type, with hepatocytes, while closer to epithelial cells, clustered quite distinctively as well. **b)** PCA excluding the 391 hepatocyte cells. **c)** Spearman's correlation computed between the mean expression profiles of each cell-type tissue pair, using the variable genes identified by *Seurat* and the set of cells used in (b). Hierarchical clustering was computed using Ward2's criterion with euclidean distance. As with the 10x data, most cells properly clustered within the expected major cell types. Indeed, blood, mesenchymal, endothelial and epithelial cells clustered mostly within their major cell types. Neural cells clustered together with different types of pancreatic endocrine cells, consistent with the strong morphological, and physiological similarities between these two types of cells[71], and with a few epithelial cells. This resembles the neuroepithelial cluster that we found in human primary cells. The exception is oligodendrocytes, that we labelled as neural, but that unexpectedly cluster within the epithelial type. Finally, a very few cell types that we labelled as mesenchymal and epithelial clustered together separated from the rest, but connected to the epithelial cluster.

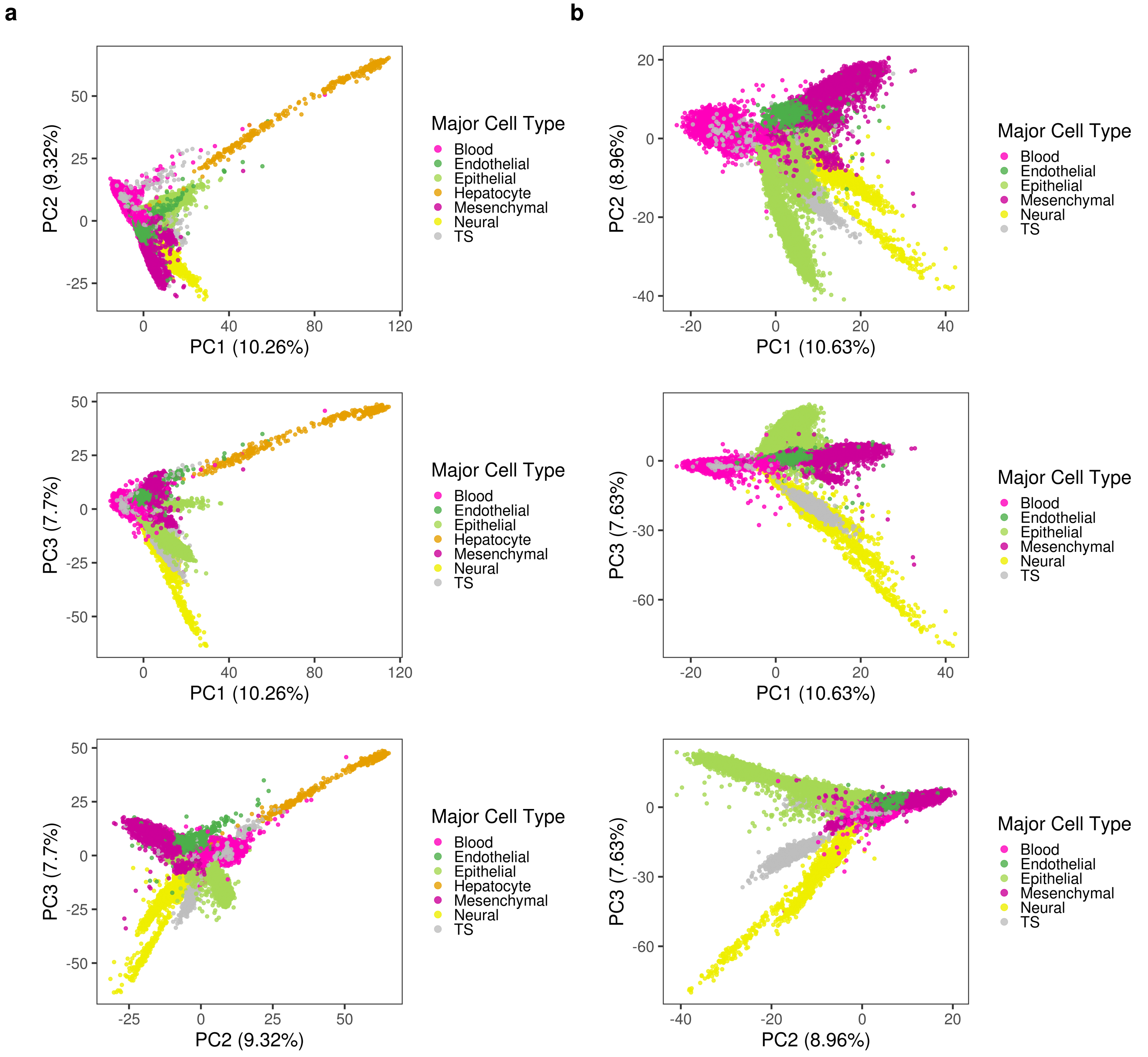

c

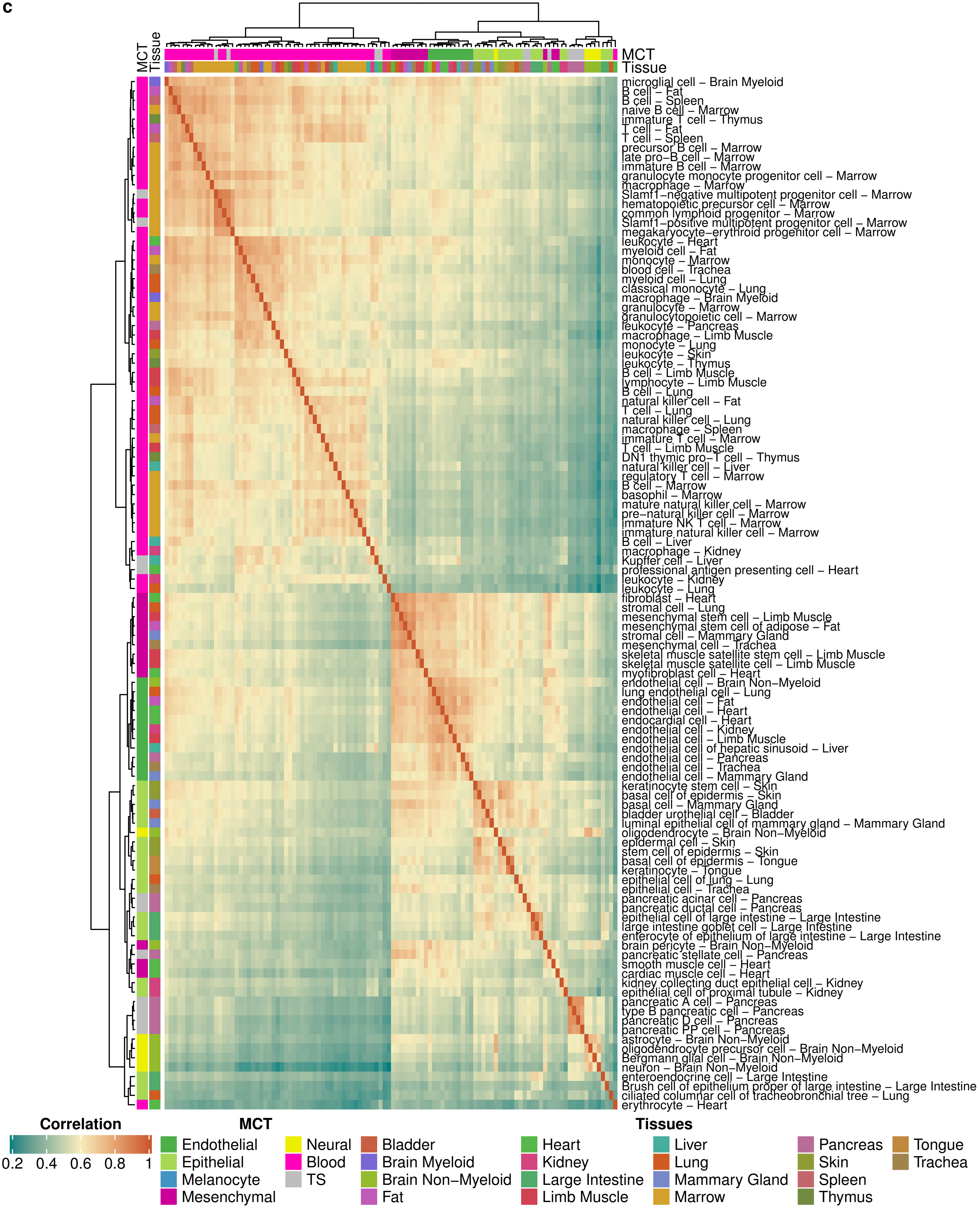
MCT
Tissue
microglial cell – Brain Myeloid
B cell – Fat
B cell – Spleen
naive B cell – Marrow
immature T cell – Thymus
T cell – Fat
T cell – Spleen
precursor B cell – Marrow
late pro−B cell – Marrow
immature B cell – Marrow
granulocyte monocyte progenitor cell – Marrow
macrophage – Marrow
Slamf1−negative multipotent progenitor cell – Marrow
hematopoietic precursor cell – Marrow
common lymphoid progenitor – Marrow
Slamf1−positive multipotent progenitor cell – Marrow
megakaryocyte−erythroid progenitor cell – Marrow
leukocyte – Heart
myeloid cell – Fat
monocyte – Marrow
blood cell – Trachea
myeloid cell – Lung
classical monocyte – Lung
macrophage – Brain Myeloid
granulocyte – Marrow
granulocytopoietic cell – Marrow
leukocyte – Pancreas
macrophage – Limb Muscle
monocyte – Lung
leukocyte – Skin
leukocyte – Thymus
B cell – Limb Muscle
lymphocyte – Limb Muscle
B cell – Lung
natural killer cell – Fat
T cell – Lung
natural killer cell – Lung
macrophage – Spleen
immature T cell – Marrow
T cell – Limb Muscle
DN1 thymic pro−T cell – Thymus
natural killer cell – Liver
regulatory T cell – Marrow
B cell – Marrow
basophil – Marrow
mature natural killer cell – Marrow
pre−natural killer cell – Marrow
immature NK T cell – Marrow
immature natural killer cell – Marrow
B cell – Liver
macrophage – Kidney
Kupffer cell – Liver
professional antigen presenting cell – Heart
leukocyte – Kidney
leukocyte – Lung
fibroblast – Heart
stromal cell – Lung
mesenchymal stem cell – Limb Muscle
mesenchymal stem cell of adipose – Fat
stromal cell – Mammary Gland
mesenchymal cell – Trachea
skeletal muscle satellite stem cell – Limb Muscle
skeletal muscle satellite cell – Limb Muscle
myofibroblast cell – Heart
endothelial cell – Brain Non−Myeloid
lung endothelial cell – Lung
endothelial cell – Fat
endothelial cell – Heart
endocardial cell – Heart
endothelial cell – Kidney
endothelial cell – Limb Muscle
endothelial cell of hepatic sinusoid – Liver
endothelial cell – Pancreas
endothelial cell – Trachea
endothelial cell – Mammary Gland
keratinocyte stem cell – Skin
basal cell of epidermis – Skin
basal cell – Mammary Gland
bladder urothelial cell – Bladder
luminal epithelial cell of mammary gland – Mammary Gland
oligodendrocyte – Brain Non−Myeloid
epidermal cell – Skin
stem cell of epidermis – Skin
basal cell of epidermis – Tongue
keratinocyte – Tongue
epithelial cell of lung – Lung
epithelial cell – Trachea
pancreatic acinar cell – Pancreas
pancreatic ductal cell – Pancreas
epithelial cell of large intestine – Large Intestine
large intestine goblet cell – Large Intestine
enterocyte of epithelium of large intestine – Large Intestine
brain pericyte – Brain Non−Myeloid
pancreatic stellate cell – Pancreas
smooth muscle cell – Heart
cardiac muscle cell – Heart
kidney collecting duct epithelial cell – Kidney
epithelial cell of proximal tubule – Kidney
pancreatic A cell – Pancreas
type B pancreatic cell – Pancreas
pancreatic D cell – Pancreas
pancreatic PP cell – Pancreas
astrocyte – Brain Non−Myeloid
oligodendrocyte precursor cell – Brain Non−Myeloid
Bergmann glial cell – Brain Non−Myeloid
neuron – Brain Non−Myeloid
enteroendocrine cell – Large Intestine
Brush cell of epithelium proper of large intestine – Large Intestine
ciliated columnar cell of tracheobronchial tree – Lung
erythrocyte – Heart
Correlation
0.2
0.4
0.6
0.8
1
MCT
Endothelial
Epithelial
Melanocyte
Mesenchymal
Neural
Blood
TS
Tissues
Bladder
Brain Myeloid
Brain Non−Myeloid
Fat
Heart
Kidney
Large Intestine
Limb Muscle
Liver
Lung
Mammary Gland
Marrow
Pancreas
Skin
Spleen
Thymus
Tongue
Trachea

**Fig. S10**. Boxplot of the pairwise Pearson's correlation coefficient of each pair of FANTOM primary cell types of the same major cell type. Transcriptional diversity is similar among cell types.

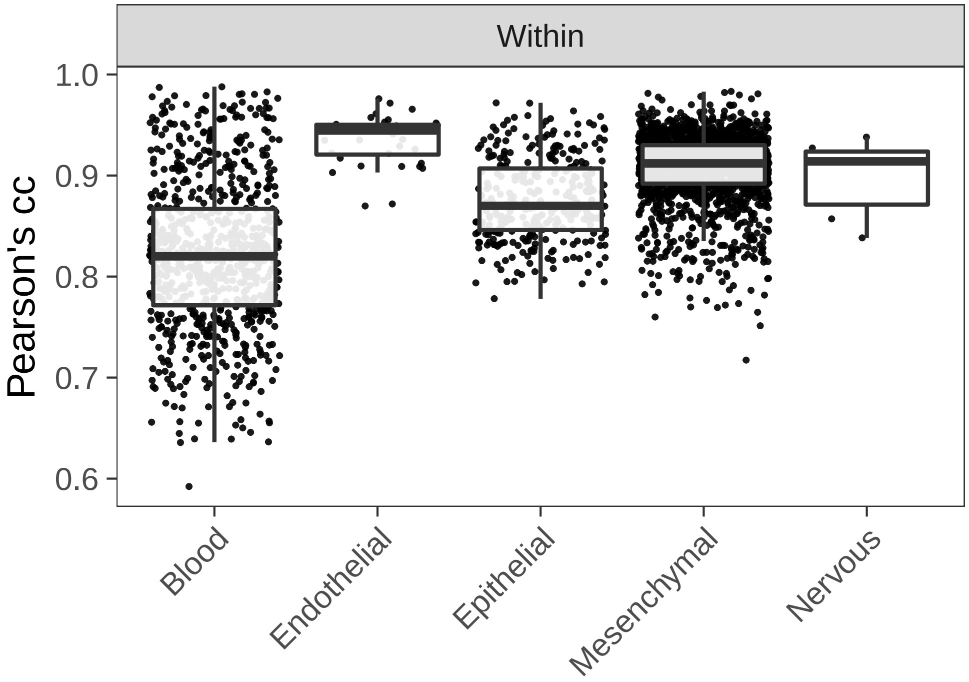

**Fig. S11**. **a)** Number of cell type specific genes for different cutoffs of FDR (rows) and log2-fold-change (columns). The most stringent combination of 0.01 FDR and 2 log2-fold-change was chosen (black square). The number of cell type specific genes is quite robust to FDR cutoff. **b)** Distribution of 2,873 cell type specific genes by cell type and gene biotype.

**a**

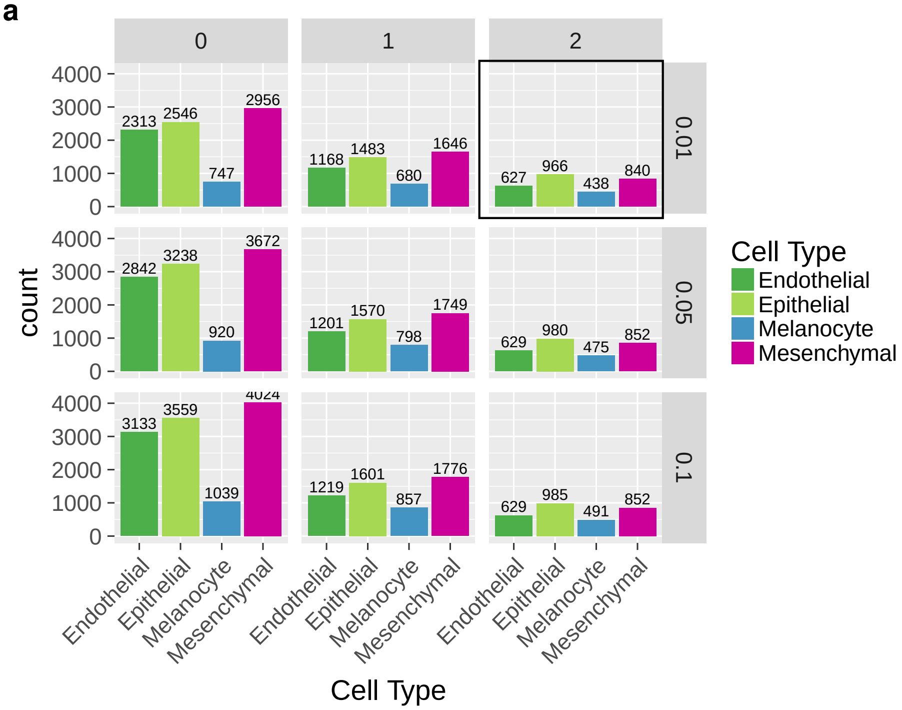


**b**

| | Endothelial | Epithelial | Melanocyte | Mesenchymal | **Total** |
|---|---|---|---|---|---|
| lncRNA | 67 | 59 | 46 | 153 | 325 |
| protein coding | 531 | 857 | 294 | 729 | 2,411 |
| pseudogene | 37 | 34 | 13 | 53 | 137 |
| **Total** | 635 | 950 | 353 | 935 | 2,873 |

**Fig. S12. a)** PCA showing the top correlated genes with the major cell types in the principal component space. This corresponds to Figure S1a. PC coordinates are normalized to the interval [-1,1]. Correlation is computed as cosine between the centroid vectors of each major cell type cluster and the expression vector of each gene in normalized principal component coordinates. Genes with cosine $>$ 0.9 are shown as grey arrows. Selected genes with literature support for their cell type specific function are colored according to the closest major cell type cluster (Table S8). **b)** Expression values of genes with cosine $>$ 0.9, mean-centered and scaled across samples.

**a**

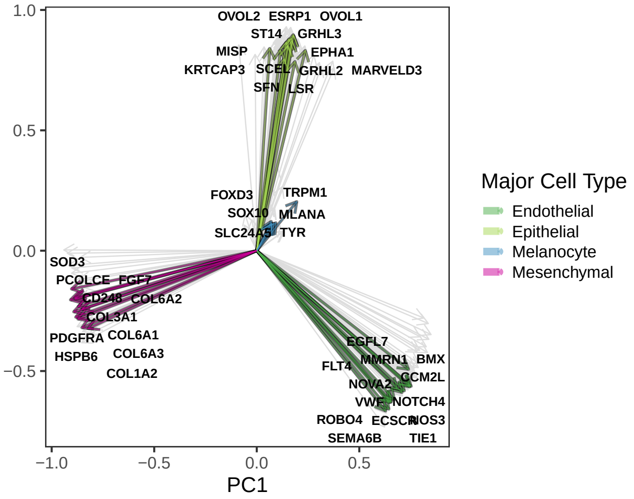


**b**

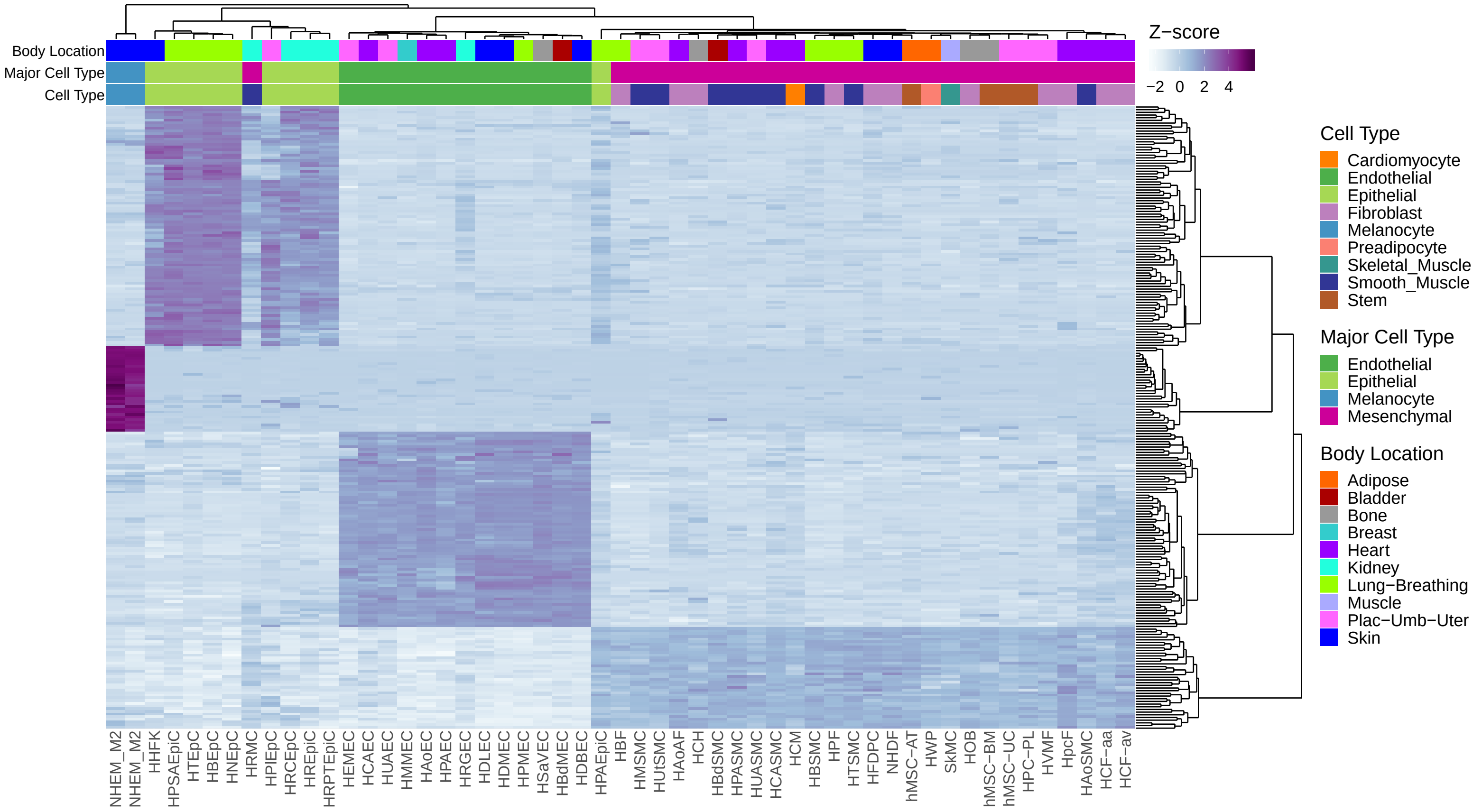

**Fig. S13**. **a)** GO term enrichments for the genes with the highest expression correlation with lncRNA RP11-536O18.1 (Pearson's correlation coefficient $> 0.9$). **b)** GO term enrichments for cell type specific genes. Only the 10 most significant terms for each cell type are shown.

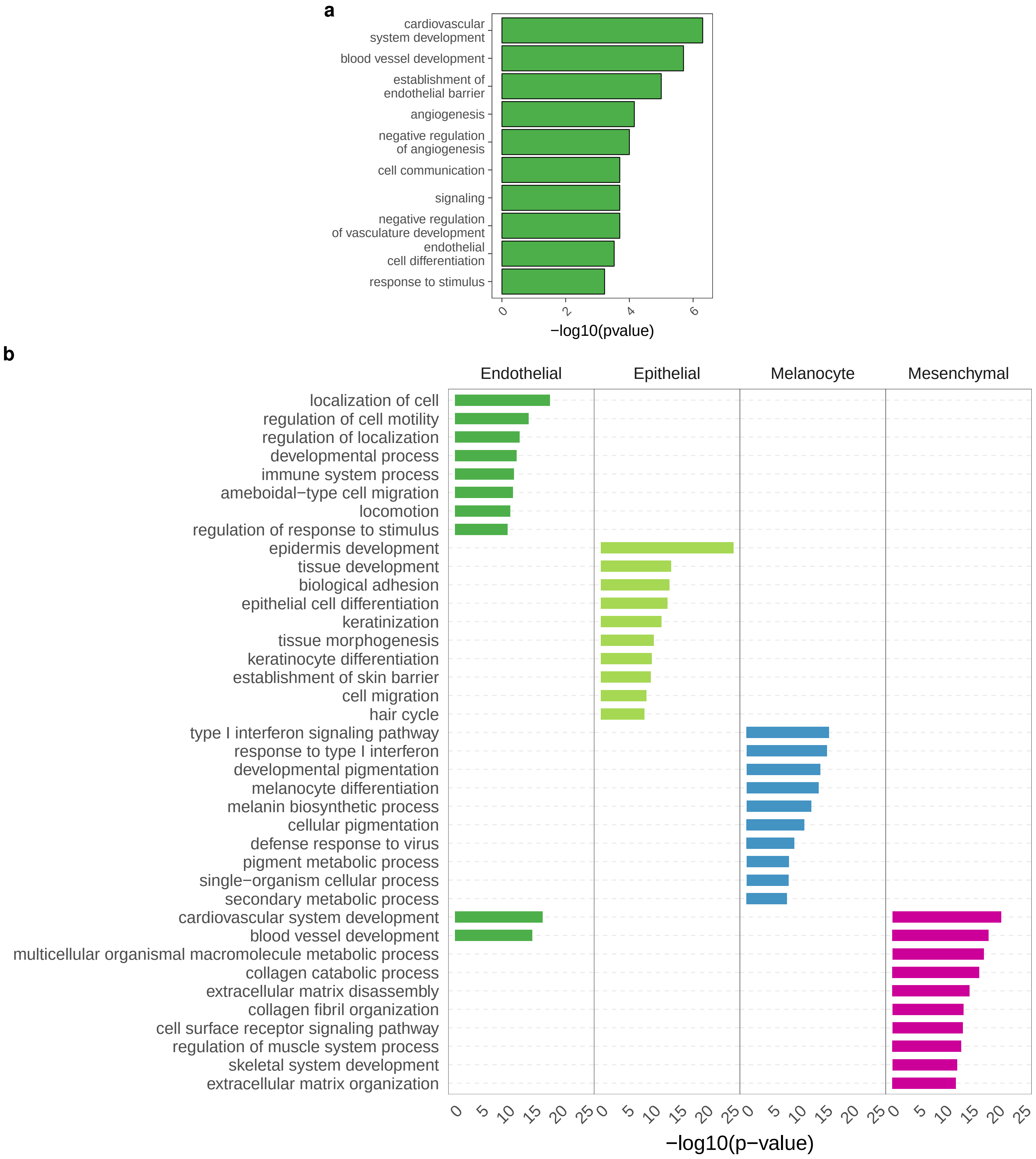

**Fig. S14.** Expression of cell-type specific genes (columns, 2,581 genes) in FANTOM5 samples (rows). The color bar on top of the columns indicates the cell type specificity defined from the ENCODE RNA-seq samples.

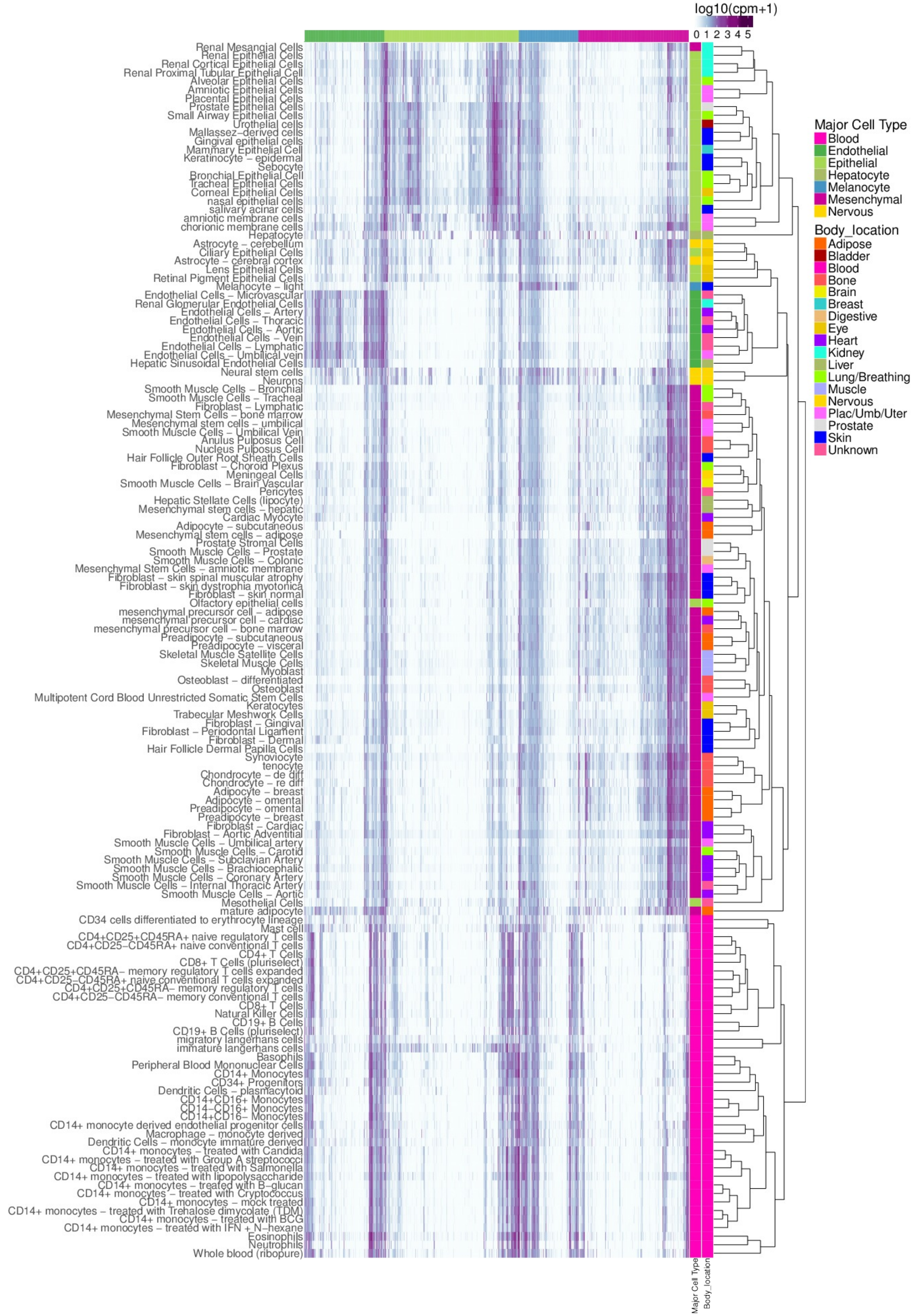

**Fig. S15.** Proportion of cell type specific genes with DNAse Hypersenstive Sites (DHSs) in their promoters (TSS - 10kb/+5kb) in each primary cell. Different marks correspond to different replicate DNAse-seq experiments in the same primary cell. For instance in HDBEC, an endothelial primary cell, between 86% and 90% of endothelial specific genes (depending on the replicate) host at least a DHS in their promoters, compared to only between 60% and 70% of epithelial specific genes. Conversely, in HPrEpiC, an epithelial primary cell, 88% of epithelial specific genes host DHSs in their promoters, compared to 74% of the endothelial specific genes.

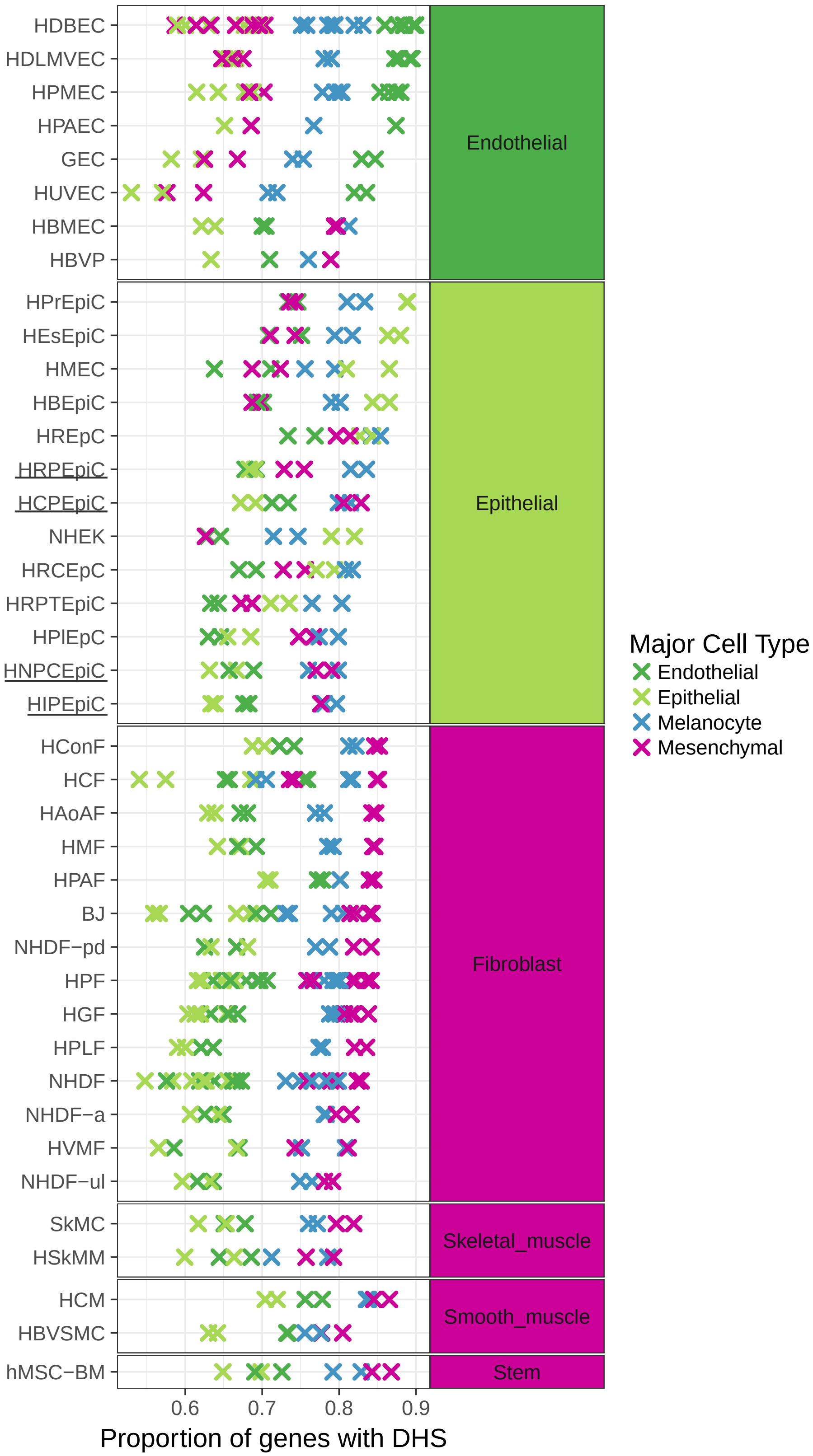

**Fig. S16**. **a)** Average profiles of H3K27ac and H3K4me3 marks at the transcription start site (TSS) +/- 2000 bp for different sets of cell type specific genes. Each line is the aggregate profile for a given cell type. The signal is mean-centered and scaled within each experiment to normalize across samples. Genes specific of a given cell type show higher signal for activating chromatin marks in the respective cell types compared to other cell types. **b)** Aggregate profiles of chromatin marks for each set of cell type specific genes and a control set of genes with similar expression in each primary cell. The profiles are centered at the TSS. Solid lines are the mean signal across all cell lines, while the width of the lines is the difference between maximum and the minimum signal across cell lines.

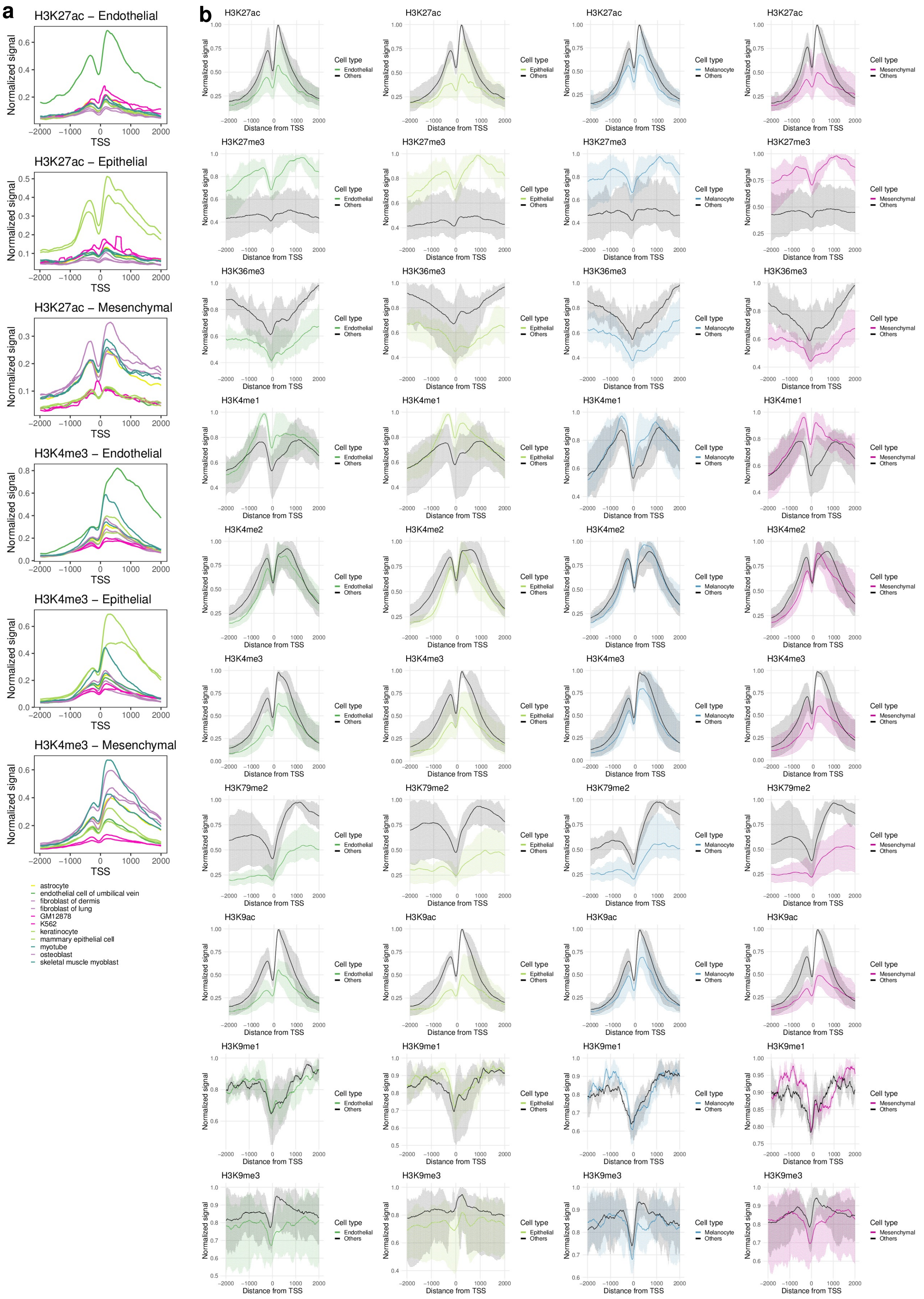

**Fig. S17**. Expression of 167 cell type specific TFs. The TFs (rows) with available motifs and with highest co-expression (Pearson's correlation >= 0.85) are highlighted with a black bar.

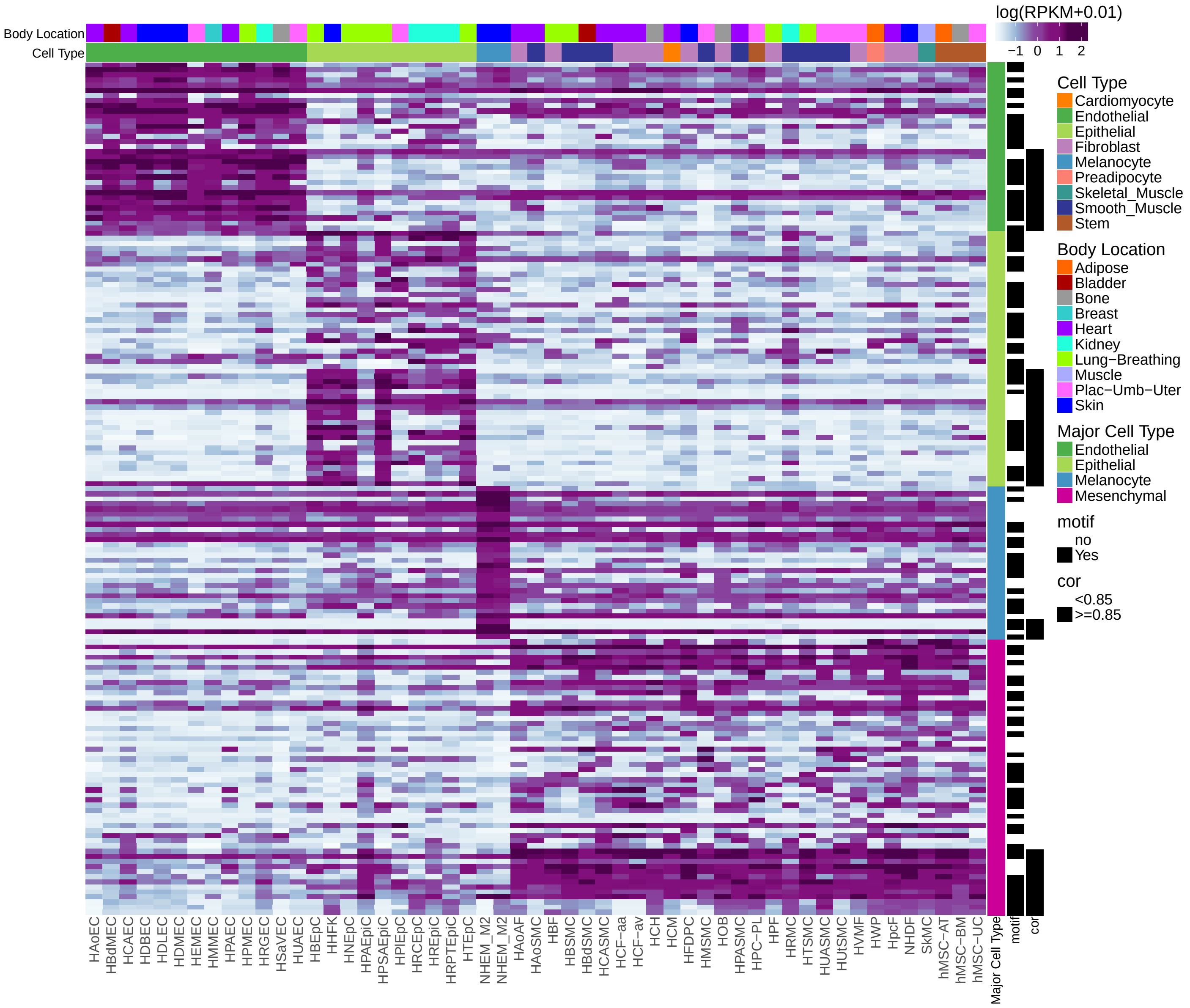

**Fig. S18.** The enrichment of cell type specific TFs in open chromatin domains of genes specific to that type is robust to the distance around the TSS. For the upstream/downstream distance pairs: -5kb/+2kb, -2kb/+1kb, -1Kb/500b, and for each TF and primary cell line we computed A: the number of genes specific from a given major cell type $i$ with motifs from TFs specific from this major cell type $i$; B: the number of genes specific from the other major cell types $j, k \neq i$ with motifs from TF specific from this type $i$; C: the total number of genes specific from $i$ with DHS minus A; D: the total number of genes specific from $j, k \neq i$ with DHS minus B; and obtained the odds ratio OR = (A/B)/(C/D). For each TF, we correlated the OR distribution for each distance pair with the OR distribution obtained for -10Kb/+5Kb, as in Fig. 2d. The Pearson's correlation coefficient is represented for each TF (y axis) and distance pair (x axis). Medians are shown in red.

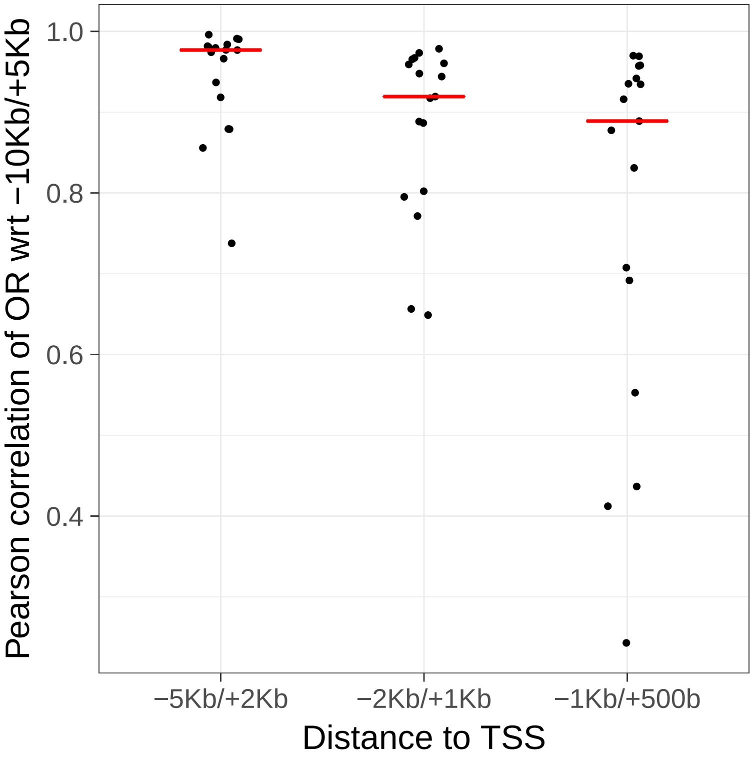

**Fig. S19**. **a)** Number of differentially expressed genes (DE, y axis) vs. number of genes with differentially spliced exons (DS, x axis), between pairs of samples of the same cell type (within, blue) or different cell types (between, red). Number of DS genes obtained using Leafcutter. As in Fig. 3b, obtained using IPSA for a given number of DS genes in a pairwise comparison of primary cells, the number of differentially expressed genes is much larger in comparisons between than within cell types. **b)** Boxplot of the number of genes with differentially spliced exons (DS, y axis) regarding the deltaPSI (differential inclusion) cutoff. The other plots are the same as **a** using different deltaPSI cutoff. Number of DS genes obtained with IPSA. The trend is robust to the deltaPSI cutoff.

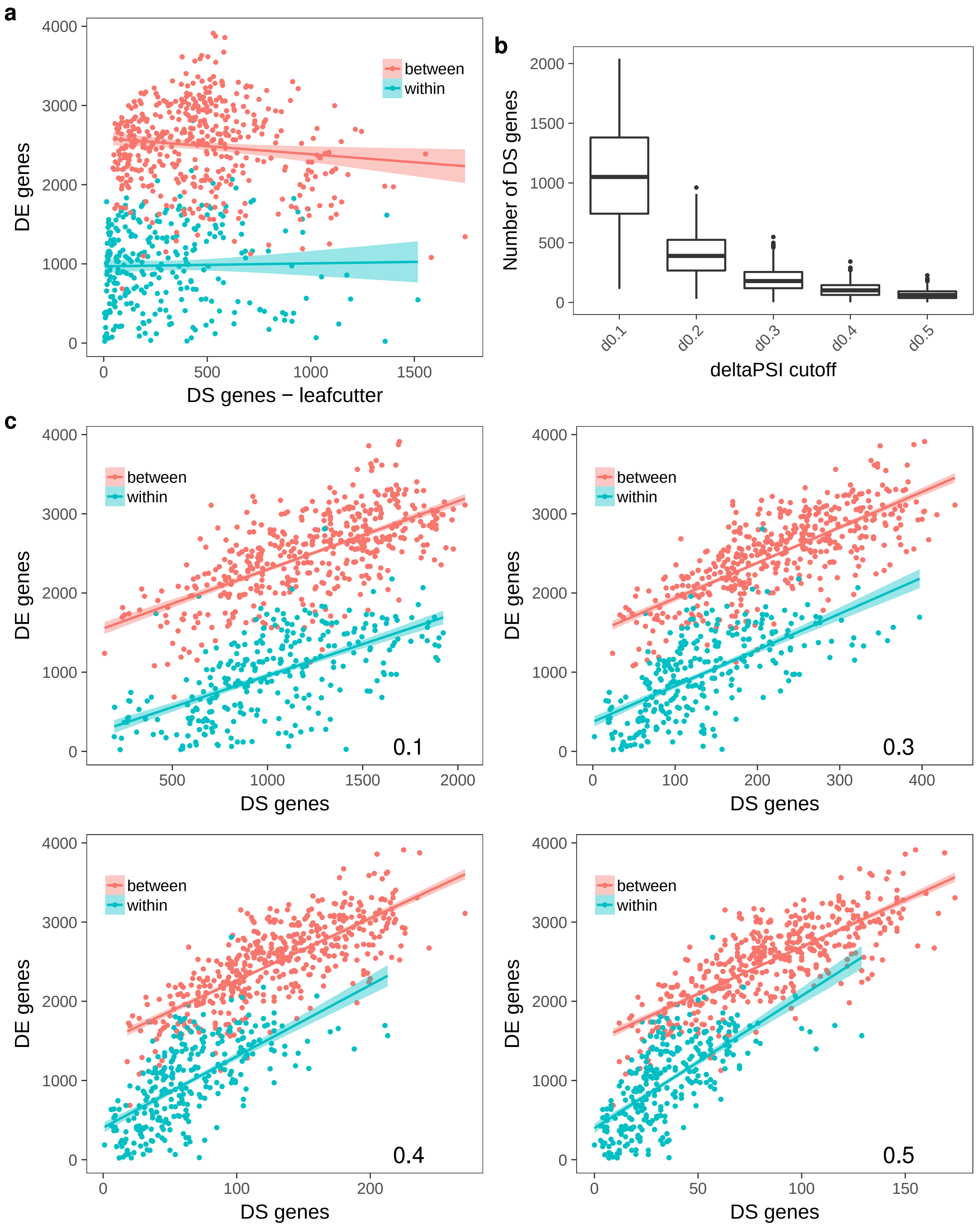

**Fig. S20**. **a**) Sashimi plot depicting differential inclusion of exon 6 of the MYL6 gene, coding for the myosin light chain 6 protein . The exon is more included in mesenchymal cells, compared to the other major cell types. The signal is the average read count for each major cell type (y axis). The average number of reads supporting each splice junctions is reported for each splice junction within each major cell type. The bottom panel shows the exonic structure of annotated transcripts in GENCODE[39]. The Sashimi plot was generated using *ggsashimi*[66]. **b**) Expression (RPKM) values for MYL6 in the ENCODE primary cell lines. **c**) Sashimi plot showing differential inclusion of exon 6 (in the middle) of the MYL6 gene in mesenchymal cells. The exon has been previously reported to be more often included in muscle cells[67]; however, we found that it is also often included in other mesenchymal cell types, including fibroblasts and stem cells, but not in adipocytes. **d**) Predicted 3-dimensional structure of MYL6 protein (SWISS-MODEL on template 3dtp.1.D). The 3 EF-hand motifs are shown in red, brown and green. Exon 6, shown in yellow, is part of the last EF-hand motif. Interestingly, the exon 6 is translated as part of the MYL6 N-terminal EF-hand domain, calcium-binding domain which mediates the interactions between actin and myosin.

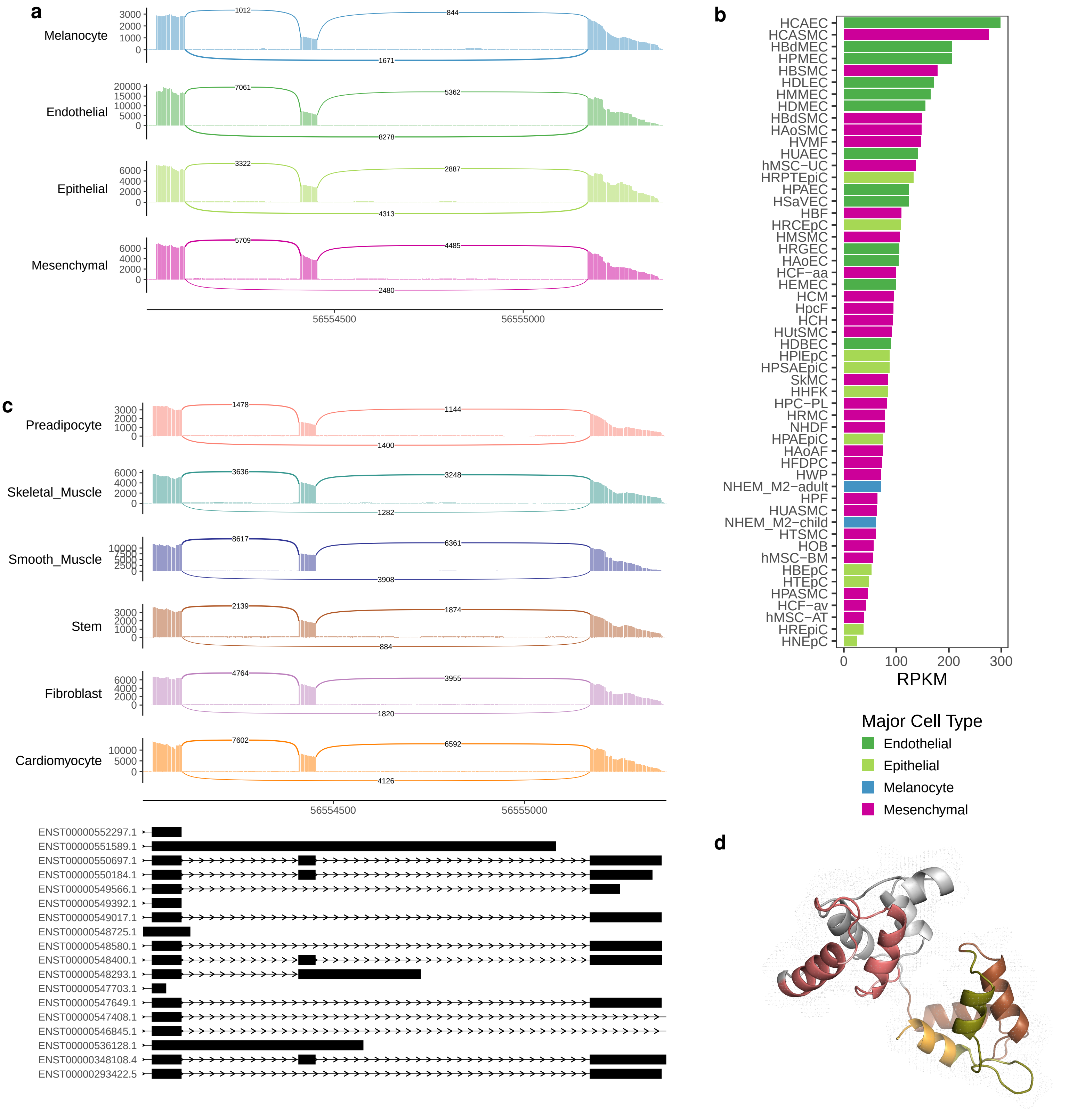

**Fig. S21**. **a)** Contribution of TSS expression (var.bwGp) and contribution of TSS alternative usage (var.bwGp.ge) to variation in TSS abundances across cell types. Genes with significant changes in TSS usage across cell types are in blue. **b)** Expression signal for the *S100A16* gene, which shows the preferential usage of the proximal TSS in endothelial cells, compared to preferential use of the distal TSS in cells from the other types. The individual signal tracks are shown for endothelial cells, whereas overlaid tracks are shown for the other major cell types. The signal is scaled to the maximum of the track height to show the relative difference in TSS usage.

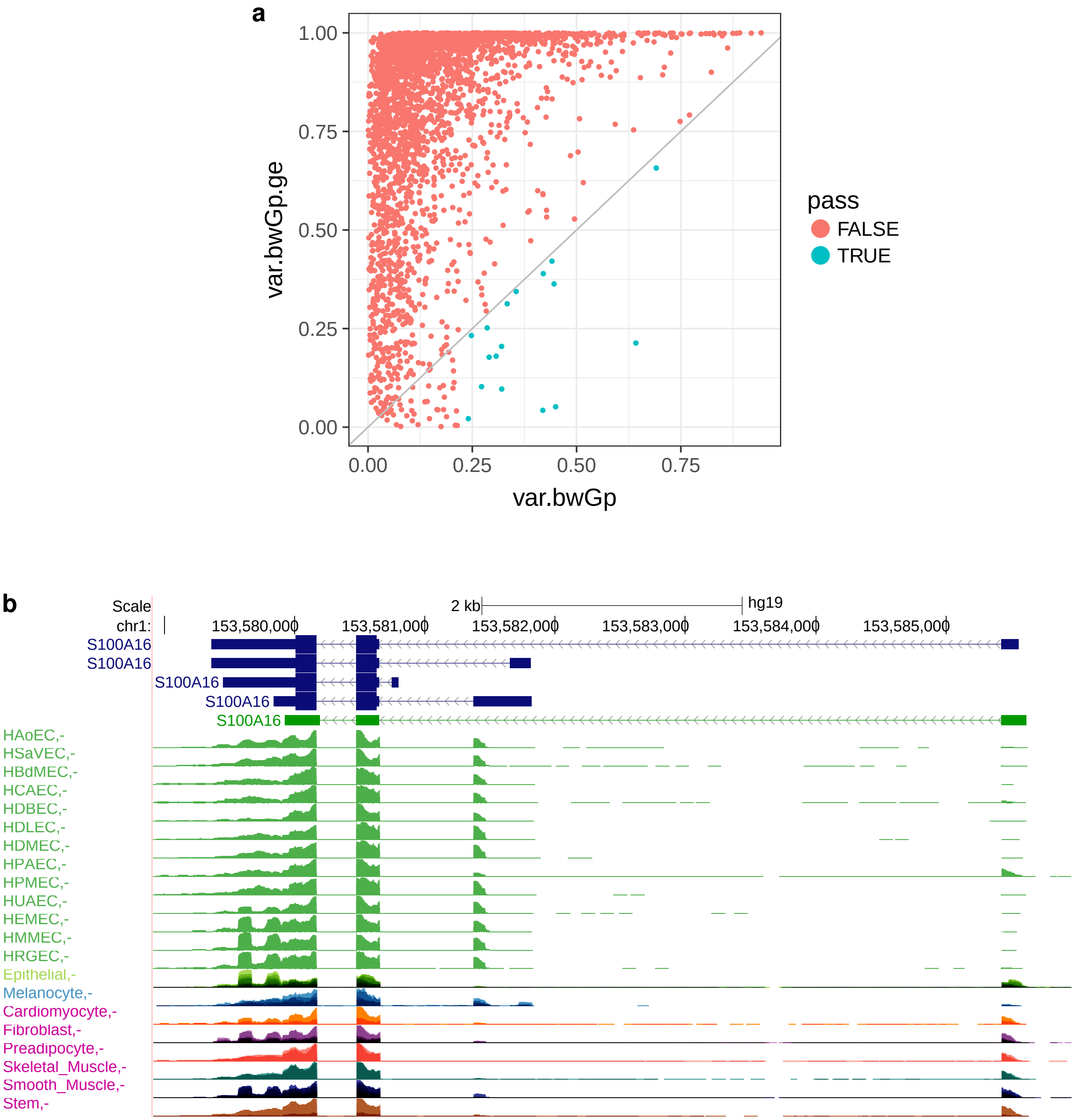

**Fig. S22.** **a)** Evolutionary conservation of cell type specific genes. Percentage of cell type specific genes (dots) with detected 1 to 1 orthologs in worm (*Caenorhabditis Elegans*) and fly (*Drosophila Melanogaster*) in comparison to the distribution (box-plots) of this metric across 1000 random samplings of the protein coding gene set with size equal to the average number of cell-type specific genes (see Fig. 3c). This plot confirms that the size differences between the gene sets do not have an impact on the result. **b)** Heatmap showing the $-\log_{10}(P)$ for the one-tailed hypergeometric tests performed to assess the significance for the enrichment of orthologous genes from each major cell type group when compared to protein coding genes overall, across vertebrates (see Fig. 3d). Significant comparisons at 5% FDR (Benjamini-Hochberg) are marked with an asterisk. **c)** Heatmap showing the $-\log_{10}(P)$ for the one-tailed Wilcoxon Rank-Sum tests performed to assess the significance for the higher conservation of the expression of major cell type specific genes with respect to that of protein coding genes overall, across vertebrates (see Fig. 3e). Significant comparisons at 5% FDR (Benjamini-Hochberg) are marked with an asterisk.

**a**

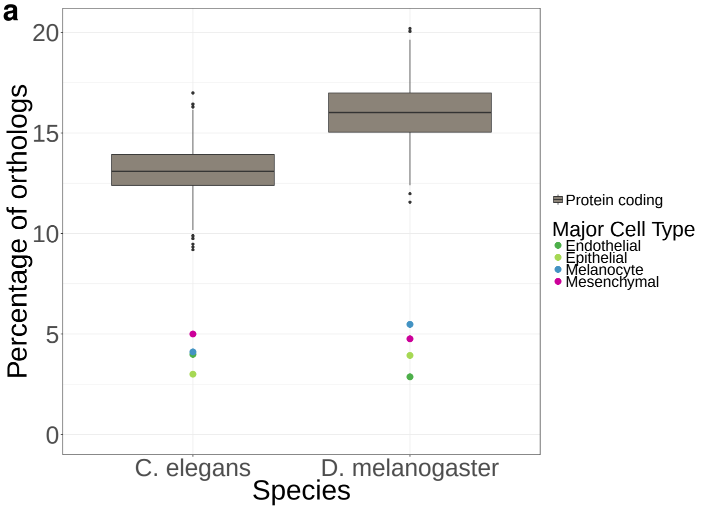


**b**

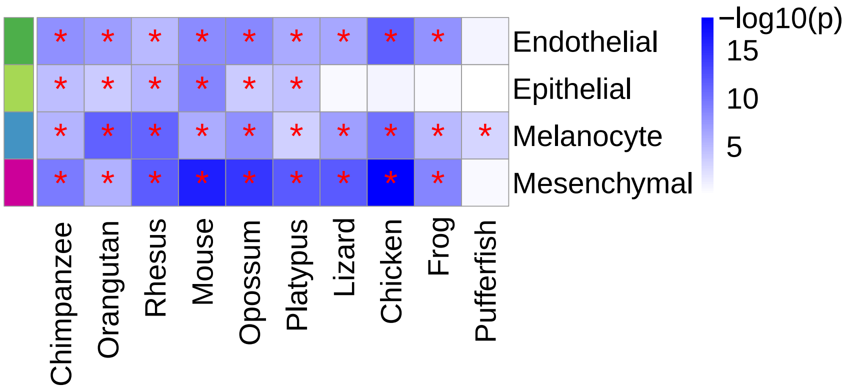


**c**

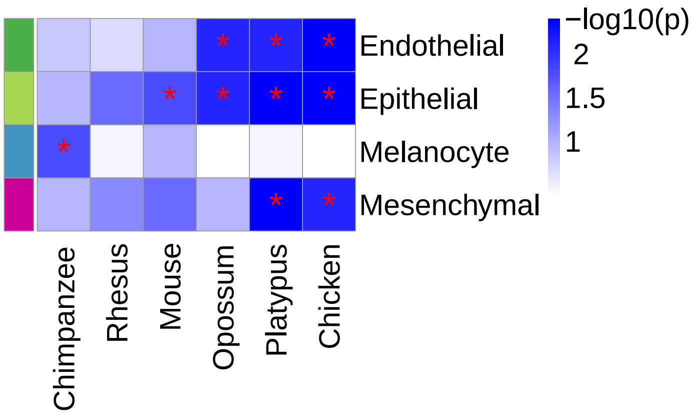

**Fig. S23**. Pearson's correlation coefficient between gene expression in each human organ, except Testis, and the corresponding one in each of the other species. The correlation is computed across all the genes for each major cell type separately. The plot shows that the effect in Fig. 3e is not driven by testis.

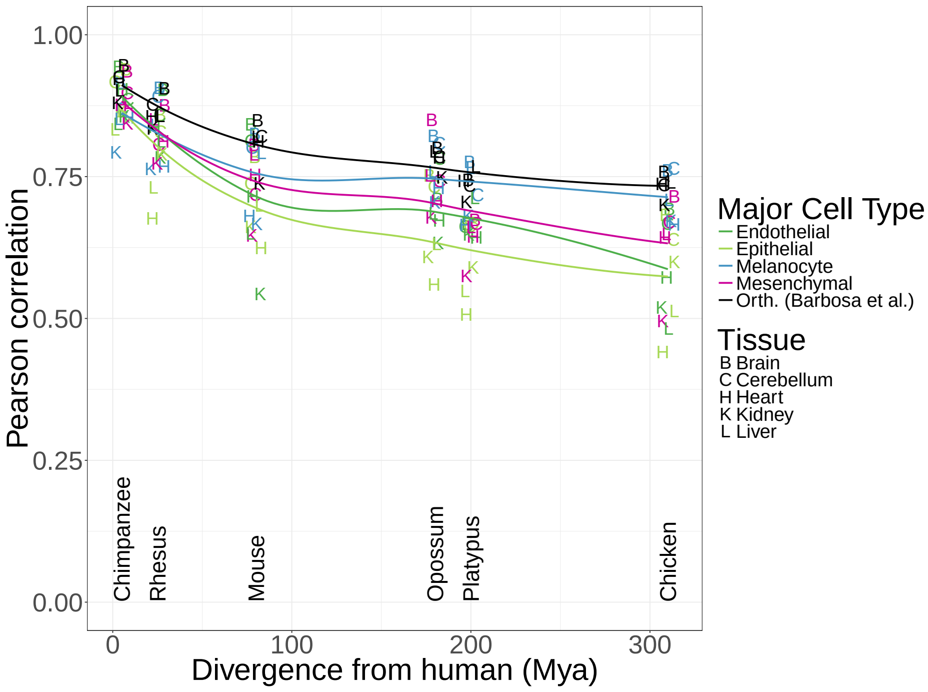

**Fig. S24.** **a)** Histology image transformations in order to estimate the proportion of fat. The top left panel shows the original image, top right shows the mask for the regions of interest, bottom left panel shows the transformed image that results from assigning to each pixel one of the two most dominant colors inside the masked region. The bottom right panel is the same as the left one but with a black background, generated for quality evaluation purposes. **b)** Histogram of the estimated fat proportion from images of subcutaneous adipose tissue from GTEx.

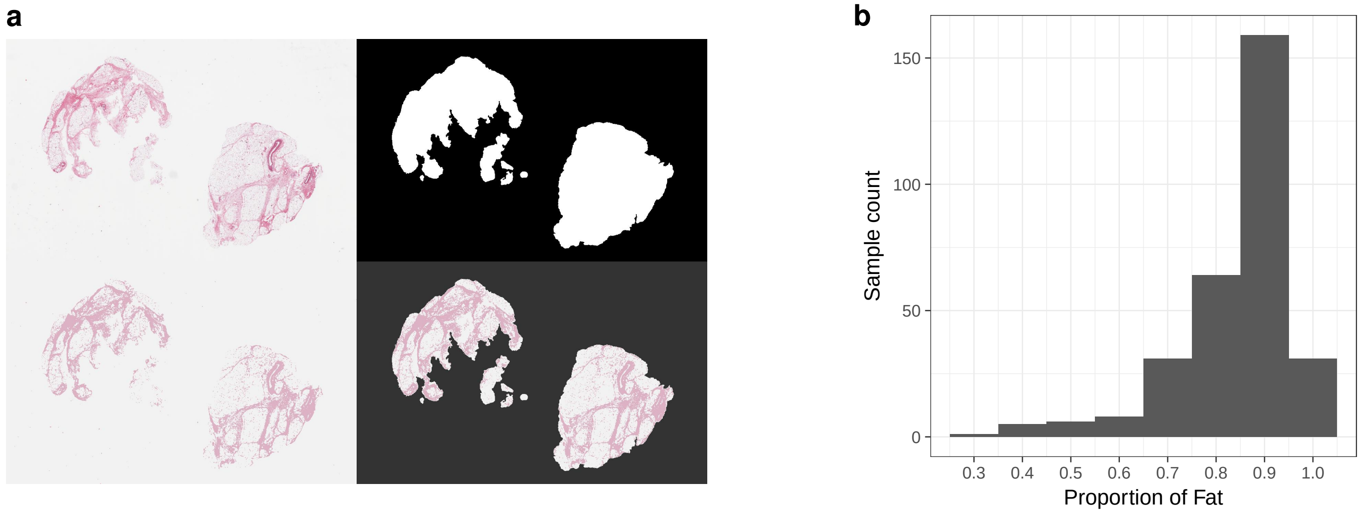

**Fig. S25. a-c)** Pseudotemporal sorting of single cells along induced human myoblast differentiation based on gene expression[25]. The cells are colored based on their discrete differentiation state (a), the calculated pseudotime (b) or the Pearson's correlation coefficient between each single cell and the average expression within ENCODE major cell types and GTEx skeletal muscle tissues (c). The correlations between each single cell and the major cell types and GTEx skeletal muscle tissues are plotted in different colors and overlaid with transparency, so that only the color of the highly correlated major cell types or tissues are evident. Increasing differentiation is observed from right to left, and more differentiated single cells show higher correlation with skeletal muscle specific genes from GTEx, while less differentiated single cells are more correlated with mesenchymal gene expression. **d)** tSNE on principal components of the joined expression data for differentiating myoblast single cells and ENCODE primary cells. Single cell myoblasts are clustered with mesenchymal cells, confirming their similarity in transcriptional programs. The two primary cells misassigned in the clustering of Fig. 1b, renal mesangial cells and lung epithelial cells are not employed here.

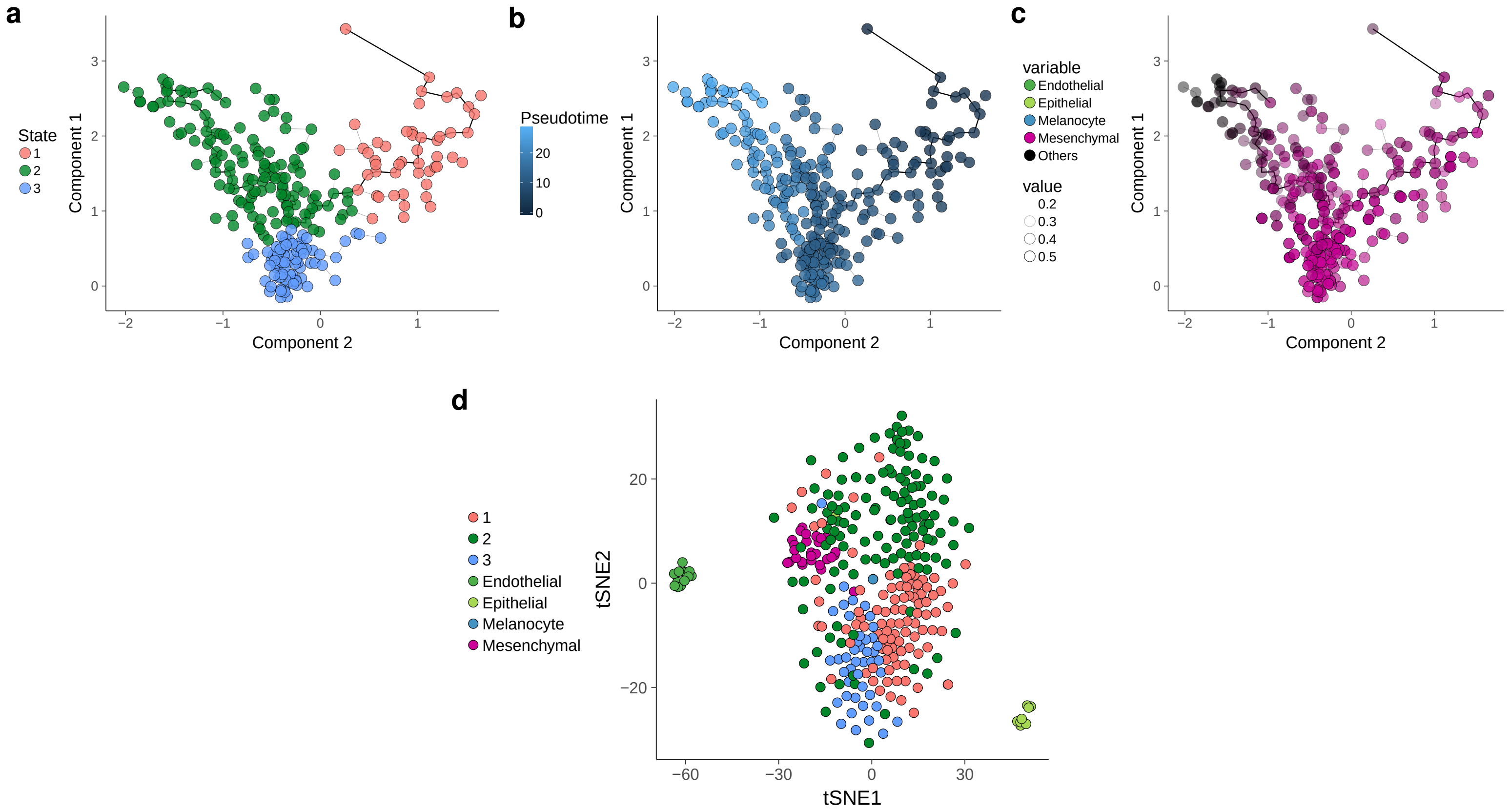

**Fig. S26**. **a)** ROC curve and AUC of the mucosa/muscularis classification of the Stomach samples either in mucosa and muscularis (red), muscularis only (blue) or mucosa only (green). To compute the specificity and sensibility values, each class is tested against the others. **b)** Confusion matrix and statistics of the Stomach samples classification. **c)** Estimated xCell enrichments of epithelial and mesenchymal cells in colon.

**a**

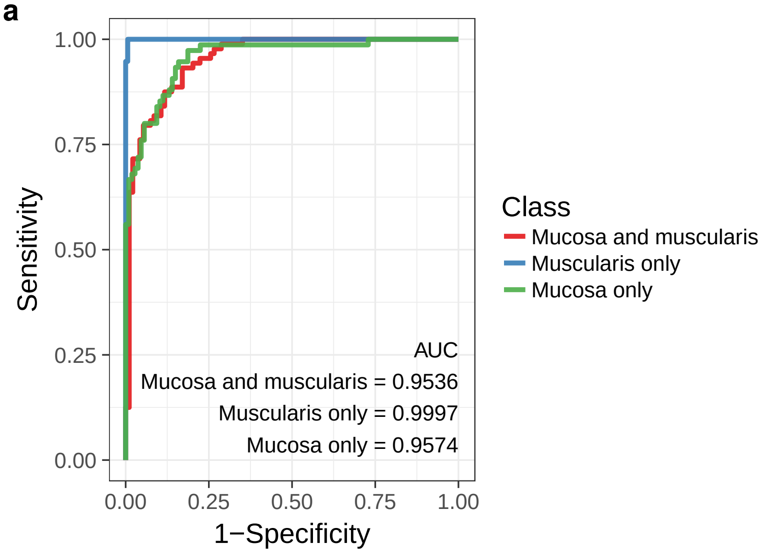


**b**

| | Reference | | |
|---|---|---|---|
| Prediction | Mucosa and muscularis | Muscularis only | Mucosa only |
| Mucosa and muscularis | 77 | 1 | 11 |
| Muscularis only | 0 | 18 | 0 |
| Mucosa only | 11 | 0 | 64 |

| | Mucosa and muscularis | Muscularis only | Mucosa only |
|---|---|---|---|
| Sensitivity | 87.50% | 94.74% | 85.33% |
| Specificity | 87.23% | 100% | 89.72% |

**c**

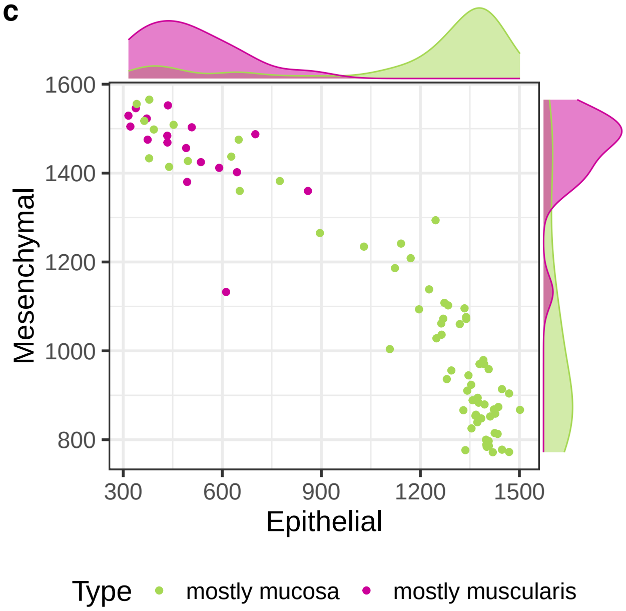

**Fig. S27.** Dimensionality reduction (tSNE and UMAP) of the 8,527 GTEx samples, using the five major cell type enrichment scores computed by xCell.

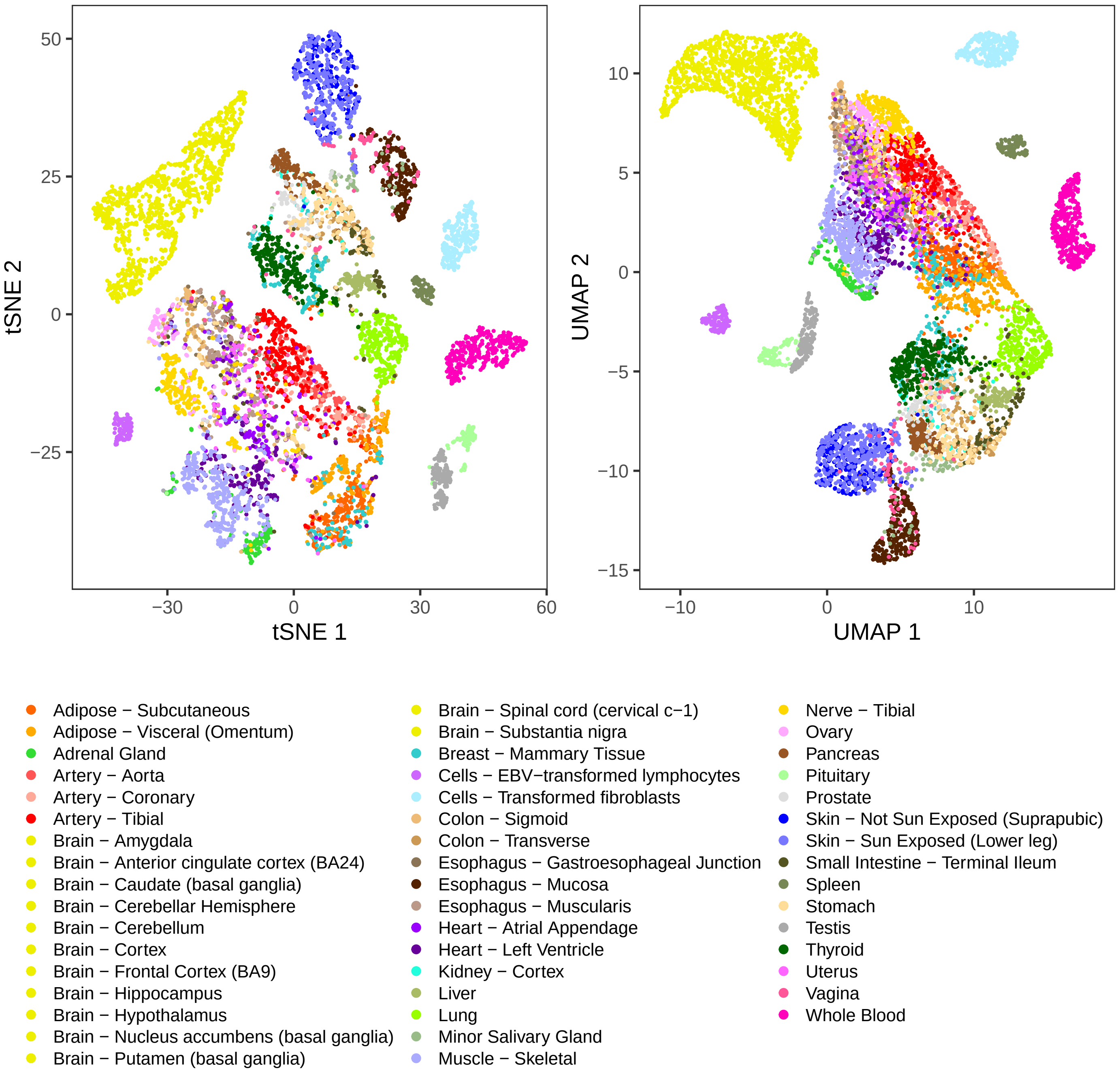

**Fig. S28**. Network modularity of GTEx samples. The network of GTEx samples is created for increasing network densities, which depend on different thresholds of pairwise Pearson's correlation coefficients. Network densities are measured as the percentage of edges over the total number of possible edges. Correlation coefficients are computed over expression values across all genes (RPKM) or over the estimated enrichment scores (xCell) for the five major cell types: epithelial, endothelial, mesenchymal, neural, and blood. The network modularity is computed in both cases at the GTEx organ subregion level. The clustering based on the cellular enrichments (five values) recapitulates tissue type as precisely as the clustering based in gene expression (46,817 values). Indeed, modularity is nearly identical when grouping the samples by gene expression or by cellular enrichments for any threshold on the correlation defining the network edges.

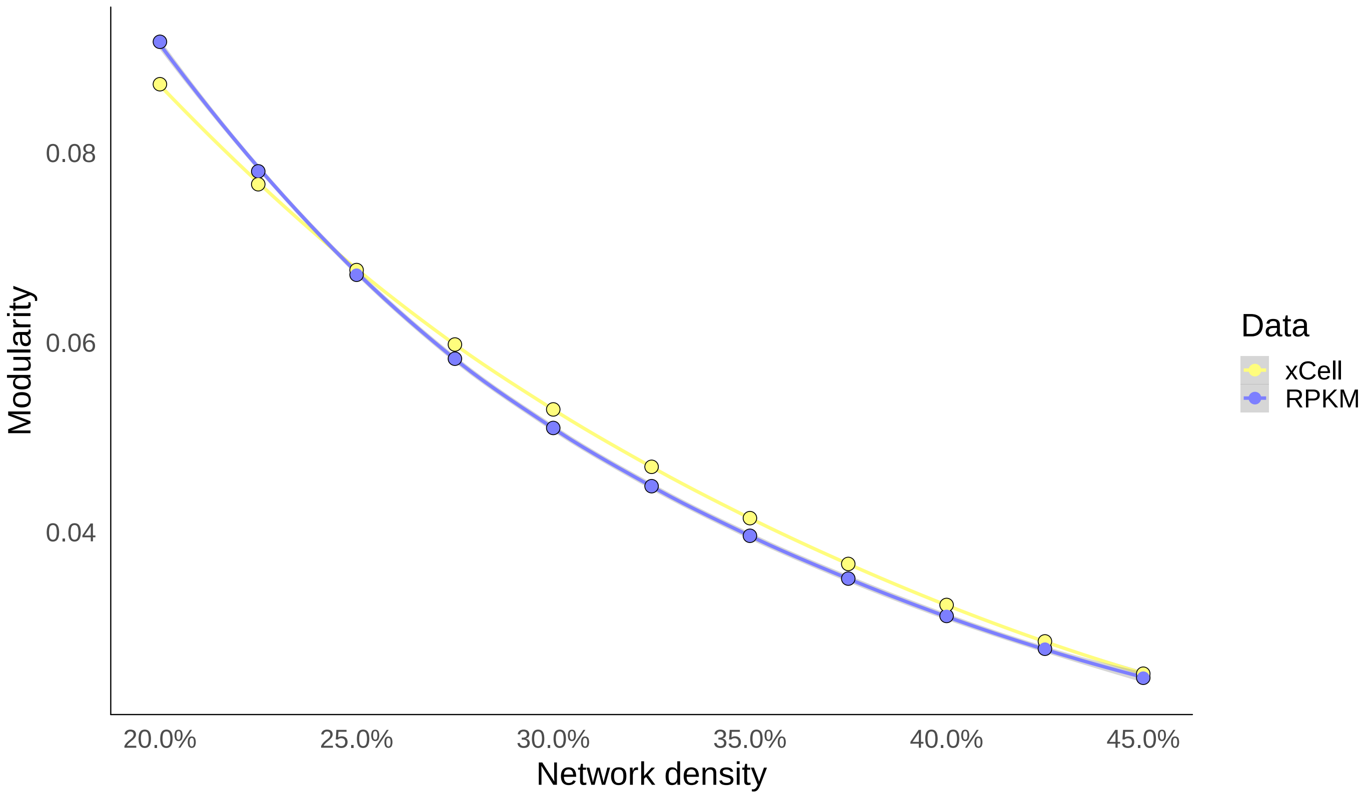

**Fig. S29.** Relationship between donor age and incidence of tibial artery atherosclerosis (**a**) and muscle atrophy (**b**).

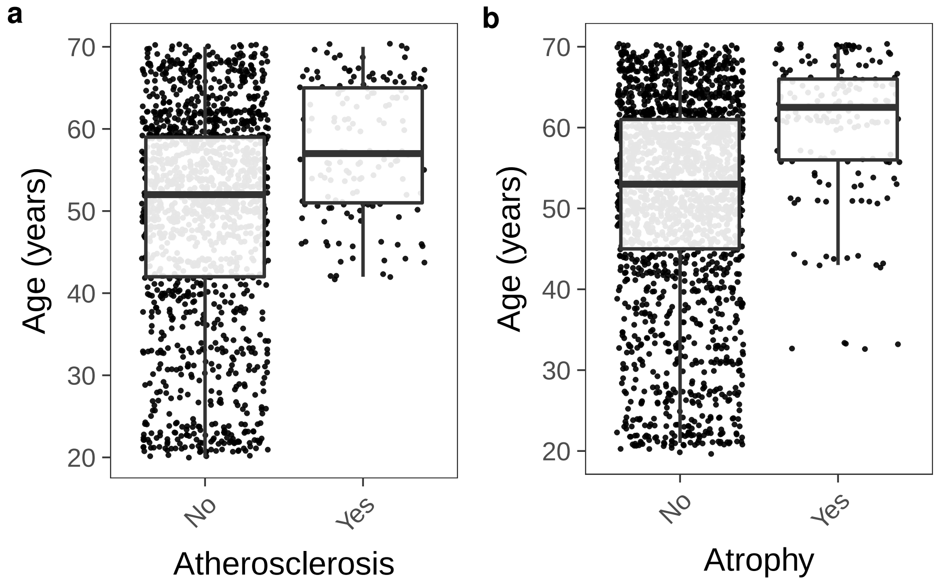

**Fig. S30**. Differences in estimated enrichments by xCell between normal liver and liver affected with congestion in the GTEx pathology reports (**a**), between reduced and active spermatogenesis in testis (**b**), between lung tissue annotated with pneumonia and normal lung (**c**). n is the number of affected cases in each histological phenotype. **d)** Relationship between estimated enrichments and respiratory cause of death in lung (13 of 320 donors). **e)** Relationship between donor age and incidence of respiratory cause of death. **f**) Differences between ovary sections from younger (pre-menopausal) and older (postmenopausal) women. Each panel shows the FDR for Wilcoxon's test. The displayed phenotypes have at least one major cell type with FDR $<$ 0.01.

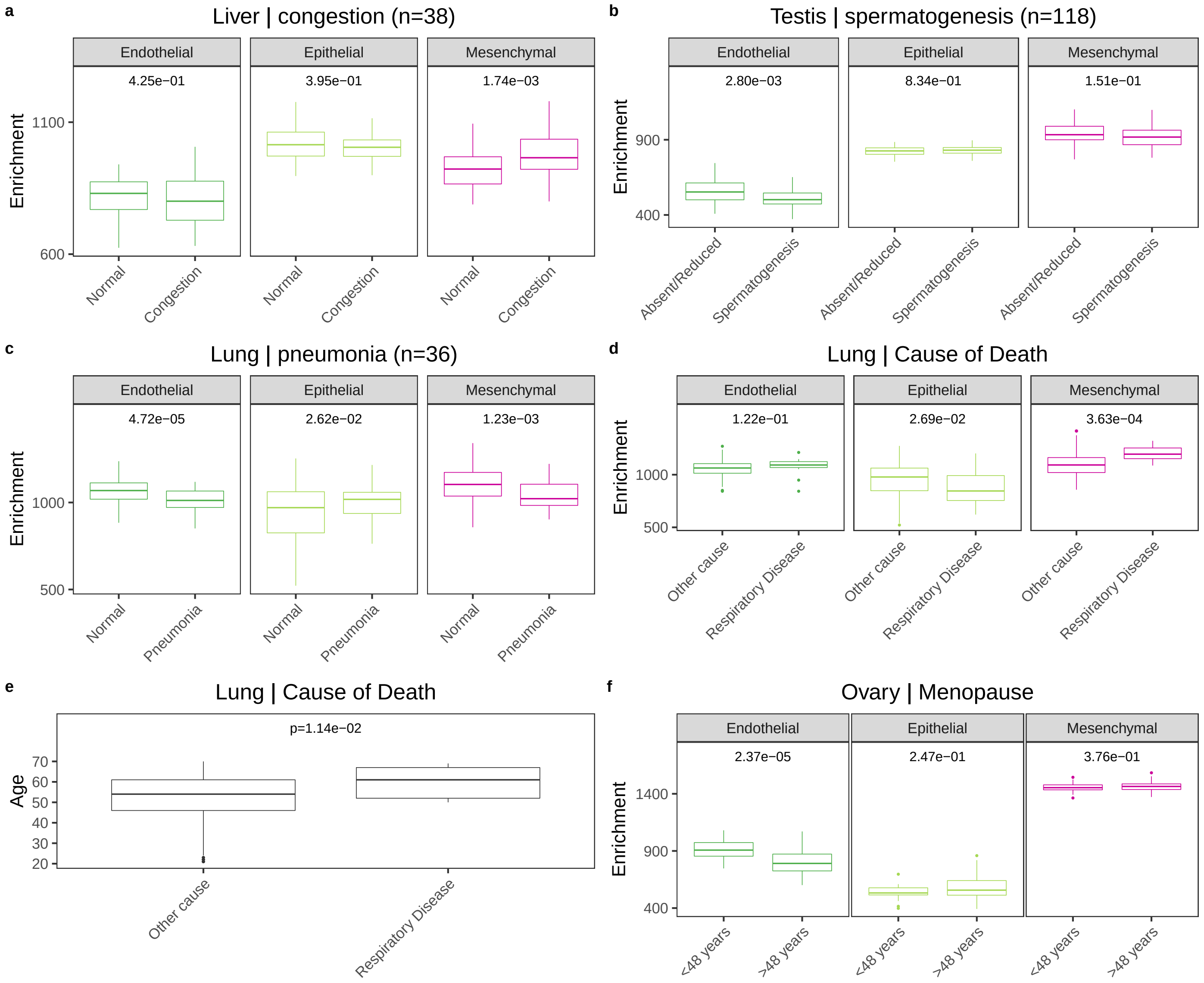

**Fig. S31.** Estimated cellular proportion by xCell for 1,963 cancer and normal samples (when available) from the Cancer Genome Atlas Pan-Cancer analysis project [34].

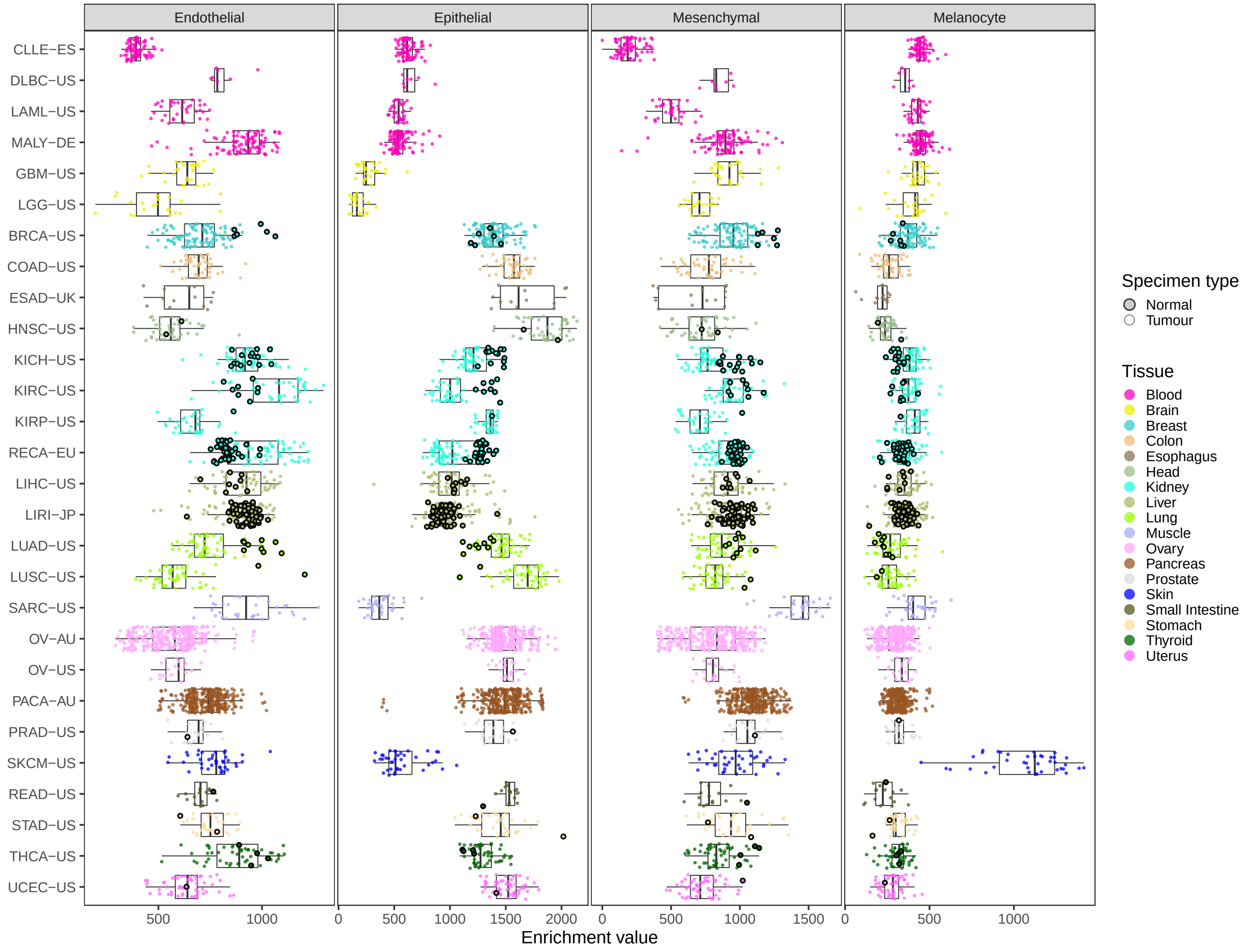


| DCC Project Code | Project Name | Country |
|---|---|---|
| CLLE-ES | Chronic Lymphocytic Leukemia - ES | Spain |
| DLBC-US | Lymphoid Neoplasm Diffuse Large B-cell Lymphoma - TCGA, US | US |
| LAML-US | Acute Myeloid Leukemia - TCGA, US | US |
| MALY-DE | Malignant Lymphoma - DE | Germany |
| GBM-US | Brain Glioblastoma Multiforme - TCGA, US | US |
| LGG-US | Brain Lower Grade Glioma - TCGA, US | US |
| BRCA-US | Breast Cancer - TCGA, US | US |
| COAD-US | Colon Adenocarcinoma - TCGA, US | US |
| ESAD-UK | Esophageal Adenocarcinoma - UK | United Kingdom |
| HNSC-US | Head and Neck Squamous Cell Carcinoma - TCGA, US | US |
| KICH-US | Kidney Chromophobe - TCGA, US | US |
| KIRC-US | Kidney Renal Clear Cell Carcinoma - TCGA, US | US |
| KIRP-US | Kidney Renal Papillary Cell Carcinoma - TCGA, US | US |
| RECA-EU | Renal Cell Cancer - EU/FR | European Union/France |

| DCC Project Code | Project Name | Country |
|---|---|---|
| LIHC-US | Liver Hepatocellular carcinoma - TCGA, US | US |
| LIRI-JP | Liver Cancer - RIKEN, JP | Japan |
| LUAD-US | Lung Adenocarcinoma - TCGA, US | US |
| LUSC-US | Lung Squamous Cell Carcinoma - TCGA, US | US |
| SARC-US | Sarcoma - TCGA, US | US |
| OV-AU | Ovarian Cancer - AU | Australia |
| OV-US | Ovarian Serous Cystadenocarcinoma - TCGA, US | US |
| PACA-AU | Pancreatic Cancer Endocrine Neoplasms- AU | Australia |
| PRAD-US | Prostate Adenocarcinoma - TCGA, US | US |
| SKCM-US | Skin Cutaneous melanoma - TCGA, US | US |
| READ-US | Rectum Adenocarcinoma - TCGA, US | US |
| STAD-US | Gastric Adenocarcinoma - TCGA, US | US |
| THCA-US | Head and Neck Thyroid Carcinoma - TCGA, US | US |
| UCEC-US | Uterine Corpus Endometrial Carcinoma- TCGA, US | US |
